# Optimisation of pembrolizumab therapy for de novo metastatic MSI-H/dMMR colorectal cancer using data-driven delay integro-differential equations

**DOI:** 10.1101/2025.05.23.655752

**Authors:** Georgio Hawi, Peter S. Kim, Peter P. Lee

## Abstract

Colorectal cancer (CRC), the third most commonly diagnosed cancer worldwide, presents a growing public health concern, with 20% of new diagnoses involving de novo metastatic disease and up to 80% of these patients presenting with unresectable metastatic lesions. Microsatellite instability-high (MSI-H) CRC and deficient mismatch repair (dMMR) CRC constitute 15% of all CRC, and 4% of metastatic CRC, and, while less responsive to conventional chemotherapy, exhibit notable sensitivity to immunotherapy, especially programmed cell death protein 1 (PD-1) checkpoint inhibitors such as pembrolizumab. Despite this, there is a significant need to optimise immunotherapeutic regimens to maximise clinical efficacy and patient quality of life whilst minimising financial burden. In this work, we adapt our mechanistic model for locally advanced MSI-H/dMMR CRC to de novo metastatic MSI-H/dMMR CRC (dnmMCRC), deriving model parameters from pharmacokinetic, bioanalytical, and radiographic studies, as well as bulk RNA-sequencing data deconvolution from the TCGA COADREAD and GSE26571 datasets. We finally optimised treatment with pembrolizumab to balance efficacy, efficiency, and toxicity in dnmMCRC, comparing against currently FDA-approved regimens, analysing factors influencing treatment success and comparing immune dynamics to those in locally advanced disease.

## 1 Introduction

Colorectal cancer (CRC) is the third most common cancer worldwide, with more than 1.85 million cases and 850,000 deaths annually [1]. Of new CRC diagnoses, 20% of patients present with de novo metastatic disease, with an estimated 75%–90% of these patients presenting with unresectable metastatic lesions [2]. Furthermore, an additional 25% of patients who initially have localised disease eventually develop metastases [1]. However, since many people will not experience symptoms in the early stages of CRC, diagnoses often occur at a later stage when the disease is more advanced, where treatment is less effective, and survival rates are significantly lower [3]. In the United States, the 5-year survival rates for stage IIIA, stage IIIB, and Stage IIIC colon cancer are 90%, 72%, and 53%, respectively, whilst stage IV CRC has a 5-year survival of only 12% [4].

Whilst many systemic therapies are available for advanced CRC, chemotherapy has been the main treatment approach, with mainstay chemotherapy regimens such as FOLFOX (folinic acid, 5-fluorouracil, oxaliplatin) and FOLFIRI (folinic acid, 5-fluorouracil, irinotecan) becoming integral to the treatment of advanced CRC [5]. However, patients with the hypermutant microsatellite instability-high (MSI-H) phenotype who have reached metastasis are less responsive to conventional chemotherapy and have a poorer prognosis compared to patients with microsatellite stable (MSS) CRC [6]. In particular, we note that in CRC, MSI-H and deficient mismatch repair (dMMR) tumours are equivalent [7], and we denote these tumours as MSI-H/dMMR for the remainder of this work. Approximately 20% of stage II, 12% of stage III, and 4% of stage IV CRC tumours are diagnosed as MSI-H/dMMR [8–10], with approximately 80% of sporadic MSI-H/dMMR CRC caused by MLH1 promoter hypermethylation [11]. This leads to a highly increased mutational rate, with MSI-H/dMMR CRC tumours having 10–100 times more somatic mutations compared to microsatellite stable (MSS) CRC tumours [12], resulting in increased tumour mutation burden (TMB) and neoantigen load, and immunogenic tumour microenvironment (TME) with dense immune cell infiltration [13, 14]. This immunogenicity results in patients with MSI-H/dMMR CRC having a good prognosis for immunotherapy treatment, in particular to immune checkpoint inhibitors (ICIs) [15].

Immune checkpoints, such as programmed cell death-1 (PD-1), cytotoxic T-lymphocyte-associated antigen 4 (CTLA-4), and lymphocyte-activation gene 3 (LAG-3), normally downregulate immune responses after antigen activation [16]. PD-1, a cell membrane receptor that is expressed on a variety of cell types, including activated T cells, activated B cells and monocytes, has been extensively researched in the context of cancer such as MSI-H/dMMR CRC [17, 18]. When PD-1 interacts with its ligands (PD-L1 and PD-L2), effector T cell activity is inhibited, resulting in the downregulation of pro-inflammatory cytokine secretion and the upregulation of immunosuppressive regulatory T cells (Tregs) [19, 20]. Cancers can exploit this by expressing PD-L1 themselves, evading immunosurveillance, and impairing the proliferation and activity of cytotoxic T lymphocytes (CTLs) [21]. Blockade of PD-1/PD-L1 complex formation reinvigorates effector T cell activity, resulting in enhanced antitumour immunity and responses, leading to improved clinical outcomes in cancer patients [22, 23].

The KEYNOTE-177 phase III trial, NCT02563002, aimed to evaluate the efficacy of first-line pembrolizumab, an anti-PD-1 antibody, in metastatic MSI-H/dMMR CRC (mMCRC) [11]. In the trial, 307 treatment-naive mMCRC patients were randomly assigned to receive pembrolizumab at a dose of 200 mg every 3 weeks or 5fluorouracilbased chemotherapy every 2 weeks. A partial or complete response was observed in 43.8% of patients allocated to pembrolizumab therapy, compared with 33.1% of patients participating in 5-fluorouracil-based therapy. Furthermore, among patients who responded, 83% in the pembrolizumab group maintained response at 24 months, compared with 35% of patients receiving chemotherapy. These results motivated the FDA to approve pembrolizumab for the first-line treatment of unresectable or mMCRC on June 29, 2020 [24].

An important question to consider is the appropriate dosing and spacing of ICI therapies to balance tumour reduction with factors such as monetary cost, toxicity, and side effects [25, 26]. Mathematical models provide a powerful framework for optimising treatment regimens, whilst avoiding the significant time and financial costs associated with human clinical trials. There are numerous immunobiological models of CRC, such as those from [27–30], and ICI therapy has been modelled extensively in other cancers, including those in [31–35]. Nonetheless, there are a multitude of limitations and drawbacks to these models of CRC and ICI therapy, as detailed in [36]. Additionally, to date, there are no pre-existing mathematical models in the literature for ICI therapy in de novo mMCRC (dnmMCRC). Furthermore, to the authors’ knowledge, [36] presents the only immunobiological model of ICI therapy in CRC, and also addresses these drawbacks, focusing on the modelling and optimisation of neoadjuvant pembrolizumab therapy in locally advanced MSI-H/dMMR CRC (laMCRC). In this work, we adapt this model to dnmMCRC and use this to optimise pembrolizumab therapy as well as compare immune dynamics to those in laMCRC.

It is prudent for us to briefly outline, as repeated in [36], the functions and processes of some immune cells in the TME since their interaction with cancer cells directly or through chemokine/cytokine signalling significantly influences the efficacy of therapeutic regimens [37]. T cell activation occurs in the lymph node and occurs through T cell receptor (TCR) recognition of cancer antigen presented by major histocompatibility complex (MHC) class I molecules, in the case of CD8+ T cells, and MHC class II molecules, in the case of CD4+ T cells, expressed on the surfaces of mature DCs [38]. CTLs recognise cancer cells through TCR detection of peptide major histocompatibility complexes (pMHCs) on cancer cell surfaces via MHC class I [39]. CD8+ cells, as well as NK cells, are amongst the most cytotoxic and important cells in cancer cell lysis [40], in addition to secreting pro-inflammatory cytokines such as interleukin-2 (IL-2), interferon-gamma (IFN-γ), and tumour necrosis factor (TNF) [41]. These are also secreted by CD4+ T helper 1 (Th1) cells and are an important part of cell-mediated immunity, allowing for neutrophil chemotaxis and macrophage activation [42]. Furthermore, we must also consider Tregs, which are vital in immune tissue homeostasis since they are able to suppress the synthesis of pro-inflammatory cytokines and control intestinal inflammatory processes [43]. This is done in a variety of ways, including the production of immunomodulatory and immunosuppressive cytokines such as TGF-β, interleukin-10 (IL-10), and interleukin-35 (IL-35) [44, 45]. We note that naive CD4+ T cells can differentiate towards multiple additional phenotypes such as Th2, Th9, Th22, Tfh and Th17 cells, each involved in the pathogenesis of cancer [46, 47].

Also of importance in CRC are macrophages, which, like T cells, are able to produce pro-inflammatory and anti-inflammatory cytokines [48]. Naive macrophages, denoted M0 macrophages, can differentiate into two main phenotypes: classically activated M1 macrophages and alternatively activated M2 macrophages. These names were given since M1 macrophages promote Th1 cell responses, and M2 macrophages promote Th2 responses, with Th1-associated cytokines downregulating M2 activity, and vice versa [49]. M1 macrophages contribute to the inflammatory response by activating endothelial cells, promoting the induction of nitric oxide synthase, and producing large amounts of pro-inflammatory cytokines such as TNF, interleukin-1β (IL-1β), and IL-12 [50]. On the other hand, M2 macrophages are responsible for wound healing and the resolution of inflammation through phagocytosing apoptotic cells and releasing anti-inflammatory mediators such as IL-10, interleukin-13α1 (IL-13α1), and CC Motif Chemokine Ligand 17 (CCL17) [51].

It is important to note that the M1/M2 macrophage dichotomy is somewhat of a simplification. Macrophages are highly plastic and have been demonstrated to integrate environmental signals to change their phenotype and physiology [52]. To account for this, in the model, we incorporate macrophage polarisation and repolarisation between its anti-tumour and immunosuppressive phenotypes by various cytokines.

## 2 Mathematical Model

### 2.1 Model Variables and Assumptions

The variables and their units in the model are shown in Table 1.

**Table 1:**
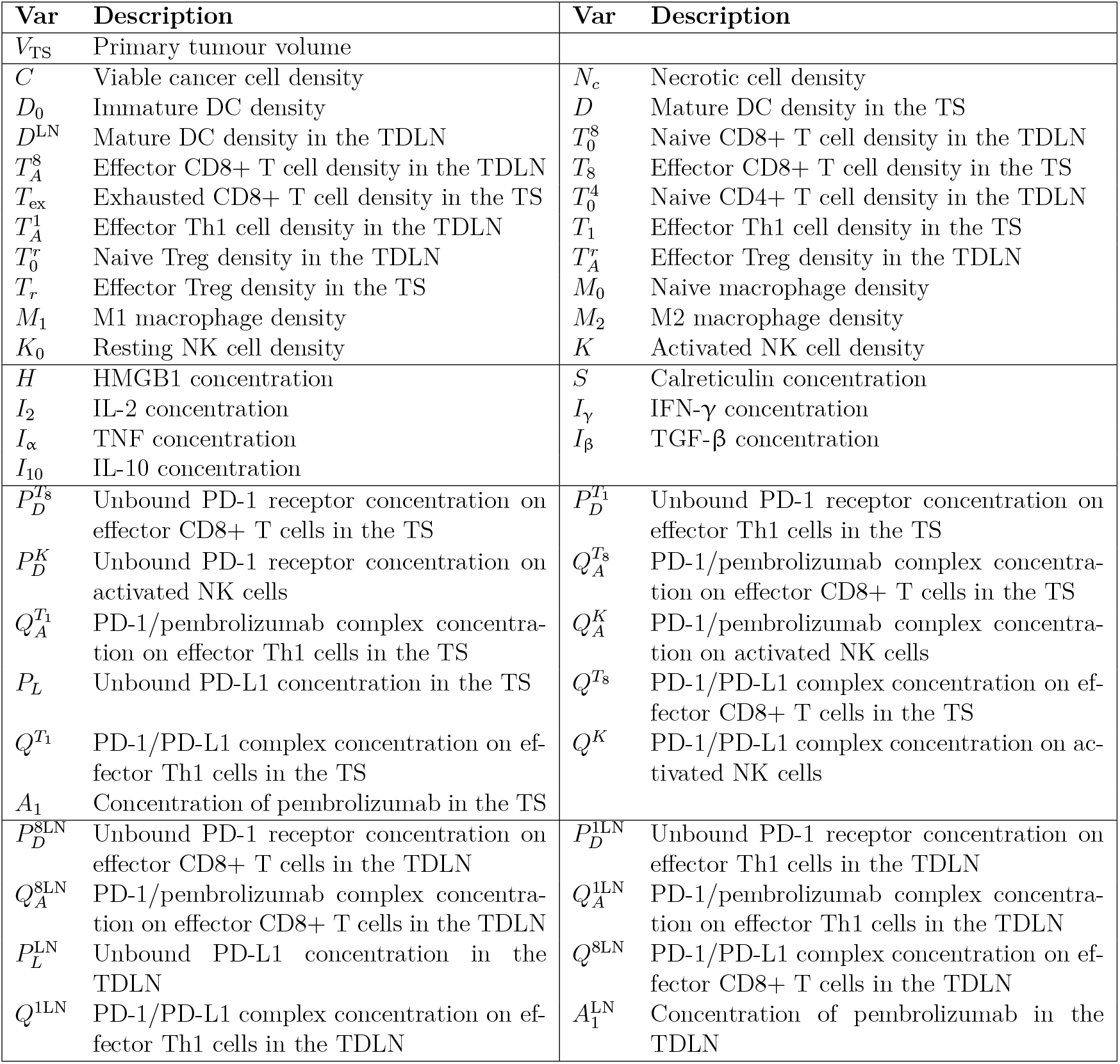
Variables used in the model. Quantities in the top box are in units of cell/cm^3^, quantities in the second box are in units of g/cm^3^, and all other quantities are in units of molec/cm^3^. All quantities pertain to the tumour site unless otherwise specified. TDLN denotes the tumour-draining lymph node, whilst TS denotes the tumour site.

For simplicity, we ignore spatial effects in the model, ignoring the effects of diffusion, advection, and chemotaxis by all species. We assume the system has two compartments: one at the TS, located in the colon or rectum, and one at the tumour-draining lymph node (TDLN). This is a simplification since dnmMCRC typically involves multiple tumour-draining lymph nodes [53]; however, for simplicity, we focus on the sentinel node and refer to it as the TDLN for the purposes of the model. In a similar fashion to nearly all models of de novo metastatic cancer, we primarily focus on the growth of the primary tumour, using it as a proxy to infer the progression of lymph node and distant metastases. We assume that cytokines in the TS are produced only by effector or activated cells and that DAMPs in the TS are only produced by necrotic cancer cells. We assume that all mature DCs considered in the TDLN are cancer-antigen-bearing and that all T cells considered in the TS are primed with cancer antigens. Furthermore, we assume that all activated T cells considered in the TDLN are activated with cancer antigens and that T cell proliferation/division follows a deterministic program. We ignore CD4+ and CD8+ memory T cells and assume that naive CD4+ T cells differentiate immediately upon activation. We also assume that all Tregs in the TS are natural Tregs (nTregs), ignoring induced Tregs (iTregs). We assume, for simplicity, that activated macrophages polarise into the M1/M2 dichotomy. We also assume that the duration of pembrolizumab infusion is negligible compared to the timescale of the model. Therefore, we treat their infusions as an intravenous bolus so that drug absorption occurs immediately after infusion. Finally, we assume a constant solution history, where the history for each species is set to its respective initial condition.

We assume that all species, *X*_*i*_, degrade/die at a rate proportional to their concentration, with decay constant 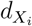. We assume that the rate of activation/polarisation of a species *X*_*i*_ by a species *X*_*j*_ follows the Michaelis-Menten kinetic law 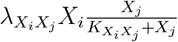, for rate constant 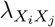, and half-saturation constant 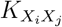. Similarly, we model the rate of inhibition of a species *X*_*i*_ by a species *X*_*j*_ using a term with form 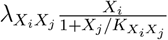 for rate constant 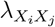, and half-saturation constant 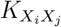. Production of *X*_*i*_ by *X*_*j*_ is modelled using mass-action kinetics unless otherwise specified, so that the rate at which *X*_*i*_ is formed is given by 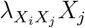 for some positive constant 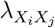. Finally, we assume that the rate of lysis of *X*_*i*_ by *X*_*j*_ follows mass-action kinetics in the case where *X*_*j*_ is a cell and follows Michaelis-Menten kinetics in the case where *X*_*j*_ is a cytokine.

### 2.2 Model Summary

We now outline some of the main processes accounted for in the model, with all processes and equations being explained in Appendix A.

1. Effector CD8+ T cells and NK cells induce apoptosis of cancer cells, with this being inhibited by TGF-β and the PD-1/PD-L1 complex. However, TNF and IFN-γ induce necroptosis of cancer cells, causing them to become necrotic before they are removed.
2. Necrotic cancer cells release DAMPs such as HMGB1 and calreticulin, which stimulate immature DCs to mature.
3. Some mature DCs migrate to the T cell zone of the TDLN and activate naive CD8+ and CD4+ T cells (including Tregs), with CD8+ T cell and Th1 cell activation being inhibited by Tregs and the PD-1/PD-L1 complex.
4. Activated T cells undergo clonal expansion and proliferate rapidly in the TDLN, with CD8+ T cell and Th1 cell proliferation being inhibited by Tregs and the PD-1/PD-L1 complex.
5. T cells that have completed proliferation migrate to the TS and perform effector functions including the production of pro-inflammatory (IL-2, IFN-γ, TNF) and immunosuppressive (TGF-β, IL-10) cytokines. Extended exposure to the cancer antigen can lead CD8+ T cells to become exhausted, however, this exhaustion can be reversed by pembrolizumab.
6. In addition, mature DCs, NK cells and macrophages secrete cytokines that can activate NK cells and polarise and repolarise macrophages into pro-inflammatory and immunosuppressive phenotypes.
7. Pembrolizumab infusion promotes the binding of unbound PD-1 receptors to pembrolizumab, forming the PD-1/pembrolizumab complex instead of the PD-1/PD-L1 complex. This reduces the inhibition of pro-inflammatory CD8+ and Th1 cell activation and proliferation while also reducing the inhibition of cancer cell lysis.

### 2.3 Model Equations

The model equations follow identically from [36], with the exception of *V*_TS_, which is time-dependent as opposed to being constant. For completeness, we provide the mathematical model below, with a full derivation of the model being found in Appendix A.

#### 3.3.1 Equations for Cancer Cells, DAMPs, and DCs

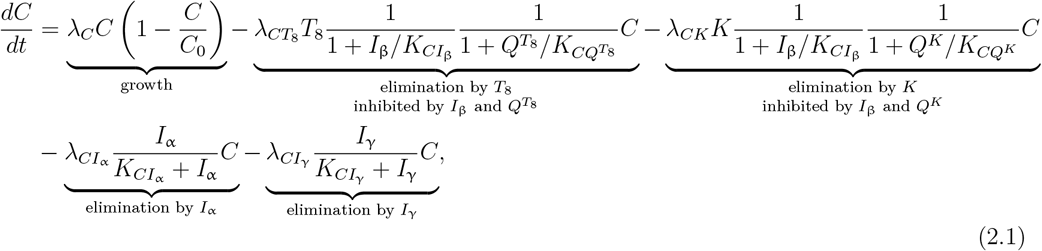

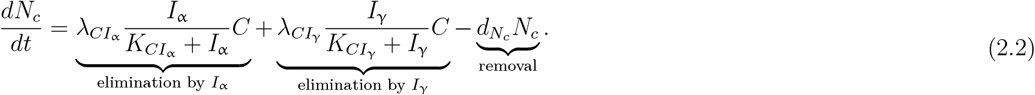

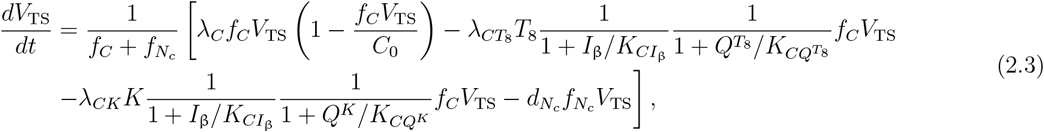

where *C*(*t*) = *f*_*C*_*V*_TS_(*t*) and 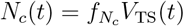.

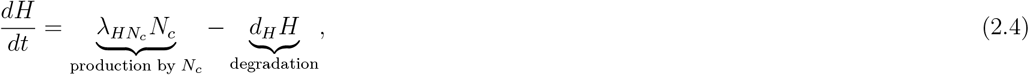

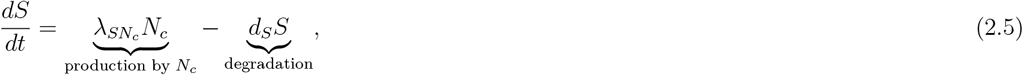

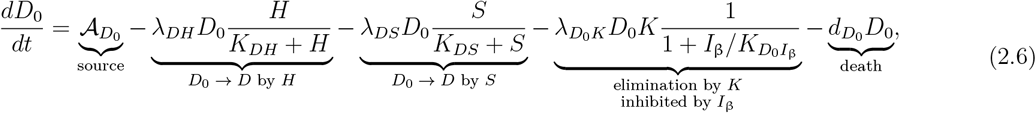

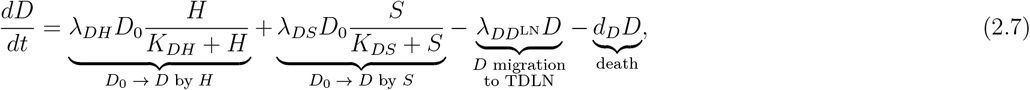

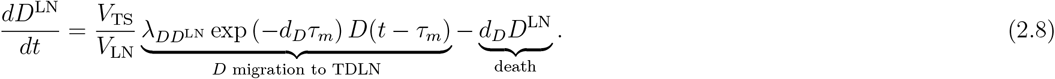

A diagram encompassing the interactions of these components is shown in Figure 1.

**Figure 1:**
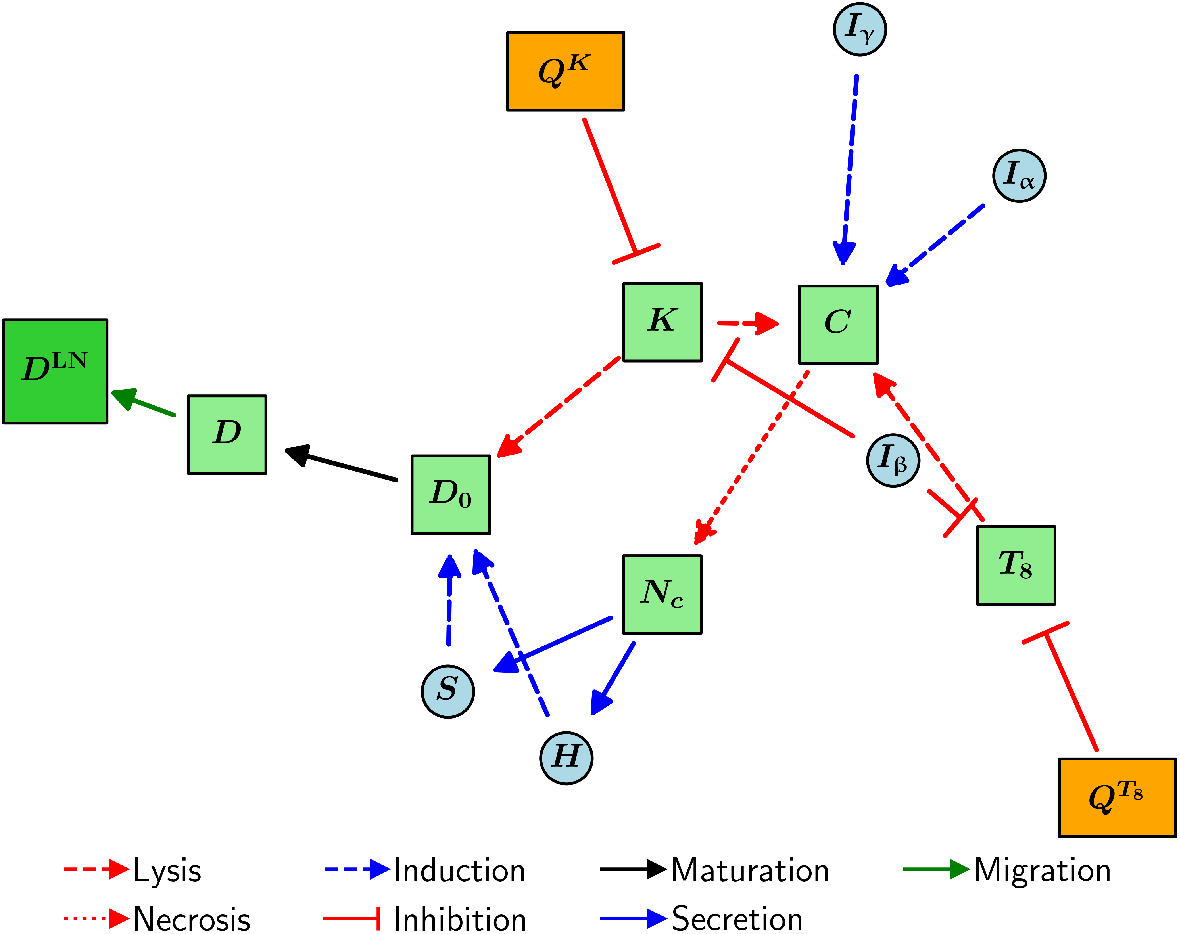
Schematic diagram of the interactions of cancer cells, DAMPs, and DCs in the model.

#### 2.3.2 Equations for T Cells

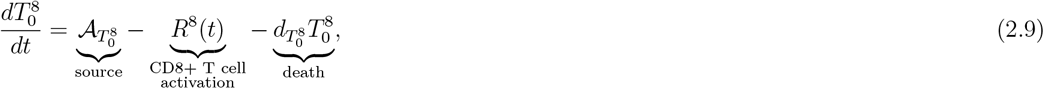

where *R*^8^(*t*) is defined as

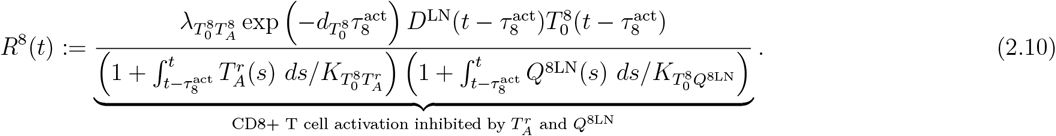

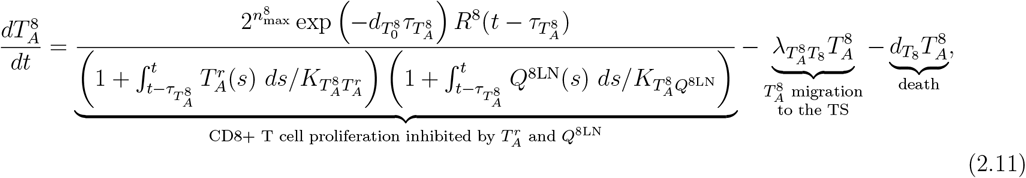

where 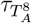 is defined as

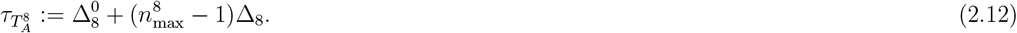

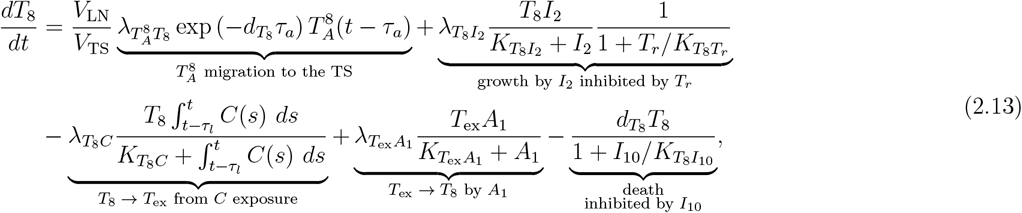

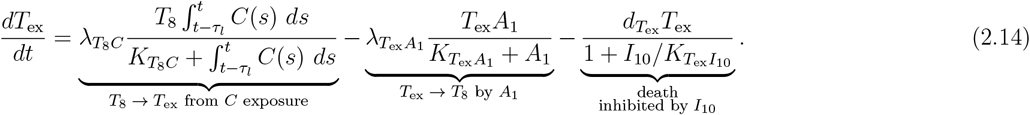

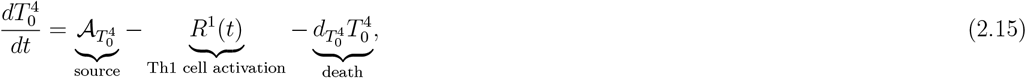

where *R*^1^(*t*) is defined as

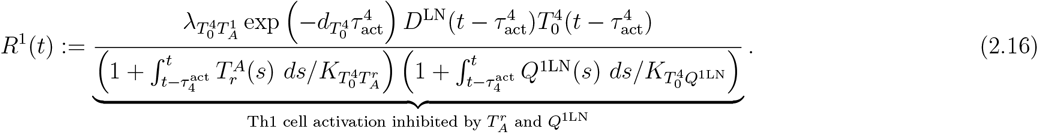

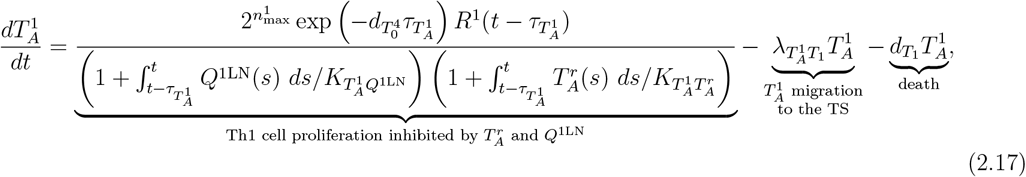

where 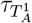 is defined as

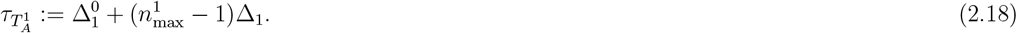

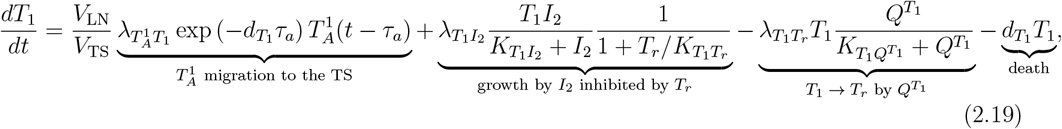

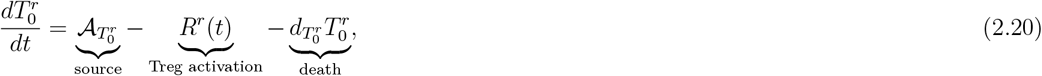

where *R*^*r*^(*t*) is defined as

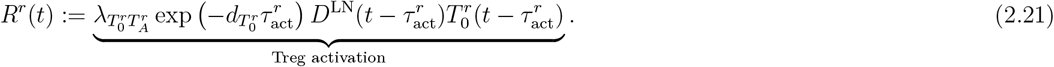

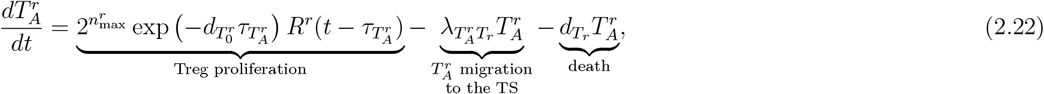

where 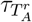 is defined as

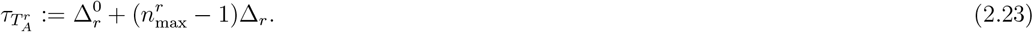

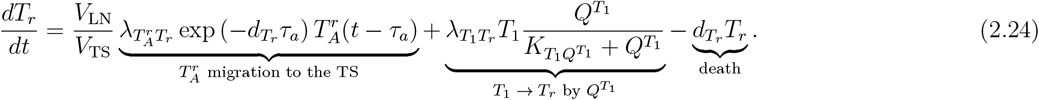

A diagram encompassing the interactions of these components is shown in Figure 2.

**Figure 2:**
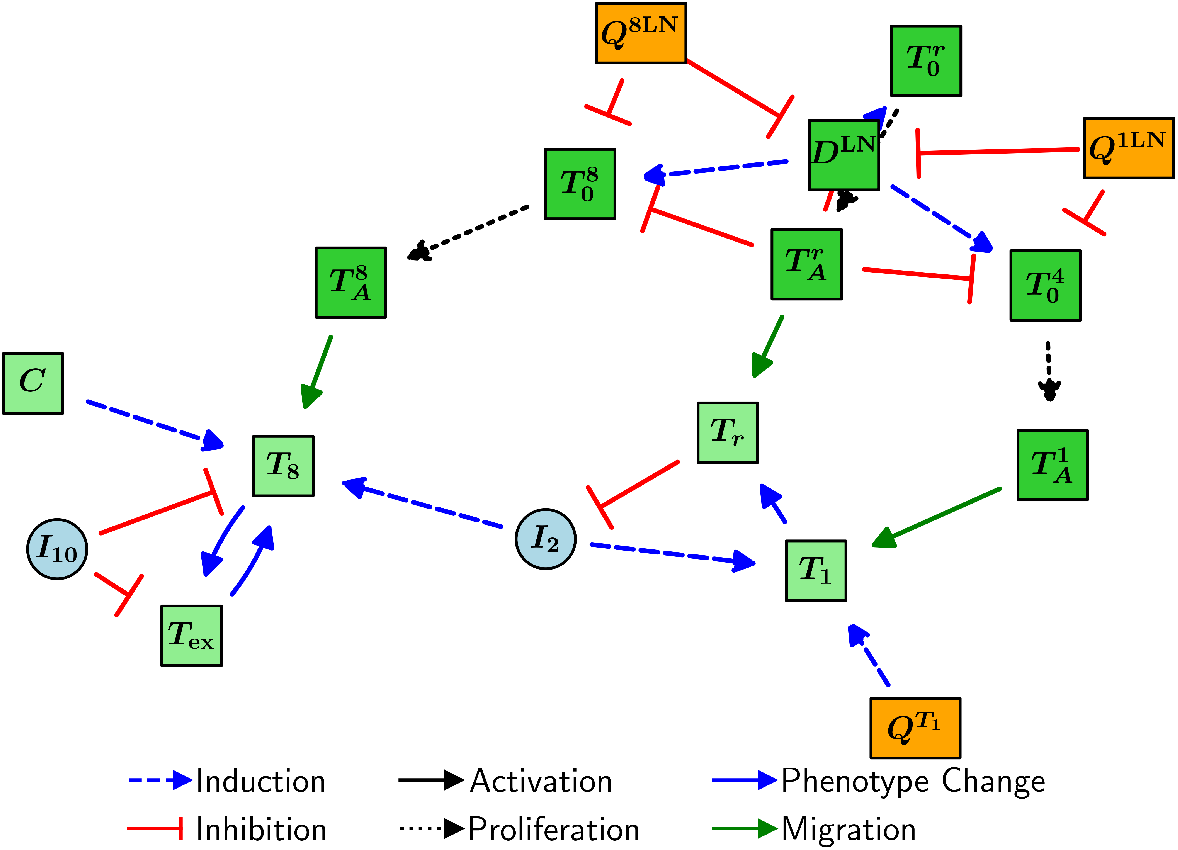
Schematic diagram of the interactions of T cells in the model.

#### 2.3.3 Equations for Macrophages and NK Cells

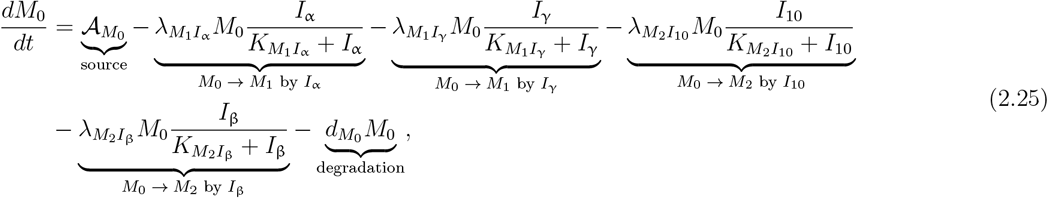

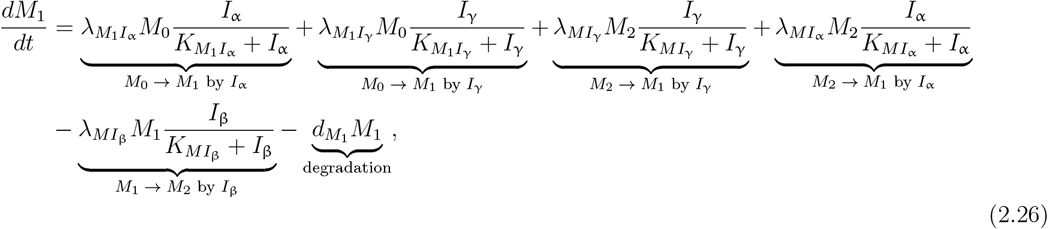

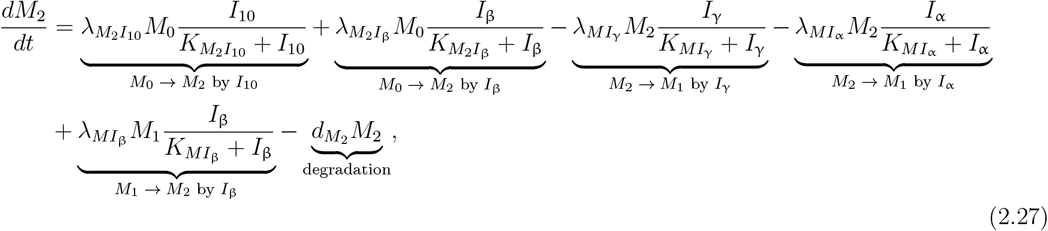

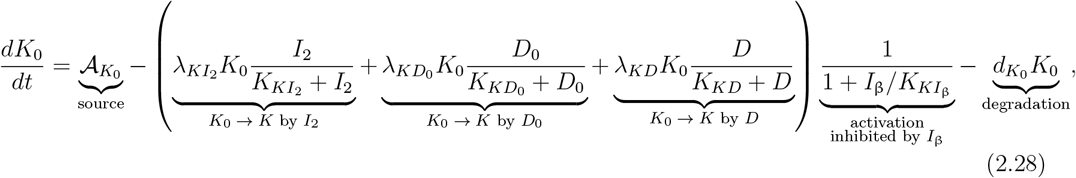

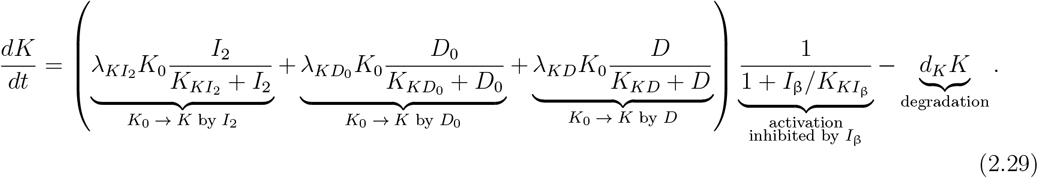

A diagram encompassing the interactions of these components is shown in Figure 3.

**Figure 3:**
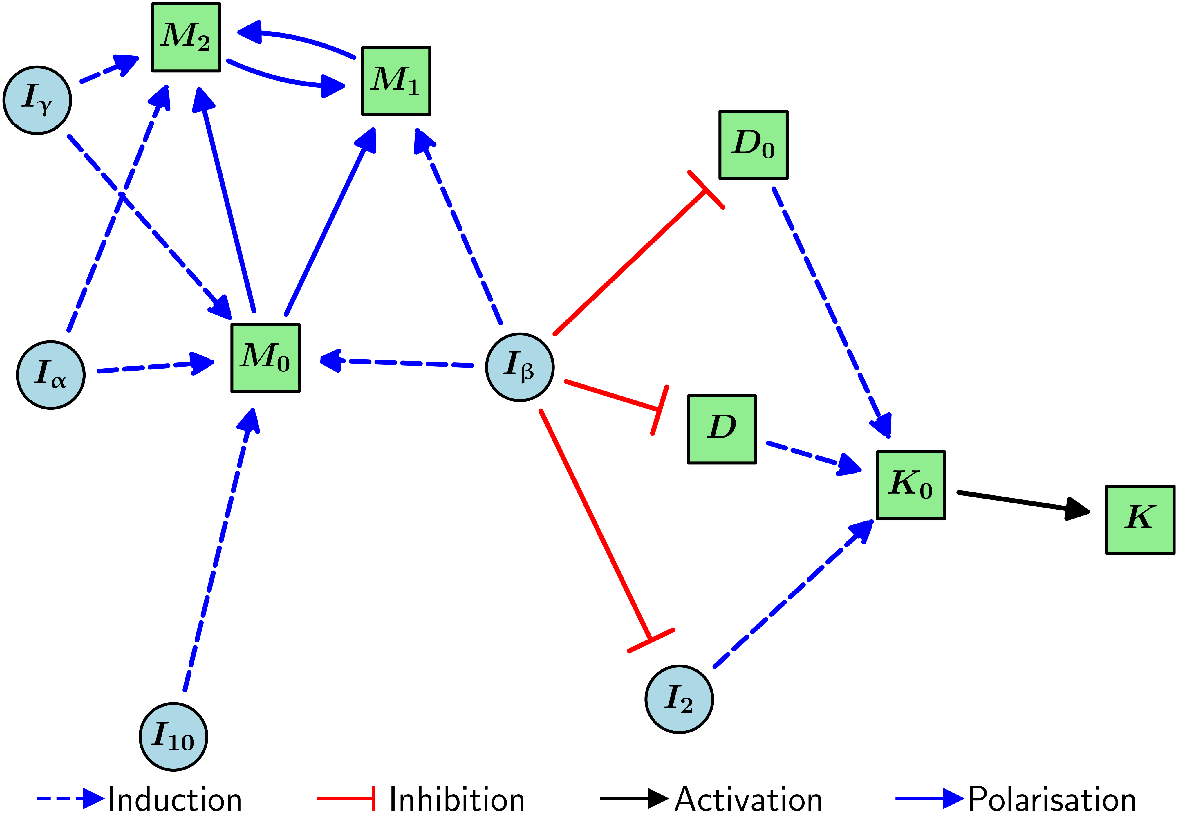
Schematic diagram of the interactions of macrophages and NK cells in the model.

#### 2.3.4 Equations for Cytokines

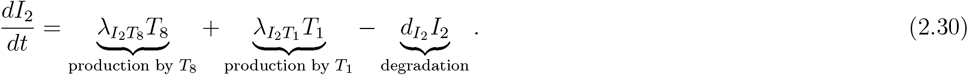

After applying a quasi-steady-state approximation (QSSA), this becomes

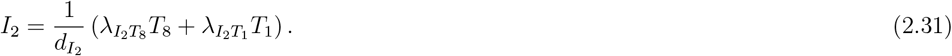

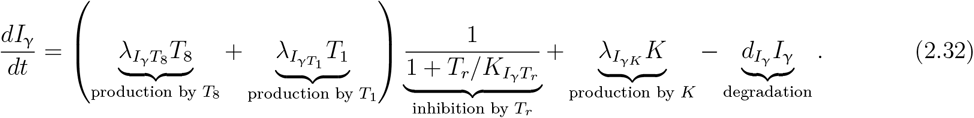

After applying a QSSA, this becomes

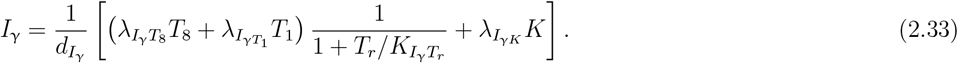

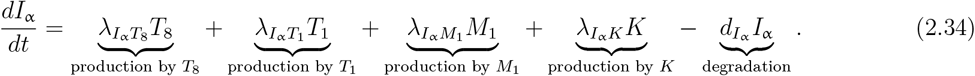

After applying a QSSA, this becomes

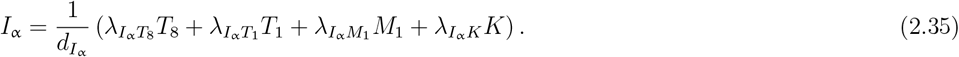

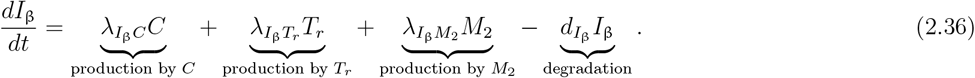

After applying a QSSA, this becomes

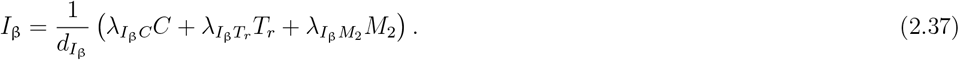

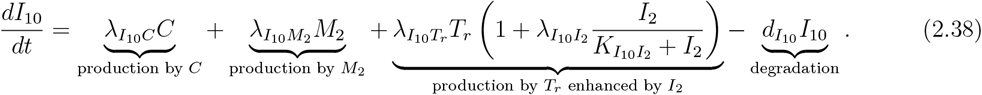

A diagram encompassing the interactions of these components is shown in Figure 4.

**Figure 4:**
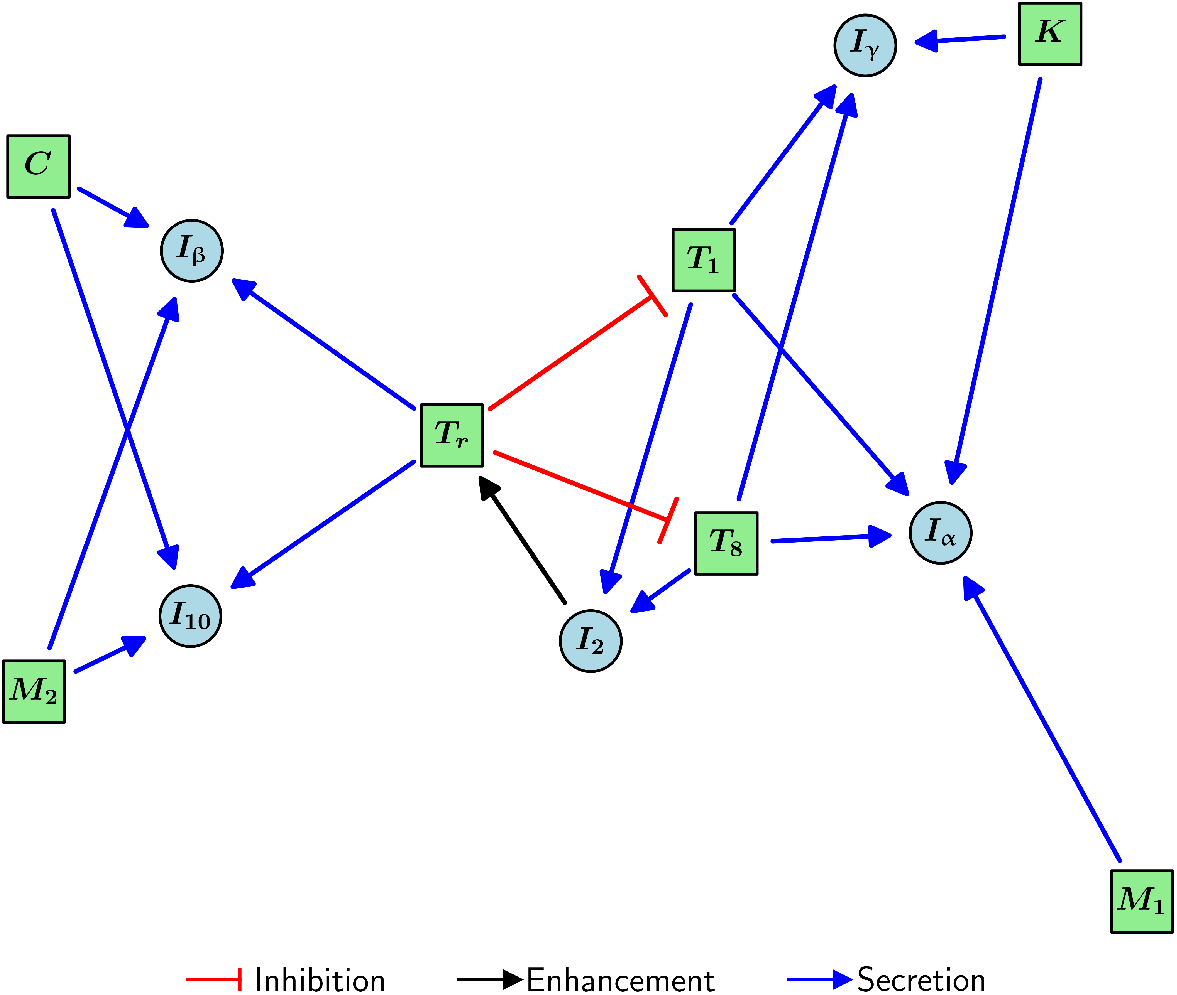
Schematic diagram of the interactions of cytokines in the model.

#### 2.3.5 Equations for Immune Checkpoint-Associated Components in the TS

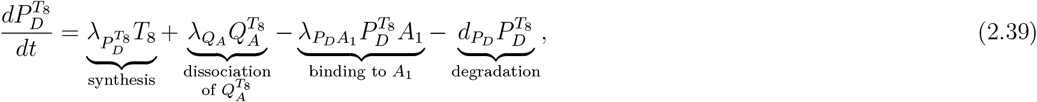

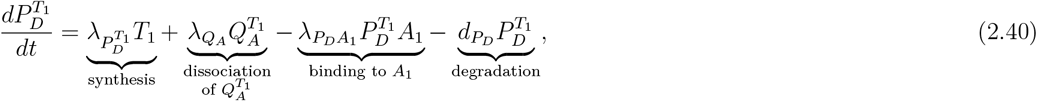

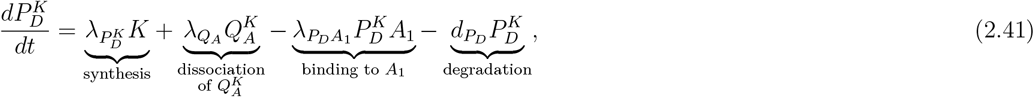

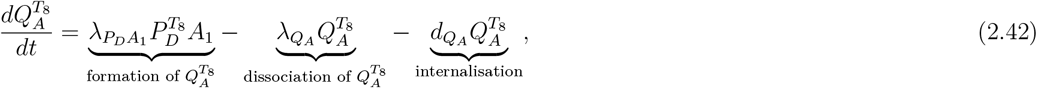

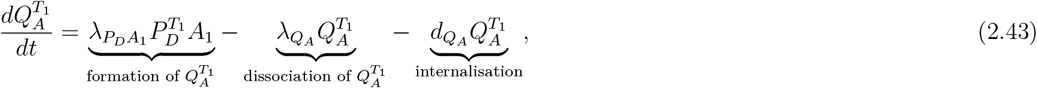

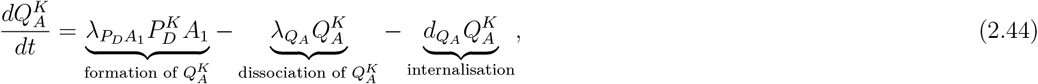

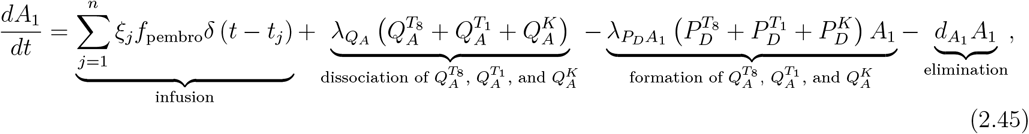

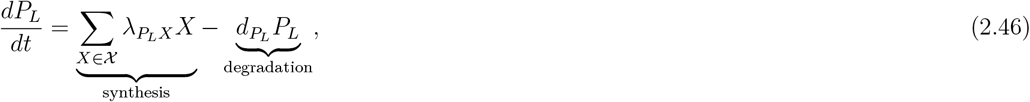

where 𝒳 := }*C, D, T*_8_, *T*_1_, *T*_*r*_, *M*_2_}.

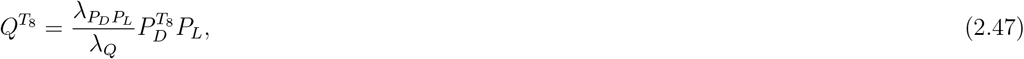

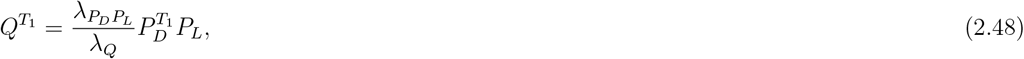

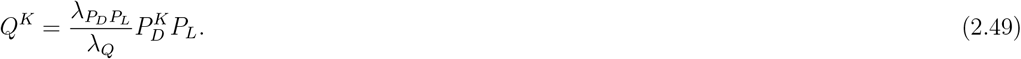

A diagram encompassing the interactions of these components is shown in Figure 5.

**Figure 5:**
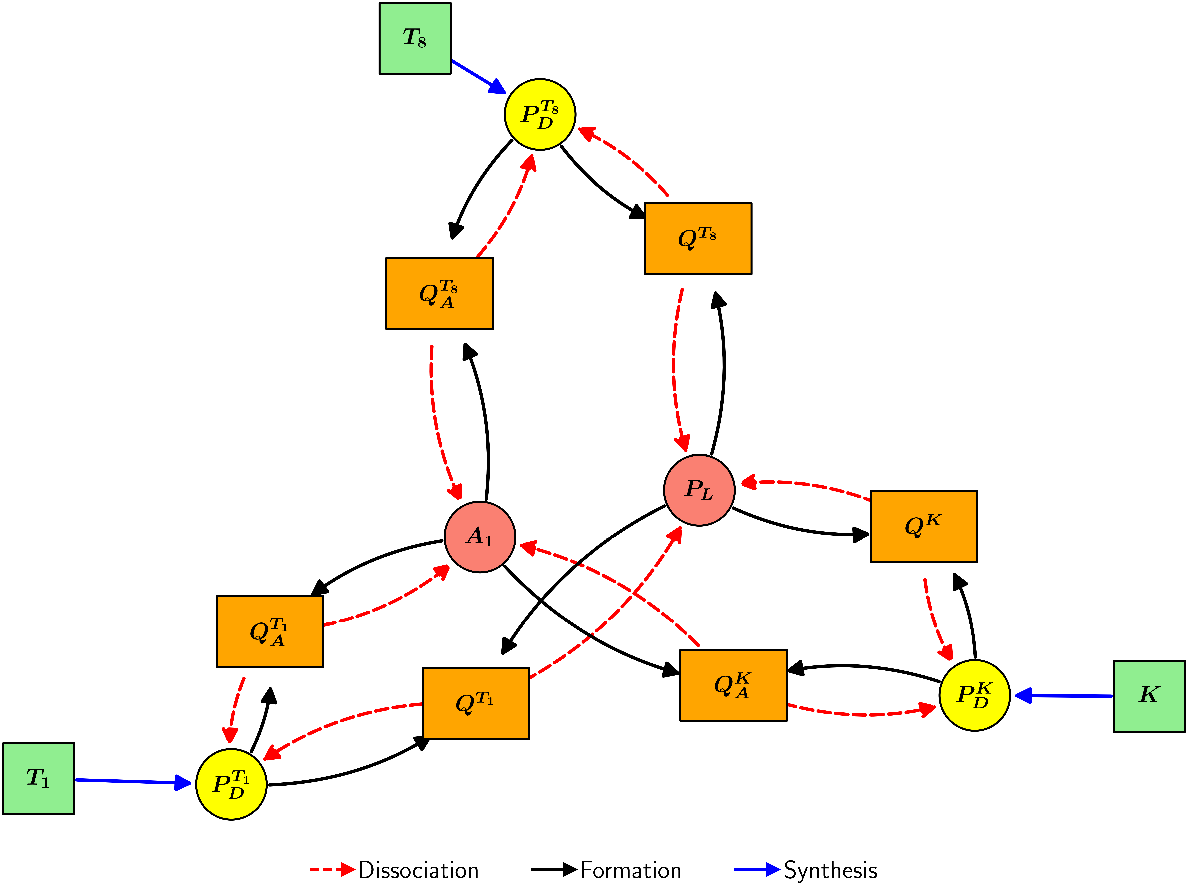
Schematic diagram of the interactions of immune checkpoint-associated components in the TS in the model.

#### 2.3.6 Equations for Immune Checkpoint-Associated Components in the TDLN

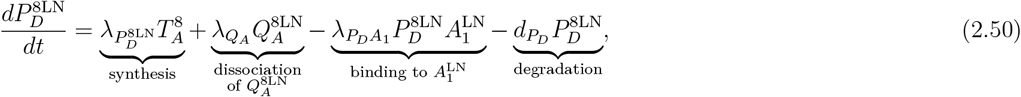

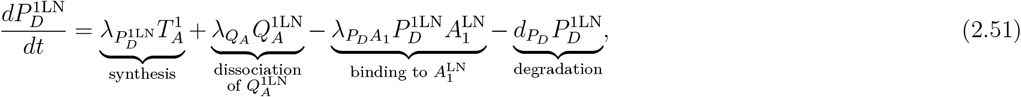

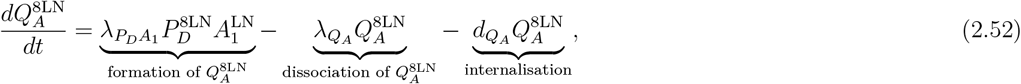

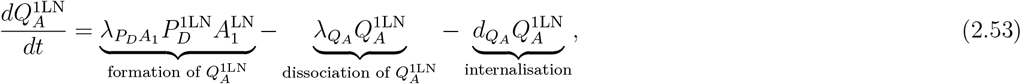

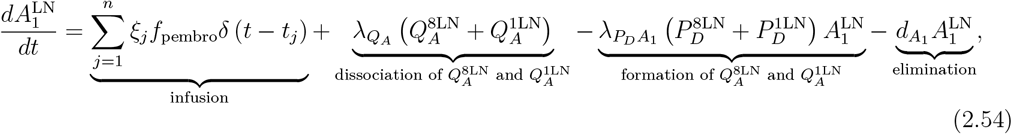

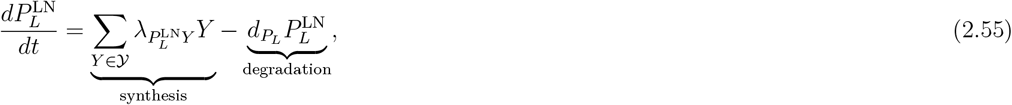

Where 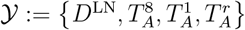.

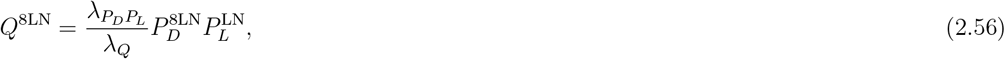

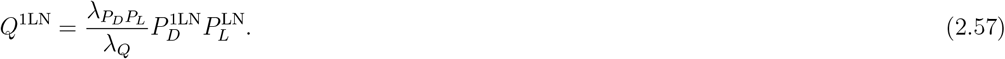

A diagram encompassing the interactions of these components is shown in Figure 6.

**Figure 6:**
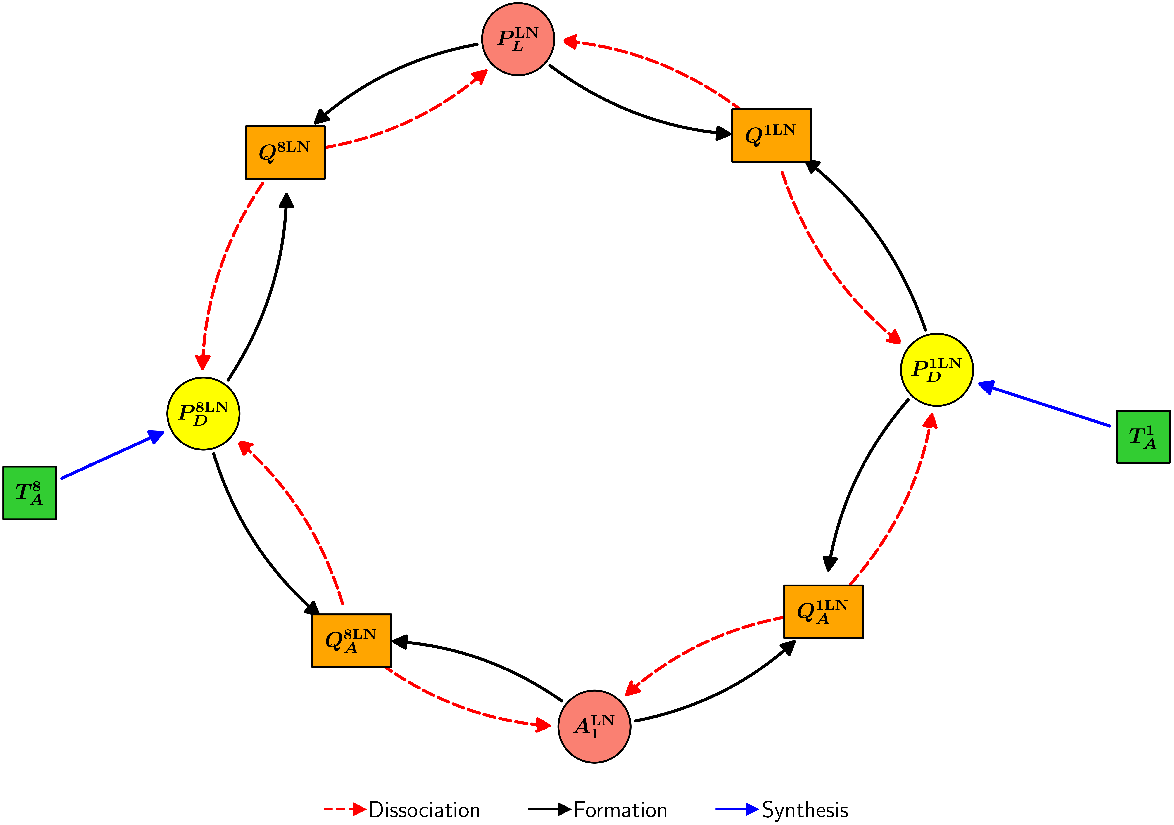
Schematic diagram of the interactions of immune checkpoint-associated components in the TDLN in the model.

### 2.4 Model Parameters

The model parameter values are estimated in Appendix C and are listed in Table 2.

**Table 2:**
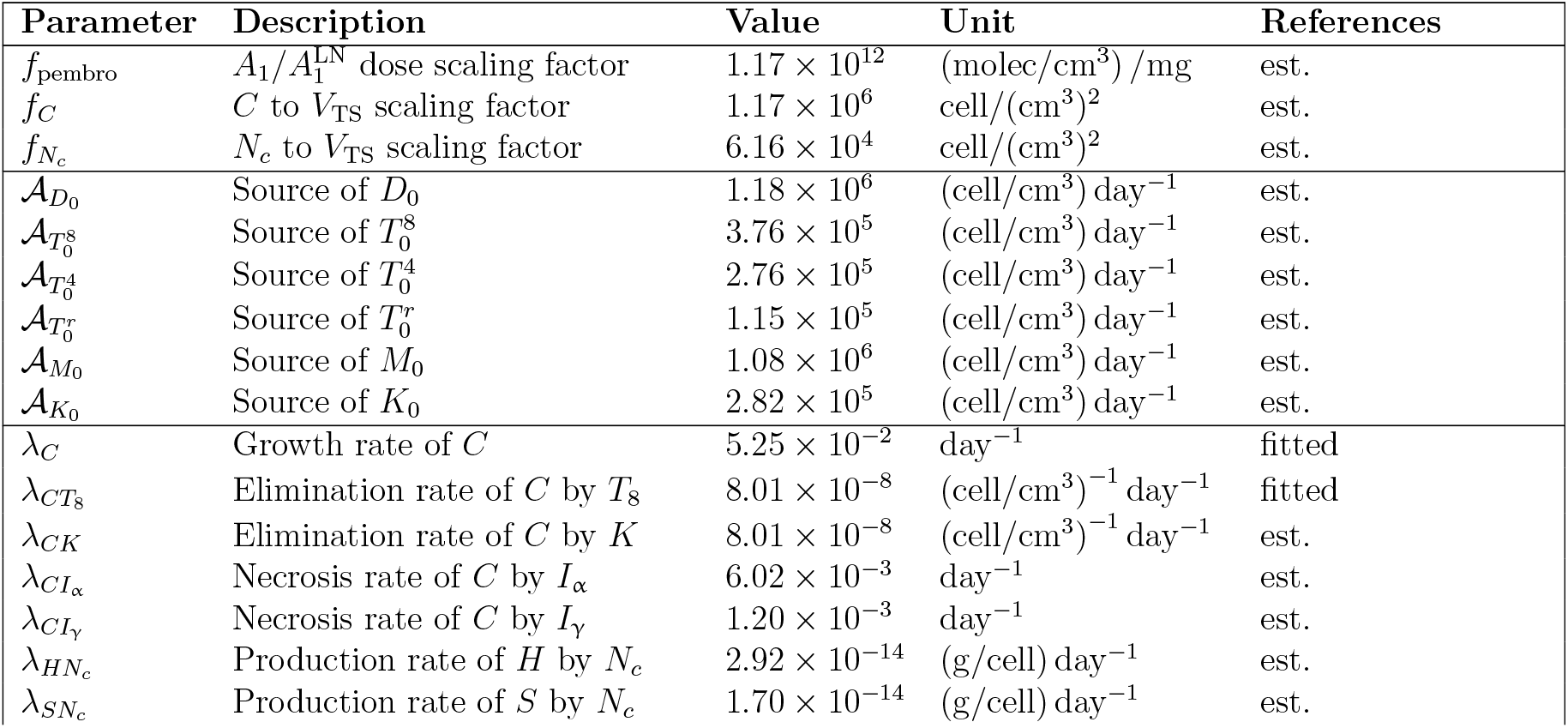
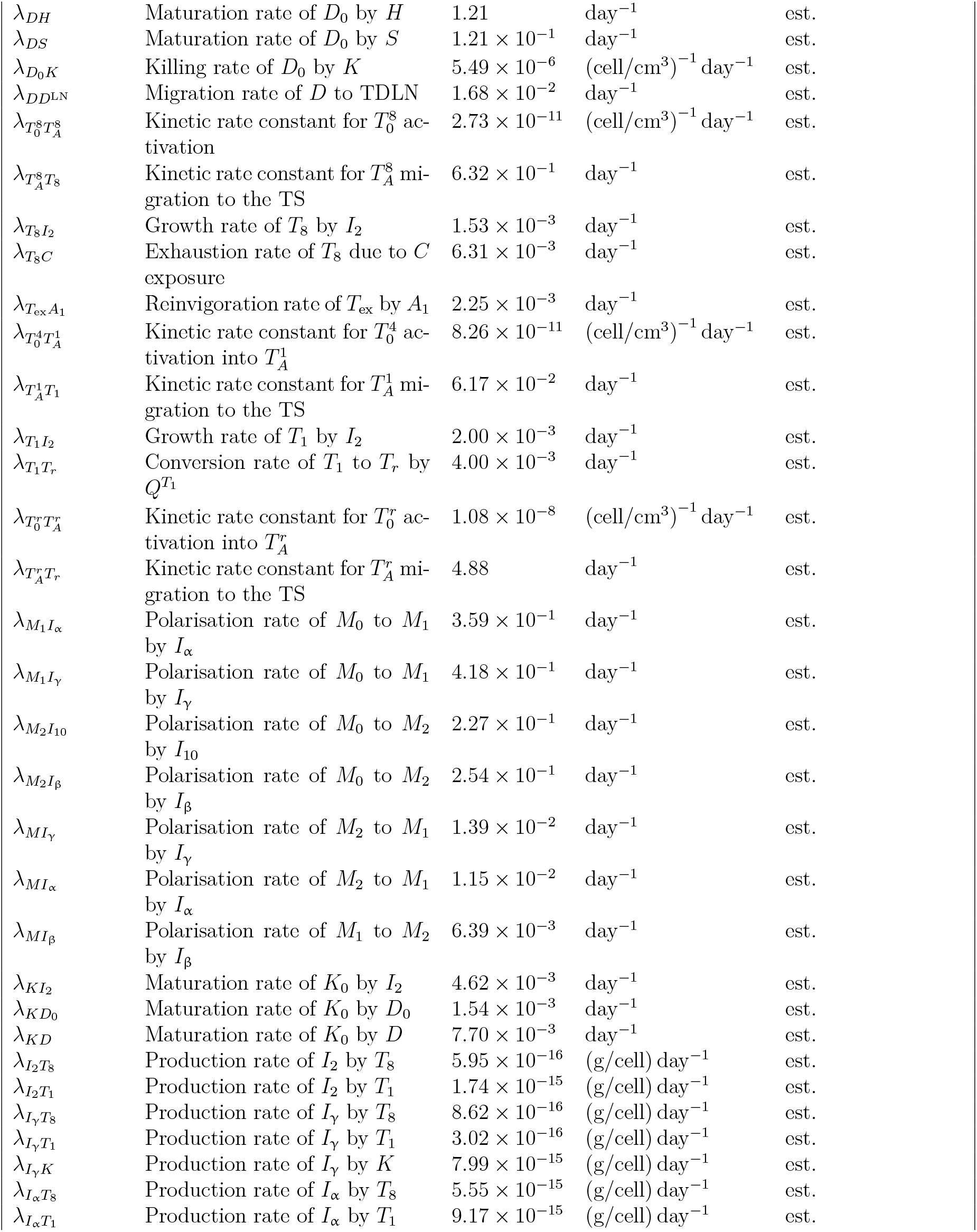
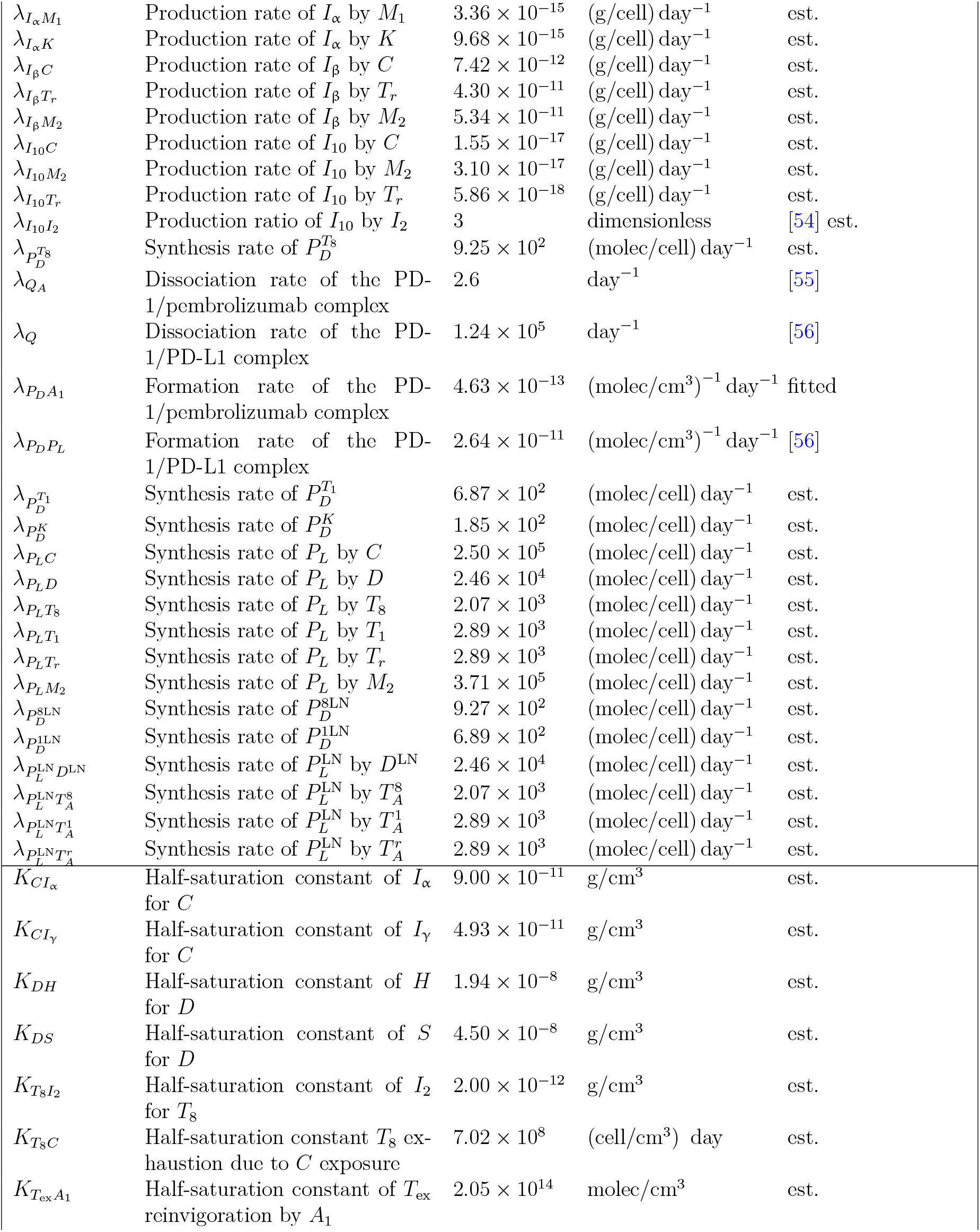
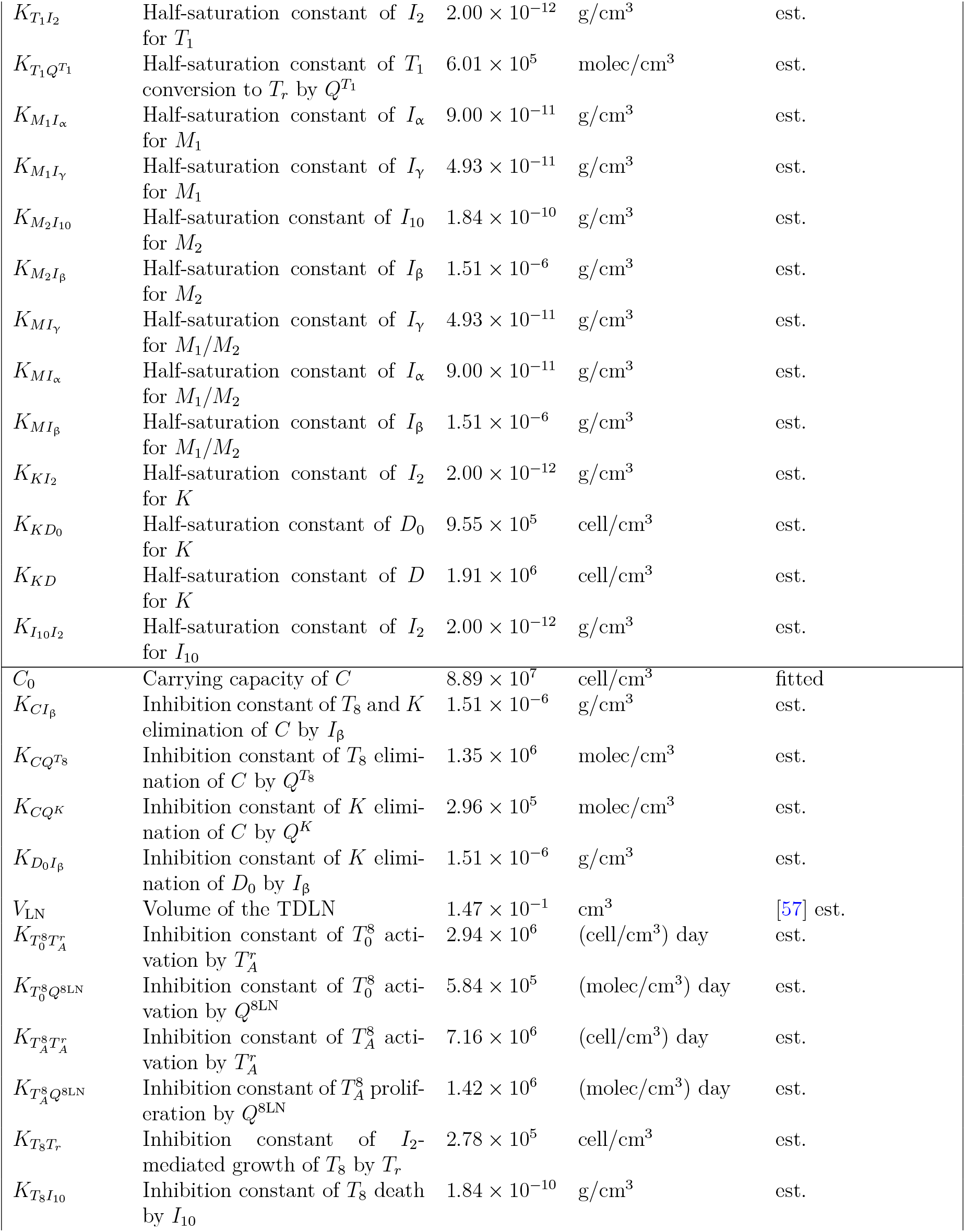
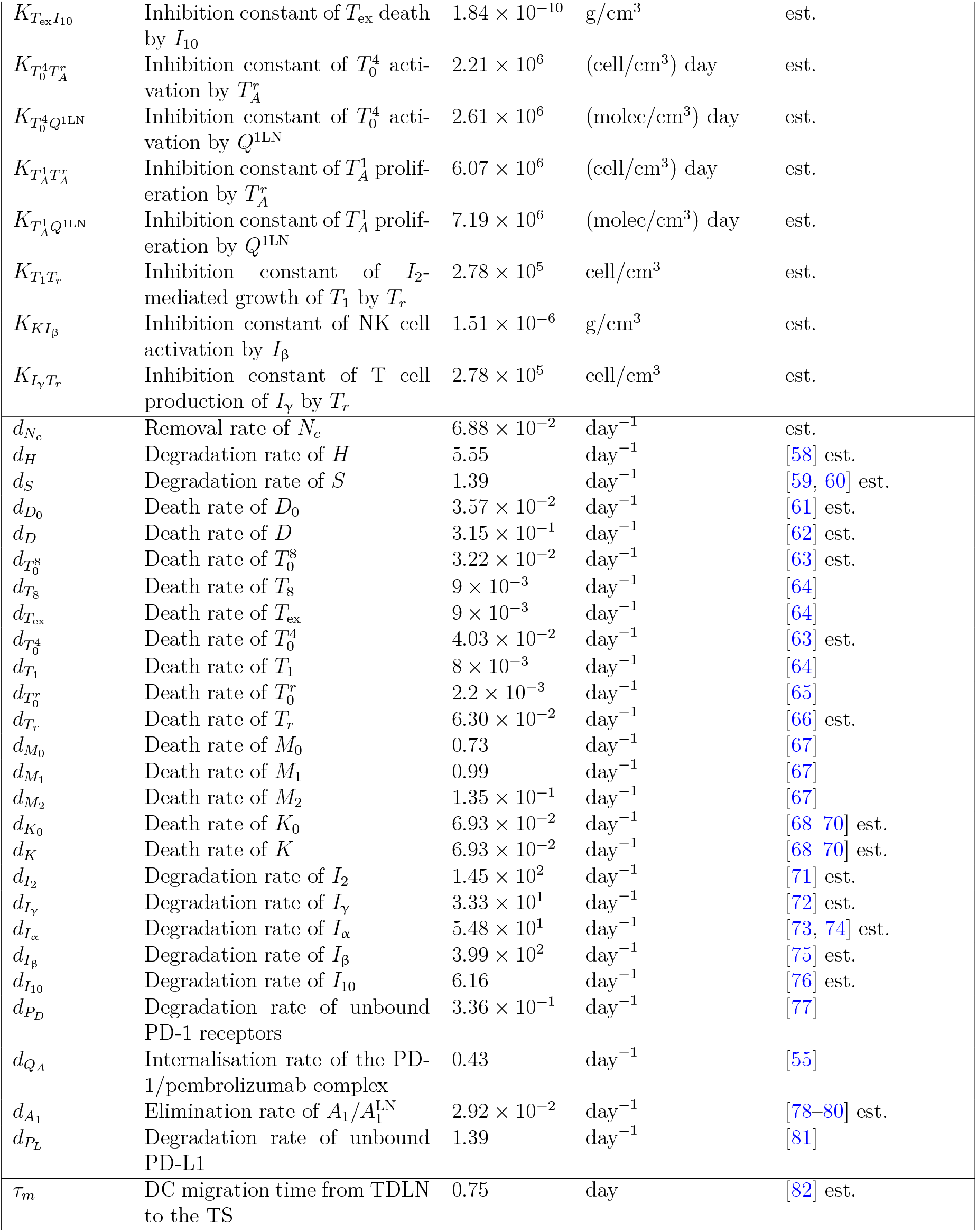
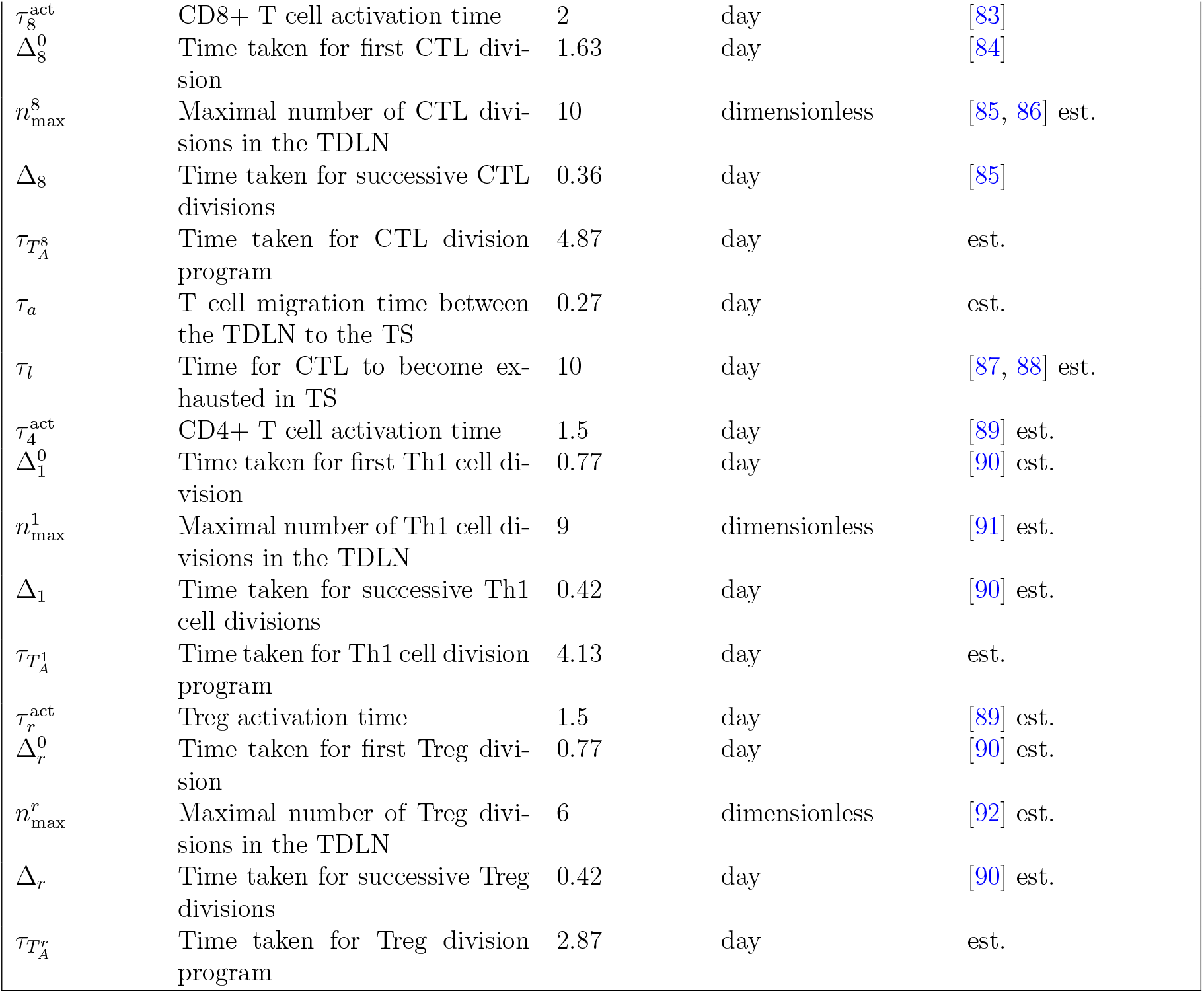
Parameter values for the model. TDLN denotes the tumour-draining lymph node, whilst TS denotes the tumour site. est. denotes estimated parameters.

| Parameter | Description | Value | Unit | References |
| --- | --- | --- | --- | --- |
| $f_{\text{pembro}}$ | $A_1/A_1^{\text{LN}}$ dose scaling factor | $1.17 \times 10^{12}$ | (molec/cm<sup>3</sup>) /mg | est. |
| $f_C$ | $C$ to $V_{\text{TS}}$ scaling factor | $1.17 \times 10^6$ | cell/(cm<sup>3</sup>)<sup>2</sup> | est. |
| $f_{N_c}$ | $N_c$ to $V_{\text{TS}}$ scaling factor | $6.16 \times 10^4$ | cell/(cm<sup>3</sup>)<sup>2</sup> | est. |
| $\mathcal{A}_{D_0}$ | Source of $D_0$ | $1.18 \times 10^6$ | (cell/cm<sup>3</sup>) day<sup>-1</sup> | est. |
| $\mathcal{A}_{T_0^8}$ | Source of $T_0^8$ | $3.76 \times 10^5$ | (cell/cm<sup>3</sup>) day<sup>-1</sup> | est. |
| $\mathcal{A}_{T_0^4}$ | Source of $T_0^4$ | $2.76 \times 10^5$ | (cell/cm<sup>3</sup>) day<sup>-1</sup> | est. |
| $\mathcal{A}_{T_0^r}$ | Source of $T_0^r$ | $1.15 \times 10^5$ | (cell/cm<sup>3</sup>) day<sup>-1</sup> | est. |
| $\mathcal{A}_{M_0}$ | Source of $M_0$ | $1.08 \times 10^6$ | (cell/cm<sup>3</sup>) day<sup>-1</sup> | est. |
| $\mathcal{A}_{K_0}$ | Source of $K_0$ | $2.82 \times 10^5$ | (cell/cm<sup>3</sup>) day<sup>-1</sup> | est. |
| $\lambda_C$ | Growth rate of $C$ | $5.25 \times 10^{-2}$ | day<sup>-1</sup> | fitted |
| $\lambda_{CT_8}$ | Elimination rate of $C$ by $T_8$ | $8.01 \times 10^{-8}$ | (cell/cm<sup>3</sup>)<sup>-1</sup> day<sup>-1</sup> | fitted |
| $\lambda_{CK}$ | Elimination rate of $C$ by $K$ | $8.01 \times 10^{-8}$ | (cell/cm<sup>3</sup>)<sup>-1</sup> day<sup>-1</sup> | est. |
| $\lambda_{CI_\alpha}$ | Necrosis rate of $C$ by $I_\alpha$ | $6.02 \times 10^{-3}$ | day<sup>-1</sup> | est. |
| $\lambda_{CI_\gamma}$ | Necrosis rate of $C$ by $I_\gamma$ | $1.20 \times 10^{-3}$ | day<sup>-1</sup> | est. |
| $\lambda_{HN_c}$ | Production rate of $H$ by $N_c$ | $2.92 \times 10^{-14}$ | (g/cell) day<sup>-1</sup> | est. |
| $\lambda_{SN_c}$ | Production rate of $S$ by $N_c$ | $1.70 \times 10^{-14}$ | (g/cell) day<sup>-1</sup> | est. |
| $\lambda_{DH}$ | Maturation rate of $D_0$ by $H$ | 1.21 | day<sup>-1</sup> | est. |
| $\lambda_{DS}$ | Maturation rate of $D_0$ by $S$ | $1.21 \times 10^{-1}$ | day<sup>-1</sup> | est. |
| $\lambda_{D_0K}$ | Killing rate of $D_0$ by $K$ | $5.49 \times 10^{-6}$ | (cell/cm<sup>3</sup>)<sup>-1</sup> day<sup>-1</sup> | est. |
| $\lambda_{DD^{LN}}$ | Migration rate of $D$ to TDLN | $1.68 \times 10^{-2}$ | day<sup>-1</sup> | est. |
| $\lambda_{T_0^8 T_A^8}$ | Kinetic rate constant for $T_0^8$ activation | $2.73 \times 10^{-11}$ | (cell/cm<sup>3</sup>)<sup>-1</sup> day<sup>-1</sup> | est. |
| $\lambda_{T_A^8 T_8}$ | Kinetic rate constant for $T_A^8$ migration to the TS | $6.32 \times 10^{-1}$ | day<sup>-1</sup> | est. |
| $\lambda_{T_8 I_2}$ | Growth rate of $T_8$ by $I_2$ | $1.53 \times 10^{-3}$ | day<sup>-1</sup> | est. |
| $\lambda_{T_8 C}$ | Exhaustion rate of $T_8$ due to $C$ exposure | $6.31 \times 10^{-3}$ | day<sup>-1</sup> | est. |
| $\lambda_{T_{ex} A_1}$ | Reinvigoration rate of $T_{ex}$ by $A_1$ | $2.25 \times 10^{-3}$ | day<sup>-1</sup> | est. |
| $\lambda_{T_0^4 T_A^1}$ | Kinetic rate constant for $T_0^4$ activation into $T_A^1$ | $8.26 \times 10^{-11}$ | (cell/cm<sup>3</sup>)<sup>-1</sup> day<sup>-1</sup> | est. |
| $\lambda_{T_A^1 T_1}$ | Kinetic rate constant for $T_A^1$ migration to the TS | $6.17 \times 10^{-2}$ | day<sup>-1</sup> | est. |
| $\lambda_{T_1 I_2}$ | Growth rate of $T_1$ by $I_2$ | $2.00 \times 10^{-3}$ | day<sup>-1</sup> | est. |
| $\lambda_{T_1 T_r}$ | Conversion rate of $T_1$ to $T_r$ by $Q^{T_1}$ | $4.00 \times 10^{-3}$ | day<sup>-1</sup> | est. |
| $\lambda_{T_0^r T_A^r}$ | Kinetic rate constant for $T_0^r$ activation into $T_A^r$ | $1.08 \times 10^{-8}$ | (cell/cm<sup>3</sup>)<sup>-1</sup> day<sup>-1</sup> | est. |
| $\lambda_{T_A^r T_r}$ | Kinetic rate constant for $T_A^r$ migration to the TS | 4.88 | day<sup>-1</sup> | est. |
| $\lambda_{M_1 I_\alpha}$ | Polarisation rate of $M_0$ to $M_1$ by $I_\alpha$ | $3.59 \times 10^{-1}$ | day<sup>-1</sup> | est. |
| $\lambda_{M_1 I_\gamma}$ | Polarisation rate of $M_0$ to $M_1$ by $I_\gamma$ | $4.18 \times 10^{-1}$ | day<sup>-1</sup> | est. |
| $\lambda_{M_2 I_{10}}$ | Polarisation rate of $M_0$ to $M_2$ by $I_{10}$ | $2.27 \times 10^{-1}$ | day<sup>-1</sup> | est. |
| $\lambda_{M_2 I_\beta}$ | Polarisation rate of $M_0$ to $M_2$ by $I_\beta$ | $2.54 \times 10^{-1}$ | day<sup>-1</sup> | est. |
| $\lambda_{M I_\gamma}$ | Polarisation rate of $M_2$ to $M_1$ by $I_\gamma$ | $1.39 \times 10^{-2}$ | day<sup>-1</sup> | est. |
| $\lambda_{M I_\alpha}$ | Polarisation rate of $M_2$ to $M_1$ by $I_\alpha$ | $1.15 \times 10^{-2}$ | day<sup>-1</sup> | est. |
| $\lambda_{M I_\beta}$ | Polarisation rate of $M_1$ to $M_2$ by $I_\beta$ | $6.39 \times 10^{-3}$ | day<sup>-1</sup> | est. |
| $\lambda_{K I_2}$ | Maturation rate of $K_0$ by $I_2$ | $4.62 \times 10^{-3}$ | day<sup>-1</sup> | est. |
| $\lambda_{K D_0}$ | Maturation rate of $K_0$ by $D_0$ | $1.54 \times 10^{-3}$ | day<sup>-1</sup> | est. |
| $\lambda_{K D}$ | Maturation rate of $K_0$ by $D$ | $7.70 \times 10^{-3}$ | day<sup>-1</sup> | est. |
| $\lambda_{I_2 T_8}$ | Production rate of $I_2$ by $T_8$ | $5.95 \times 10^{-16}$ | (g/cell) day<sup>-1</sup> | est. |
| $\lambda_{I_2 T_1}$ | Production rate of $I_2$ by $T_1$ | $1.74 \times 10^{-15}$ | (g/cell) day<sup>-1</sup> | est. |
| $\lambda_{I_\gamma T_8}$ | Production rate of $I_\gamma$ by $T_8$ | $8.62 \times 10^{-16}$ | (g/cell) day<sup>-1</sup> | est. |
| $\lambda_{I_\gamma T_1}$ | Production rate of $I_\gamma$ by $T_1$ | $3.02 \times 10^{-16}$ | (g/cell) day<sup>-1</sup> | est. |
| $\lambda_{I_\gamma K}$ | Production rate of $I_\gamma$ by $K$ | $7.99 \times 10^{-15}$ | (g/cell) day<sup>-1</sup> | est. |
| $\lambda_{I_\alpha T_8}$ | Production rate of $I_\alpha$ by $T_8$ | $5.55 \times 10^{-15}$ | (g/cell) day<sup>-1</sup> | est. |
| $\lambda_{I_\alpha T_1}$ | Production rate of $I_\alpha$ by $T_1$ | $9.17 \times 10^{-15}$ | (g/cell) day<sup>-1</sup> | est. |
| $\lambda_{I_\alpha M_1}$ | Production rate of $I_\alpha$ by $M_1$ | $3.36 \times 10^{-15}$ | (g/cell) day<sup>-1</sup> | est. |
| $\lambda_{I_\alpha K}$ | Production rate of $I_\alpha$ by $K$ | $9.68 \times 10^{-15}$ | (g/cell) day<sup>-1</sup> | est. |
| $\lambda_{I_\beta C}$ | Production rate of $I_\beta$ by $C$ | $7.42 \times 10^{-12}$ | (g/cell) day<sup>-1</sup> | est. |
| $\lambda_{I_\beta T_r}$ | Production rate of $I_\beta$ by $T_r$ | $4.30 \times 10^{-11}$ | (g/cell) day<sup>-1</sup> | est. |
| $\lambda_{I_\beta M_2}$ | Production rate of $I_\beta$ by $M_2$ | $5.34 \times 10^{-11}$ | (g/cell) day<sup>-1</sup> | est. |
| $\lambda_{I_{10} C}$ | Production rate of $I_{10}$ by $C$ | $1.55 \times 10^{-17}$ | (g/cell) day<sup>-1</sup> | est. |
| $\lambda_{I_{10} M_2}$ | Production rate of $I_{10}$ by $M_2$ | $3.10 \times 10^{-17}$ | (g/cell) day<sup>-1</sup> | est. |
| $\lambda_{I_{10} T_r}$ | Production rate of $I_{10}$ by $T_r$ | $5.86 \times 10^{-18}$ | (g/cell) day<sup>-1</sup> | est. |
| $\lambda_{I_{10} I_2}$ | Production ratio of $I_{10}$ by $I_2$ | 3 | dimensionless | [54] est. |
| $\lambda_{P_D^{T_8}}$ | Synthesis rate of $P_D^{T_8}$ | $9.25 \times 10^2$ | (molec/cell) day<sup>-1</sup> | est. |
| $\lambda_{Q_A}$ | Dissociation rate of the PD-1/pembrolizumab complex | 2.6 | day<sup>-1</sup> | [55] |
| $\lambda_Q$ | Dissociation rate of the PD-1/PD-L1 complex | $1.24 \times 10^5$ | day<sup>-1</sup> | [56] |
| $\lambda_{P_D A_1}$ | Formation rate of the PD-1/pembrolizumab complex | $4.63 \times 10^{-13}$ | (molec/cm<sup>3</sup>)<sup>-1</sup> day<sup>-1</sup> | fitted |
| $\lambda_{P_D P_L}$ | Formation rate of the PD-1/PD-L1 complex | $2.64 \times 10^{-11}$ | (molec/cm<sup>3</sup>)<sup>-1</sup> day<sup>-1</sup> | [56] |
| $\lambda_{P_D^{T_1}}$ | Synthesis rate of $P_D^{T_1}$ | $6.87 \times 10^2$ | (molec/cell) day<sup>-1</sup> | est. |
| $\lambda_{P_D^K}$ | Synthesis rate of $P_D^K$ | $1.85 \times 10^2$ | (molec/cell) day<sup>-1</sup> | est. |
| $\lambda_{P_L C}$ | Synthesis rate of $P_L$ by $C$ | $2.50 \times 10^5$ | (molec/cell) day<sup>-1</sup> | est. |
| $\lambda_{P_L D}$ | Synthesis rate of $P_L$ by $D$ | $2.46 \times 10^4$ | (molec/cell) day<sup>-1</sup> | est. |
| $\lambda_{P_L T_8}$ | Synthesis rate of $P_L$ by $T_8$ | $2.07 \times 10^3$ | (molec/cell) day<sup>-1</sup> | est. |
| $\lambda_{P_L T_1}$ | Synthesis rate of $P_L$ by $T_1$ | $2.89 \times 10^3$ | (molec/cell) day<sup>-1</sup> | est. |
| $\lambda_{P_L T_r}$ | Synthesis rate of $P_L$ by $T_r$ | $2.89 \times 10^3$ | (molec/cell) day<sup>-1</sup> | est. |
| $\lambda_{P_L M_2}$ | Synthesis rate of $P_L$ by $M_2$ | $3.71 \times 10^5$ | (molec/cell) day<sup>-1</sup> | est. |
| $\lambda_{P_D^{8LN}}$ | Synthesis rate of $P_D^{8LN}$ | $9.27 \times 10^2$ | (molec/cell) day<sup>-1</sup> | est. |
| $\lambda_{P_D^{1LN}}$ | Synthesis rate of $P_D^{1LN}$ | $6.89 \times 10^2$ | (molec/cell) day<sup>-1</sup> | est. |
| $\lambda_{P_L^{LN} D^{LN}}$ | Synthesis rate of $P_L^{LN}$ by $D^{LN}$ | $2.46 \times 10^4$ | (molec/cell) day<sup>-1</sup> | est. |
| $\lambda_{P_L^{LN} T_A^8}$ | Synthesis rate of $P_L^{LN}$ by $T_A^8$ | $2.07 \times 10^3$ | (molec/cell) day<sup>-1</sup> | est. |
| $\lambda_{P_L^{LN} T_A^1}$ | Synthesis rate of $P_L^{LN}$ by $T_A^1$ | $2.89 \times 10^3$ | (molec/cell) day<sup>-1</sup> | est. |
| $\lambda_{P_L^{LN} T_A^r}$ | Synthesis rate of $P_L^{LN}$ by $T_A^r$ | $2.89 \times 10^3$ | (molec/cell) day<sup>-1</sup> | est. |
| $K_{CI_\alpha}$ | Half-saturation constant of $I_\alpha$ for $C$ | $9.00 \times 10^{-11}$ | g/cm<sup>3</sup> | est. |
| $K_{CI_\gamma}$ | Half-saturation constant of $I_\gamma$ for $C$ | $4.93 \times 10^{-11}$ | g/cm<sup>3</sup> | est. |
| $K_{DH}$ | Half-saturation constant of $H$ for $D$ | $1.94 \times 10^{-8}$ | g/cm<sup>3</sup> | est. |
| $K_{DS}$ | Half-saturation constant of $S$ for $D$ | $4.50 \times 10^{-8}$ | g/cm<sup>3</sup> | est. |
| $K_{T_8 I_2}$ | Half-saturation constant of $I_2$ for $T_8$ | $2.00 \times 10^{-12}$ | g/cm<sup>3</sup> | est. |
| $K_{T_8 C}$ | Half-saturation constant $T_8$ exhaustion due to $C$ exposure | $7.02 \times 10^8$ | (cell/cm<sup>3</sup>) day | est. |
| $K_{T_{ex} A_1}$ | Half-saturation constant of $T_{ex}$ reinvigoration by $A_1$ | $2.05 \times 10^{14}$ | molec/cm<sup>3</sup> | est. |
| $K_{T_1 I_2}$ | Half-saturation constant of $I_2$ for $T_1$ | $2.00 \times 10^{-12}$ | g/cm<sup>3</sup> | est. |
| $K_{T_1 Q^{T_1}}$ | Half-saturation constant of $T_1$ conversion to $T_r$ by $Q^{T_1}$ | $6.01 \times 10^5$ | molec/cm<sup>3</sup> | est. |
| $K_{M_1 I_\alpha}$ | Half-saturation constant of $I_\alpha$ for $M_1$ | $9.00 \times 10^{-11}$ | g/cm<sup>3</sup> | est. |
| $K_{M_1 I_\gamma}$ | Half-saturation constant of $I_\gamma$ for $M_1$ | $4.93 \times 10^{-11}$ | g/cm<sup>3</sup> | est. |
| $K_{M_2 I_{10}}$ | Half-saturation constant of $I_{10}$ for $M_2$ | $1.84 \times 10^{-10}$ | g/cm<sup>3</sup> | est. |
| $K_{M_2 I_\beta}$ | Half-saturation constant of $I_\beta$ for $M_2$ | $1.51 \times 10^{-6}$ | g/cm<sup>3</sup> | est. |
| $K_{M I_\gamma}$ | Half-saturation constant of $I_\gamma$ for $M_1/M_2$ | $4.93 \times 10^{-11}$ | g/cm<sup>3</sup> | est. |
| $K_{M I_\alpha}$ | Half-saturation constant of $I_\alpha$ for $M_1/M_2$ | $9.00 \times 10^{-11}$ | g/cm<sup>3</sup> | est. |
| $K_{M I_\beta}$ | Half-saturation constant of $I_\beta$ for $M_1/M_2$ | $1.51 \times 10^{-6}$ | g/cm<sup>3</sup> | est. |
| $K_{K I_2}$ | Half-saturation constant of $I_2$ for $K$ | $2.00 \times 10^{-12}$ | g/cm<sup>3</sup> | est. |
| $K_{K D_0}$ | Half-saturation constant of $D_0$ for $K$ | $9.55 \times 10^5$ | cell/cm<sup>3</sup> | est. |
| $K_{K D}$ | Half-saturation constant of $D$ for $K$ | $1.91 \times 10^6$ | cell/cm<sup>3</sup> | est. |
| $K_{I_{10} I_2}$ | Half-saturation constant of $I_2$ for $I_{10}$ | $2.00 \times 10^{-12}$ | g/cm<sup>3</sup> | est. |
| $C_0$ | Carrying capacity of $C$ | $8.89 \times 10^7$ | cell/cm<sup>3</sup> | fitted |
| $K_{C I_\beta}$ | Inhibition constant of $T_8$ and $K$ elimination of $C$ by $I_\beta$ | $1.51 \times 10^{-6}$ | g/cm<sup>3</sup> | est. |
| $K_{C Q^{T_8}}$ | Inhibition constant of $T_8$ elimination of $C$ by $Q^{T_8}$ | $1.35 \times 10^6$ | molec/cm<sup>3</sup> | est. |
| $K_{C Q^K}$ | Inhibition constant of $K$ elimination of $C$ by $Q^K$ | $2.96 \times 10^5$ | molec/cm<sup>3</sup> | est. |
| $K_{D_0 I_\beta}$ | Inhibition constant of $K$ elimination of $D_0$ by $I_\beta$ | $1.51 \times 10^{-6}$ | g/cm<sup>3</sup> | est. |
| $V_{LN}$ | Volume of the TDLN | $1.47 \times 10^{-1}$ | cm<sup>3</sup> | [57] est. |
| $K_{T_0^8 T_A^r}$ | Inhibition constant of $T_0^8$ activation by $T_A^r$ | $2.94 \times 10^6$ | (cell/cm<sup>3</sup>) day | est. |
| $K_{T_0^8 Q^{8LN}}$ | Inhibition constant of $T_0^8$ activation by $Q^{8LN}$ | $5.84 \times 10^5$ | (molec/cm<sup>3</sup>) day | est. |
| $K_{T_A^8 T_A^r}$ | Inhibition constant of $T_A^8$ activation by $T_A^r$ | $7.16 \times 10^6$ | (cell/cm<sup>3</sup>) day | est. |
| $K_{T_A^8 Q^{8LN}}$ | Inhibition constant of $T_A^8$ proliferation by $Q^{8LN}$ | $1.42 \times 10^6$ | (molec/cm<sup>3</sup>) day | est. |
| $K_{T_8 T_r}$ | Inhibition constant of $I_2$-mediated growth of $T_8$ by $T_r$ | $2.78 \times 10^5$ | cell/cm<sup>3</sup> | est. |
| $K_{T_8 I_{10}}$ | Inhibition constant of $T_8$ death by $I_{10}$ | $1.84 \times 10^{-10}$ | g/cm<sup>3</sup> | est. |
| $K_{T_{\text{ex}}I_{10}}$ | Inhibition constant of $T_{\text{ex}}$ death by $I_{10}$ | $1.84 \times 10^{-10}$ | $\text{g}/\text{cm}^3$ | est. |
| $K_{T_0^4 T_A^r}$ | Inhibition constant of $T_0^4$ activation by $T_A^r$ | $2.21 \times 10^6$ | $(\text{cell}/\text{cm}^3) \text{ day}$ | est. |
| $K_{T_0^4 Q^{1\text{LN}}}$ | Inhibition constant of $T_0^4$ activation by $Q^{1\text{LN}}$ | $2.61 \times 10^6$ | $(\text{molec}/\text{cm}^3) \text{ day}$ | est. |
| $K_{T_A^1 T_A^r}$ | Inhibition constant of $T_A^1$ proliferation by $T_A^r$ | $6.07 \times 10^6$ | $(\text{cell}/\text{cm}^3) \text{ day}$ | est. |
| $K_{T_A^1 Q^{1\text{LN}}}$ | Inhibition constant of $T_A^1$ proliferation by $Q^{1\text{LN}}$ | $7.19 \times 10^6$ | $(\text{molec}/\text{cm}^3) \text{ day}$ | est. |
| $K_{T_1 T_r}$ | Inhibition constant of $I_2$-mediated growth of $T_1$ by $T_r$ | $2.78 \times 10^5$ | $\text{cell}/\text{cm}^3$ | est. |
| $K_{K I_\beta}$ | Inhibition constant of NK cell activation by $I_\beta$ | $1.51 \times 10^{-6}$ | $\text{g}/\text{cm}^3$ | est. |
| $K_{I_\gamma T_r}$ | Inhibition constant of T cell production of $I_\gamma$ by $T_r$ | $2.78 \times 10^5$ | $\text{cell}/\text{cm}^3$ | est. |
| $d_{N_c}$ | Removal rate of $N_c$ | $6.88 \times 10^{-2}$ | $\text{day}^{-1}$ | est. |
| $d_H$ | Degradation rate of $H$ | 5.55 | $\text{day}^{-1}$ | [58] est. |
| $d_S$ | Degradation rate of $S$ | 1.39 | $\text{day}^{-1}$ | [59, 60] est. |
| $d_{D_0}$ | Death rate of $D_0$ | $3.57 \times 10^{-2}$ | $\text{day}^{-1}$ | [61] est. |
| $d_D$ | Death rate of $D$ | $3.15 \times 10^{-1}$ | $\text{day}^{-1}$ | [62] est. |
| $d_{T_0^8}$ | Death rate of $T_0^8$ | $3.22 \times 10^{-2}$ | $\text{day}^{-1}$ | [63] est. |
| $d_{T_8}$ | Death rate of $T_8$ | $9 \times 10^{-3}$ | $\text{day}^{-1}$ | [64] |
| $d_{T_{\text{ex}}}$ | Death rate of $T_{\text{ex}}$ | $9 \times 10^{-3}$ | $\text{day}^{-1}$ | [64] |
| $d_{T_0^4}$ | Death rate of $T_0^4$ | $4.03 \times 10^{-2}$ | $\text{day}^{-1}$ | [63] est. |
| $d_{T_1}$ | Death rate of $T_1$ | $8 \times 10^{-3}$ | $\text{day}^{-1}$ | [64] |
| $d_{T_0^r}$ | Death rate of $T_0^r$ | $2.2 \times 10^{-3}$ | $\text{day}^{-1}$ | [65] |
| $d_{T_r}$ | Death rate of $T_r$ | $6.30 \times 10^{-2}$ | $\text{day}^{-1}$ | [66] est. |
| $d_{M_0}$ | Death rate of $M_0$ | 0.73 | $\text{day}^{-1}$ | [67] |
| $d_{M_1}$ | Death rate of $M_1$ | 0.99 | $\text{day}^{-1}$ | [67] |
| $d_{M_2}$ | Death rate of $M_2$ | $1.35 \times 10^{-1}$ | $\text{day}^{-1}$ | [67] |
| $d_{K_0}$ | Death rate of $K_0$ | $6.93 \times 10^{-2}$ | $\text{day}^{-1}$ | [68–70] est. |
| $d_K$ | Death rate of $K$ | $6.93 \times 10^{-2}$ | $\text{day}^{-1}$ | [68–70] est. |
| $d_{I_2}$ | Degradation rate of $I_2$ | $1.45 \times 10^2$ | $\text{day}^{-1}$ | [71] est. |
| $d_{I_\gamma}$ | Degradation rate of $I_\gamma$ | $3.33 \times 10^1$ | $\text{day}^{-1}$ | [72] est. |
| $d_{I_\alpha}$ | Degradation rate of $I_\alpha$ | $5.48 \times 10^1$ | $\text{day}^{-1}$ | [73, 74] est. |
| $d_{I_\beta}$ | Degradation rate of $I_\beta$ | $3.99 \times 10^2$ | $\text{day}^{-1}$ | [75] est. |
| $d_{I_{10}}$ | Degradation rate of $I_{10}$ | 6.16 | $\text{day}^{-1}$ | [76] est. |
| $d_{P_D}$ | Degradation rate of unbound PD-1 receptors | $3.36 \times 10^{-1}$ | $\text{day}^{-1}$ | [77] |
| $d_{Q_A}$ | Internalisation rate of the PD-1/pembrolizumab complex | 0.43 | $\text{day}^{-1}$ | [55] |
| $d_{A_1}$ | Elimination rate of $A_1/A_1^{\text{LN}}$ | $2.92 \times 10^{-2}$ | $\text{day}^{-1}$ | [78–80] est. |
| $d_{P_L}$ | Degradation rate of unbound PD-L1 | 1.39 | $\text{day}^{-1}$ | [81] |
| $\tau_m$ | DC migration time from TDLN to the TS | 0.75 | day | [82] est. |
| $\tau_8^{\text{act}}$ | CD8+ T cell activation time | 2 | day | [83] |
| $\Delta_8^0$ | Time taken for first CTL division | 1.63 | day | [84] |
| $n_{\text{max}}^8$ | Maximal number of CTL divisions in the TDLN | 10 | dimensionless | [85, 86] est. |
| $\Delta_8$ | Time taken for successive CTL divisions | 0.36 | day | [85] |
| $\tau_{T_A}^8$ | Time taken for CTL division program | 4.87 | day | est. |
| $\tau_a$ | T cell migration time between the TDLN to the TS | 0.27 | day | est. |
| $\tau_l$ | Time for CTL to become exhausted in TS | 10 | day | [87, 88] est. |
| $\tau_4^{\text{act}}$ | CD4+ T cell activation time | 1.5 | day | [89] est. |
| $\Delta_1^0$ | Time taken for first Th1 cell division | 0.77 | day | [90] est. |
| $n_{\text{max}}^1$ | Maximal number of Th1 cell divisions in the TDLN | 9 | dimensionless | [91] est. |
| $\Delta_1$ | Time taken for successive Th1 cell divisions | 0.42 | day | [90] est. |
| $\tau_{T_A}^1$ | Time taken for Th1 cell division program | 4.13 | day | est. |
| $\tau_r^{\text{act}}$ | Treg activation time | 1.5 | day | [89] est. |
| $\Delta_r^0$ | Time taken for first Treg division | 0.77 | day | [90] est. |
| $n_{\text{max}}^r$ | Maximal number of Treg divisions in the TDLN | 6 | dimensionless | [92] est. |
| $\Delta_r$ | Time taken for successive Treg divisions | 0.42 | day | [90] est. |
| $\tau_{T_A}^r$ | Time taken for Treg division program | 2.87 | day | est. |

## 3 Initial Conditions

For all species in the model, we assume a constant solution history, where the history for each species is set to its respective initial condition.

### 3.1 Initial Conditions for Cells in the TS

We chose the initial conditions for cells in the TS to be as in Table 3, with justification for the choice of these values in Appendix B.1.

**Table 3:**
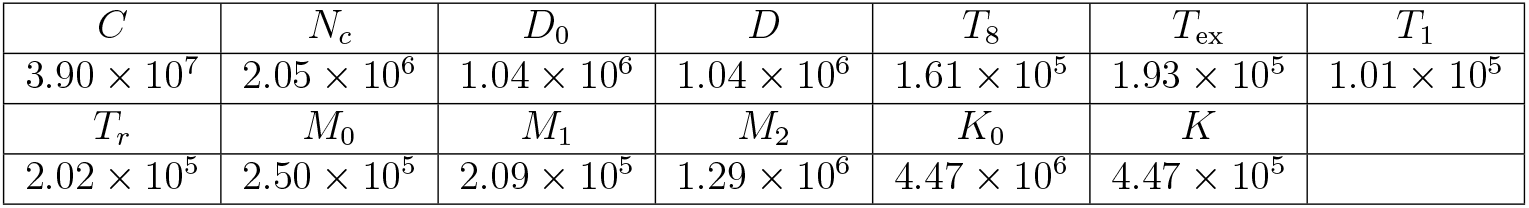
TS initial condition cell densities for the model. All values are in cell/cm^3^.

### 3.2 Initial Conditions for Cells in the TDLN

We estimated the initial conditions for T cells in the TDLN to be as in Table 4, with justification for the choice of these values in Appendix B.2.

**Table 4:**
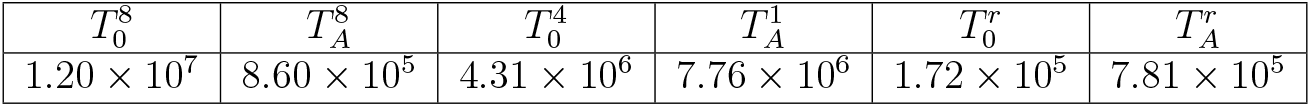
TDLN initial condition cell densities for the model. All values are in cell/cm^3^.

We also estimated the initial condition for *D*^LN^ to be 1.78 × 10^7^ cell/cm^3^, with justification for this in Appendix C.7.1.

### 3.3 Initial Conditions for DAMPs

We chose the DAMP initial conditions to be as in Table 5, with justification for the choice of these values in Appendix B.3.

**Table 5:**
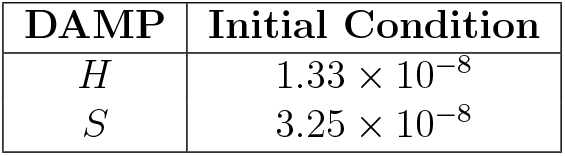
DAMP initial conditions for the model. All values are in units of g/cm^3^.

| DAMP | Initial Condition |
| --- | --- |
| $H$ | $1.33 \times 10^{-8}$ |
| $S$ | $3.25 \times 10^{-8}$ |

### 3.4 Initial Conditions for Cytokines

We chose the cytokine initial conditions to be as in Table 6, with justification for the choice of these values in Appendix B.4 and Appendix C.6.

**Table 6:** Cytokine initial conditions for the model. All values are in units of g/cm^3^.

| Cytokine | Initial Condition |
| --- | --- |
| $I_2$ | $1.87 \times 10^{-12}$ |
| $I_\gamma$ | $1.10 \times 10^{-10}$ |
| $I_\alpha$ | $1.25 \times 10^{-10}$ |
| $I_\beta$ | $9.20 \times 10^{-7}$ |
| $I_{10}$ | $1.15 \times 10^{-10}$ |

### 3.5 Initial Conditions for Immune Checkpoint-Associated Components in the TS

We chose the TS immune checkpoint-associated component initial conditions to be as in Table 7, with justification for the choice of these values is done in Appendix C.10.

**Table 7:** TS immune checkpoint-associated component initial conditions for the model. All values are in units of molec/cm^3^.

| Protein | Initial Condition |
| --- | --- |
| $P_D^{T_8}$ | $4.44 \times 10^8$ |
| $P_D^{T_1}$ | $2.07 \times 10^8$ |
| $P_D^K$ | $2.46 \times 10^8$ |
| $P_L$ | $7.40 \times 10^{12}$ |
| $Q^{T_8}$ | $6.99 \times 10^5$ |
| $Q^{T_1}$ | $3.26 \times 10^5$ |
| $Q^K$ | $3.88 \times 10^5$ |

We also set the initial condition for all pembrolizumab-associated components in the TS to be 0, as shown in Table 8.

**Table 8:** Initial conditions for pembrolizumab-associated components in the TS in the model. All values are in units of molec/cm^3^.

| Protein | Initial Condition |
| --- | --- |
| $Q_A^{T_8}$ | 0 |
| $Q_A^{T_1}$ | 0 |
| $Q_A^K$ | 0 |
| $A_1$ | 0 |

### 3.6 Initial Conditions for Immune Checkpoint-Associated Components in the TDLN

We chose the TDLN immune checkpoint-associated component initial conditions to be as in Table 9, with justification for the choice of these values in Appendix C.11.

**Table 9:** TDLN immune checkpoint-associated component initial conditions for the model. All values are in units of molec/cm^3^.

| Protein | Initial Condition |
| --- | --- |
| $P_D^{8LN}$ | $2.37 \times 10^9$ |
| $P_D^{1LN}$ | $1.59 \times 10^{10}$ |
| $P_L^{LN}$ | $3.34 \times 10^{11}$ |
| $Q^{8LN}$ | $1.69 \times 10^5$ |
| $Q^{1LN}$ | $1.13 \times 10^6$ |

We also set the initial condition for all pembrolizumab-related quantities in the TDLN to be 0, as shown in Table 10.

**Table 10:** Initial conditions for pembrolizumab-associated components in the TDLN in the model. All values are in units of molec/cm^3^.

| Protein | Initial Condition |
| --- | --- |
| $Q_A^{8LN}$ | 0 |
| $Q_A^{1LN}$ | 0 |
| $A_1^{LN}$ | 0 |

## 4 Results

We now aim to optimise pembrolizumab therapy for dnmMCRC. For simplicity, we assume that pembrolizumab is given at a constant dosage, and the spacing between consecutive pembrolizumab infusions is constant. We also assume that the patient has pembrolizumab at *t* = 0 days, and we consider a treatment regimen lasting for 96 weeks so that the time for the latest allowed infusion is *t* = 96 weeks = 672 days. Furthermore, we assume that the patient has a mass of *m* = 80 kg. In our optimisation of pembrolizumab therapy, we consider the following four objectives: tumour volume reduction (TVR), efficacy, efficiency, and toxicity.

We denote *V*_TS_(*ξ*_pembro_; *η*_pembro_, *t*) as the primary tumour volume at time *t* with treatment with pembrolizumab doses of *ξ*_pembro_ (in mg/kg) at a dosing interval of *η*_pembro_ (in weeks), omitting the *ξ*_pembro_ and *η*_pembro_ arguments in the case that no treatment is given. We define the TVR from this regimen to be

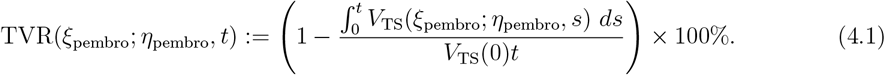

We also define the efficacy similarly as

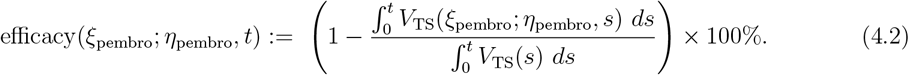

In particular, efficacy represents the extent of tumour volume shrinkage throughout its growth course in comparison to no treatment, i.e the extent of tumour growth inhibition, whereas the TVR reveals how much the tumour volume has reduced since the commencement of treatment. We see that the TVR and efficacy are linearly related, so that an increase in treatment efficacy results in increased TVR, and vice versa, via the formula

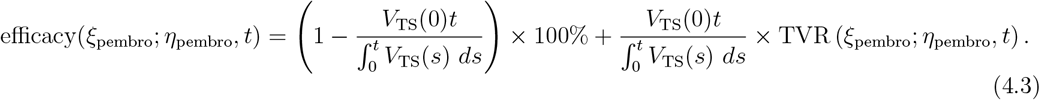

We can also consider the efficiency of the treatment regimen, with a dosing interval of *η*_pembro_ weeks and dosage *ξ*_pembro_ mg/kg given by

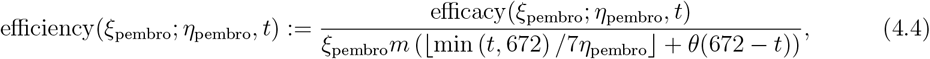

where *θ*(*t*) is the Heaviside function which equals 1 if *t* ≥ 0, and 0 otherwise. In particular, *ξ*_pembro_*m* (⌊min (*t*, 672) /7*η*_pembro_⌋ + *θ*(672 − *t*)) is the total dose of pembrolizumab administered by time *t*, recalling that no treatment is given for *t* ≥ 672 days. This corresponds to the ratio between the efficacy and the total dose of pembrolizumab administered.

Finally, we can define the toxicity of the treatment regimen, noting that large enough pembrolizumab concentrations can potentially cause hepatotoxicity and ocular toxicity [93, 94], as well as increase the probability of serious infections and malignancies. Experiments show that dosages of pembrolizumab between 0.1 mg/kg and 10 mg/kg, given every 2 weeks, is safe and tolerable [95, 96]. We thus assume that the threshold for pembrolizumab toxicity is 10 mg/kg every 2 weeks, with higher doses being deemed toxic. To rigorise this notion, we define the toxicity of the treatment regimen, with a dosing interval of *η*_pembro_ weeks and dosage *ξ*_pembro_ mg/kg, as

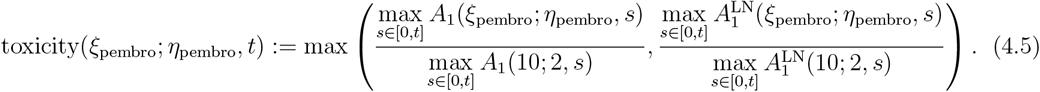

In particular, *A*_1_(*ξ*_pembro_; *η*_pembro_, *s*) and 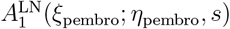 denote the concentrations of *A*_1_ and 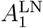 at time *s*, with pembrolizumab doses of *ξ*_pembro_ at a dosing interval of *η*_pembro_, respectively. In particular, the toxicity quantifies the ratio of the maximum pembrolizumab concentrations from the regimen to those of a 10 mg/kg dose given every 2 weeks, taking the highest value of this ratio between the TDLN and TS. A toxicity greater than 1 indicates a toxic and unsafe regimen, whereas a toxicity of 1 or less signifies a non-toxic and safe regimen, with lower toxicity values corresponding to safer treatments.

Furthermore, we use the two FDA-approved pembrolizumab regimens for the first-line treatment of mMCRC in adults as a benchmark for comparison [97]:

- <u>Treatment 1</u>: 200 mg of pembrolizumab administered by intravenous infusion over a duration of 30 minutes every 3 weeks until disease progression or unacceptable toxicity.
- <u>Treatment 2</u>: 400 mg of pembrolizumab administered by intravenous infusion over a duration of 30 minutes every 6 weeks until disease progression or unacceptable toxicity.

These correspond to the following parameter values in the model:

- Treatment 1: *ξ*_*j*_ = 200 mg, *t*_*j*_ = 21(*j −* 1), *n* = 32, *ξ*_pembro_ = 2.5 mg/kg, *η*_pembro_ = 3 weeks,
- Treatment 2: *ξ*_*j*_ = 400 mg, *t*_*j*_ = 42(*j −* 1), *n* = 16, *ξ*_pembro_ = 5 mg/kg, *η*_pembro_ = 6 weeks.

Denoting the dosing interval of pembrolizumab as *η*_pembro_, we perform a sweep across the space *η*_pembro_ ∈ {1, 2, 3, 4, 6, 8, 12} weeks. These values are integer factors of 96 weeks, and each *η*_pembro_ corresponds to a distinct number of doses administered. This approach ensures practicality whilst preventing any artefacts that could occur from selecting a treatment regimen that ends at a fixed time of 96 weeks. Taking practicality constraints into account, we consider linearly spaced dosages in the domain *ξ*_pembro_ ∈ [0.1, 10] mg/kg, with a spacing of 0.0125 mg/kg. This corresponds to *ξ*_*j*_ ∈ [0.1*m*, 10*m*] mg = [8, 800] mg with an increment of 1 mg.

We can determine the optimal pembrolizumab therapy by considering the regimen that achieves an acceptable efficacy at 96 weeks whilst maximising treatment efficiency as much as possible and ensuring a toxicity of less than 1. The efficacies of Treatment 1 and Treatment 2 at 96 weeks were calculated to be 62.14%. As such, to ensure that the TVR of the optimal treatment is comparable to current FDA-approved pembrolizumab regimens, we consider threshold efficacies of 62.14%, 62%, 61%, and 60%. We also consider constraints due to practicality, so that *ξ*_pembro_ is an integer multiple of 0.1*m* mg/kg, corresponding to an integer multiple of 8 mg, leaving the domain for *η*_pembro_ unchanged. Denoting the and *η* space of (*ξ*_pembro_, *η*_pembro_) pairs that satisfy these criteria as *𝒮*^prac^, the optimal pembrolizumab dosing and spacing, denoted 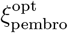 and 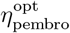, respectively, for a given threshold efficacy ℰ_thresh_, satisfy

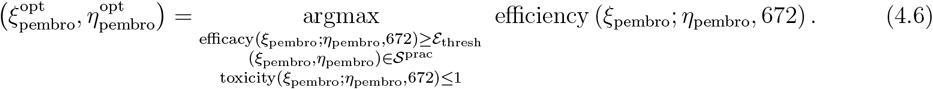

Solutions of (4.6) with the previously given threshold efficacies compared to Treatments 1 and 2 are shown in Table 11.

**Table 11:** Comparison of 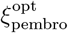, dosage, spacing (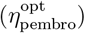), TVR, efficacy, efficiency, and toxicity between FDA-approved regimens and optimal treatment regimens for various ℰ_thresh_, assuming a patient mass of 80 kg. Tx No. denotes the treatment number, with FDA-approved therapies labelled as Treatments 1 and 2, and optimal regimens labelled as Treatments 3–6.

| | Tx Num. | $\mathcal{E}_{\text{thresh}}$ (%) | $\eta_{\text{pembro}}^{\text{opt}}$ (mg/kg) | Dosage (mg) | Spacing (weeks) | TVR (%) | Efficacy (%) | Efficiency (%/mg) | Toxicity |
| --- | --- | --- | --- | --- | --- | --- | --- | --- | --- |
| FDA | 1 | — | — | 200 | 3 | 34.07 | 62.14 | $9.71 \times 10^{-3}$ | $1.83 \times 10^{-1}$ |
| | 2 | — | — | 400 | 6 | 34.08 | 62.14 | $9.71 \times 10^{-3}$ | $2.37 \times 10^{-1}$ |
| Optimal | 3 | 62.14 | 3.3 | 264 | 4 | 34.08 | 62.14 | $9.81 \times 10^{-3}$ | $1.98 \times 10^{-1}$ |
| | 4 | 62 | 3.0 | 240 | 4 | 33.87 | 62.02 | $1.08 \times 10^{-2}$ | $1.80 \times 10^{-1}$ |
| | 5 | 61 | 1.7 | 136 | 4 | 32.17 | 61.05 | $1.87 \times 10^{-2}$ | $1.02 \times 10^{-1}$ |
| | 6 | 60 | 1.2 | 96 | 4 | 30.66 | 60.18 | $2.61 \times 10^{-2}$ | $7.21 \times 10^{-2}$ |

Heatmaps of TVR, efficacy, efficiency, and toxicity at *t* = 96 weeks for various *η*_pembro_ and *ξ*_pembro_ values are shown in Figure 7. All simulations were done in MATLAB using the dde23 solver with the initial conditions stated in Section 3.

**Figure 7:**
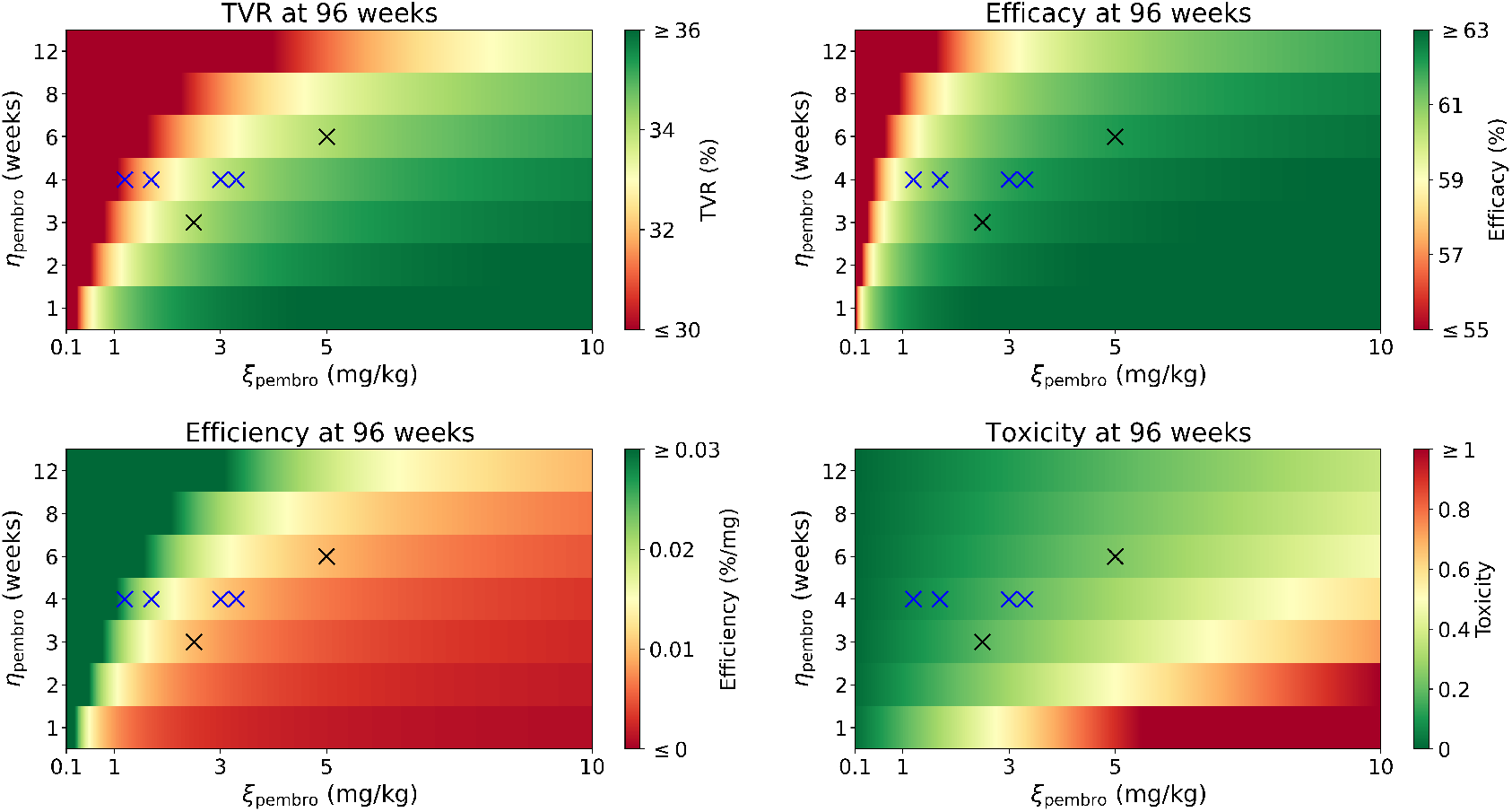
TVR (top left), efficacy (top right), efficiency (bottom left), and toxicity (bottom right) at 96 weeks for *η*_pembro_ ∈ {1, 2, 3, 4, 6, 8, 12} weeks. We sweep across *ξ*_pembro_ ∈ [0.1, 10] mg/kg with an increment of 0.0125 mg/kg. The FDA-approved regimens (Treatments 1 and 2) for mMCRC are shown in black, and the optimal regimens (Treatments 3–6) are shown in blue.

Time traces of TVR, efficacy, efficiency, and toxicity for Treatments 1–6 are shown in Figure 8.

**Figure 8:**
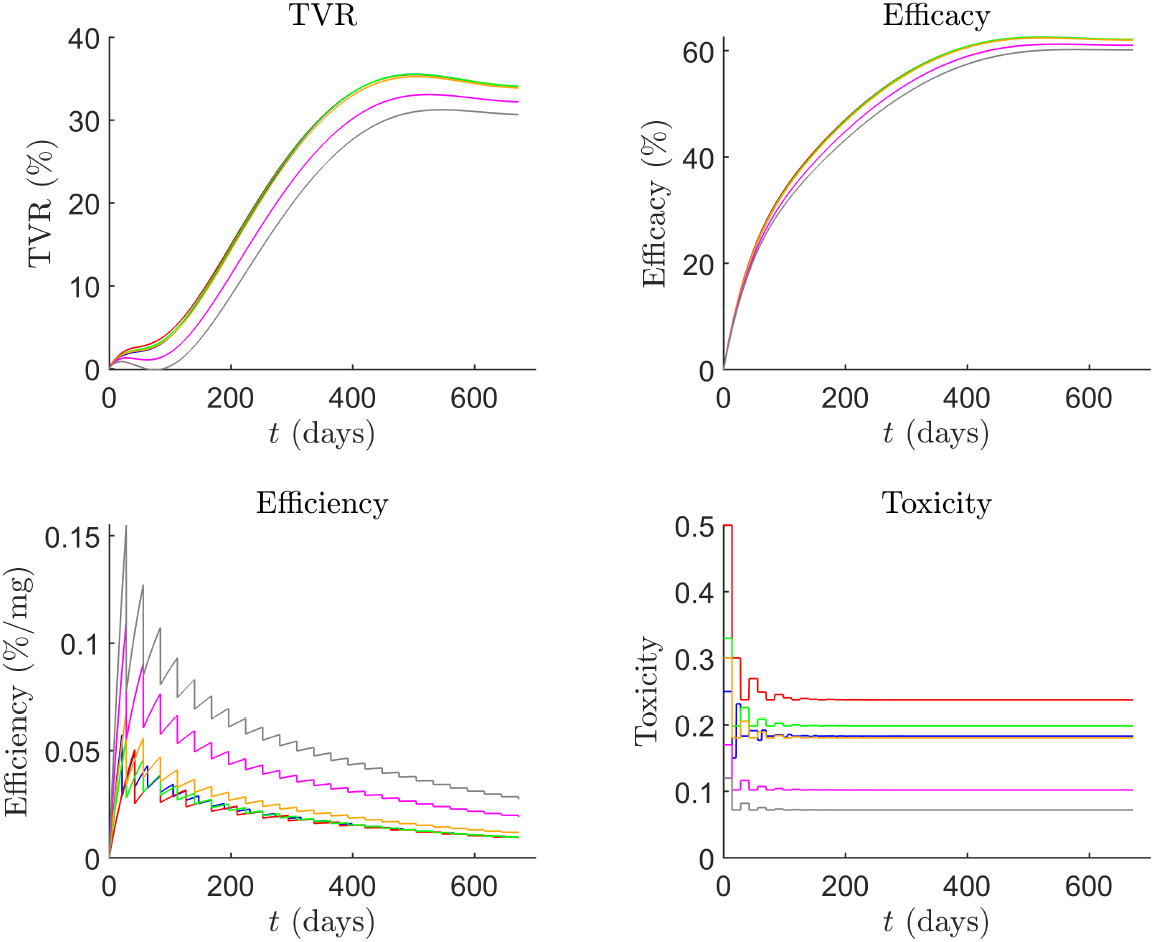
Time traces of TVR (top left), efficacy (top right), efficiency (bottom left), and toxicity (bottom right) for Treatments 1–6 in blue, red, green, orange, magenta, and grey, respectively.

Time traces for the primary tumour volume, *V*_TS_, with Treatments 1–6 compared to no treatment are shown in Figure 9.

**Figure 9:**
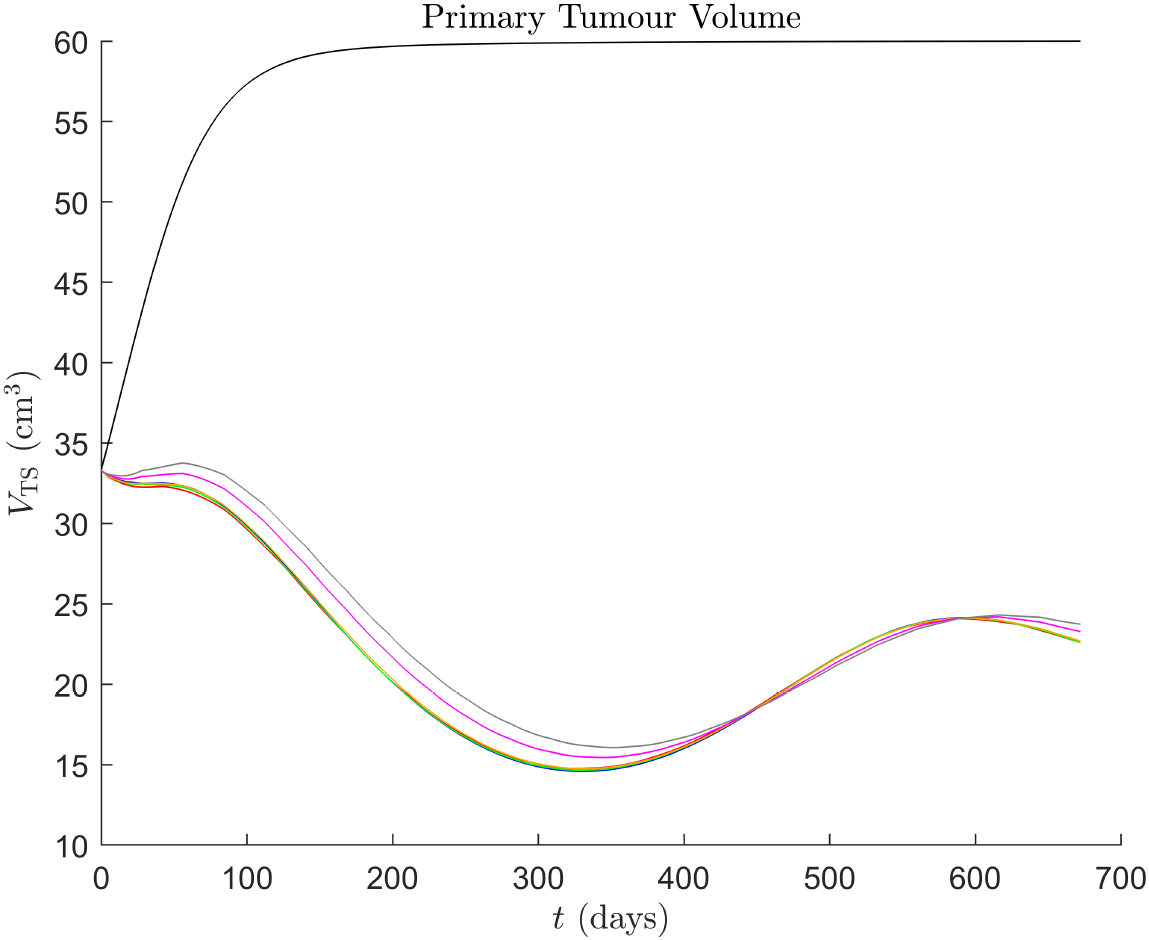
Time traces of *V*_TS_ up to 96 weeks from commencement, with no treatment in black, and Treatments 1–6 in blue, red, green, orange, magenta, and grey, respectively.

We can also compare the effects of optimal pembrolizumab therapies and FDA-approved regimens to those of no treatment on the TME, with time traces of model variables shown in Figure 10 and average immune cell and cytokine concentrations at 96 weeks shown in Table 12.

**Table 12:**
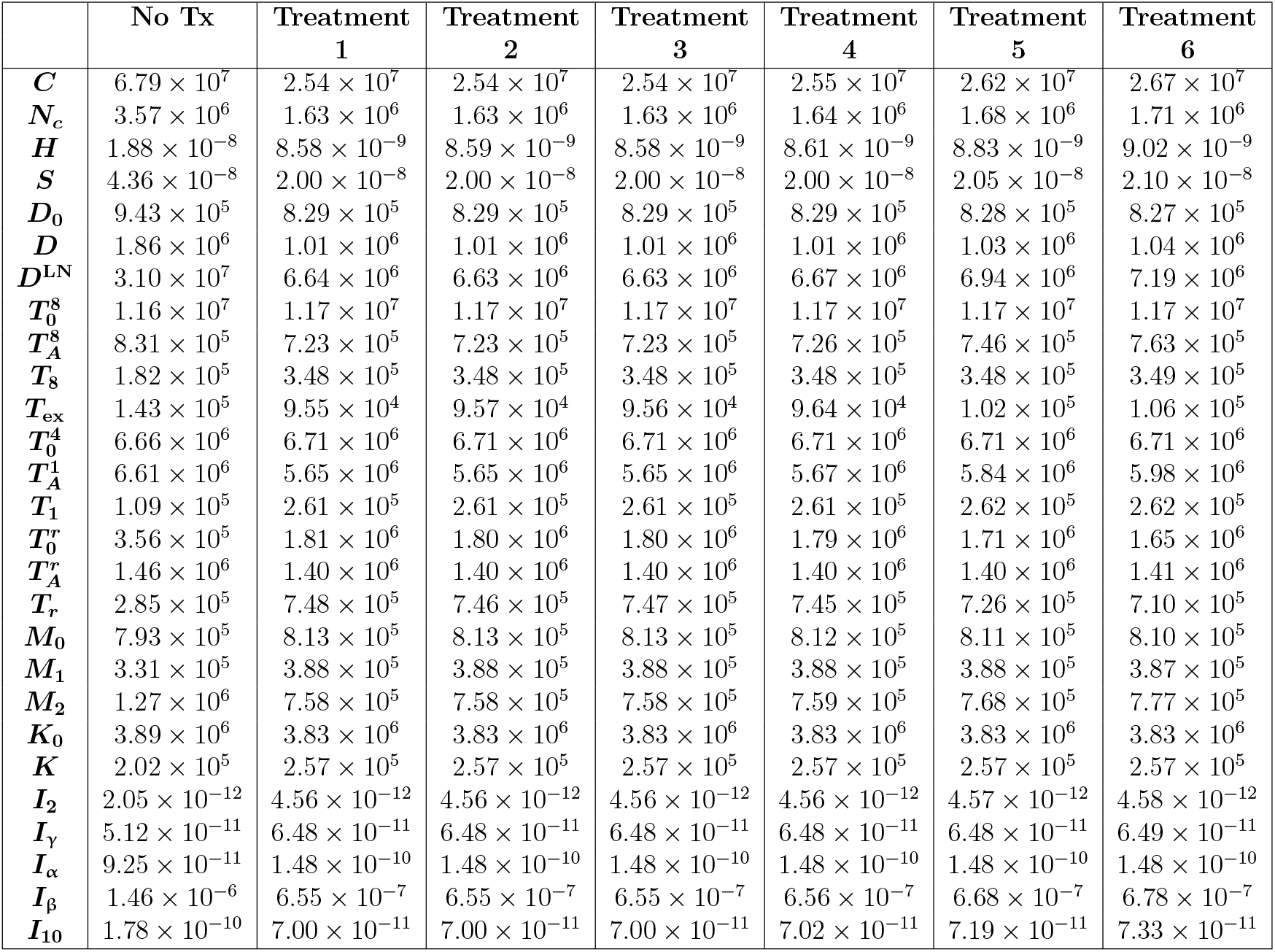
Comparison of average immune cell and cytokine concentrations at 96 weeks between no treatment and Treatments 1–6. Units of variables are as in Table 1.

|  | No Tx | Treatment<br>1 | Treatment<br>2 | Treatment<br>3 | Treatment<br>4 | Treatment<br>5 | Treatment<br>6 |
| --- | --- | --- | --- | --- | --- | --- | --- |
| $C$ | $6.79 \times 10^7$ | $2.54 \times 10^7$ | $2.54 \times 10^7$ | $2.54 \times 10^7$ | $2.55 \times 10^7$ | $2.62 \times 10^7$ | $2.67 \times 10^7$ |
| $N_c$ | $3.57 \times 10^6$ | $1.63 \times 10^6$ | $1.63 \times 10^6$ | $1.63 \times 10^6$ | $1.64 \times 10^6$ | $1.68 \times 10^6$ | $1.71 \times 10^6$ |
| $H$ | $1.88 \times 10^{-8}$ | $8.58 \times 10^{-9}$ | $8.59 \times 10^{-9}$ | $8.58 \times 10^{-9}$ | $8.61 \times 10^{-9}$ | $8.83 \times 10^{-9}$ | $9.02 \times 10^{-9}$ |
| $S$ | $4.36 \times 10^{-8}$ | $2.00 \times 10^{-8}$ | $2.00 \times 10^{-8}$ | $2.00 \times 10^{-8}$ | $2.00 \times 10^{-8}$ | $2.05 \times 10^{-8}$ | $2.10 \times 10^{-8}$ |
| $D_0$ | $9.43 \times 10^5$ | $8.29 \times 10^5$ | $8.29 \times 10^5$ | $8.29 \times 10^5$ | $8.29 \times 10^5$ | $8.28 \times 10^5$ | $8.27 \times 10^5$ |
| $D$ | $1.86 \times 10^6$ | $1.01 \times 10^6$ | $1.01 \times 10^6$ | $1.01 \times 10^6$ | $1.01 \times 10^6$ | $1.03 \times 10^6$ | $1.04 \times 10^6$ |
| $D^{\text{LN}}$ | $3.10 \times 10^7$ | $6.64 \times 10^6$ | $6.63 \times 10^6$ | $6.63 \times 10^6$ | $6.67 \times 10^6$ | $6.94 \times 10^6$ | $7.19 \times 10^6$ |
| $T_0^8$ | $1.16 \times 10^7$ | $1.17 \times 10^7$ | $1.17 \times 10^7$ | $1.17 \times 10^7$ | $1.17 \times 10^7$ | $1.17 \times 10^7$ | $1.17 \times 10^7$ |
| $T_A^8$ | $8.31 \times 10^5$ | $7.23 \times 10^5$ | $7.23 \times 10^5$ | $7.23 \times 10^5$ | $7.26 \times 10^5$ | $7.46 \times 10^5$ | $7.63 \times 10^5$ |
| $T_8$ | $1.82 \times 10^5$ | $3.48 \times 10^5$ | $3.48 \times 10^5$ | $3.48 \times 10^5$ | $3.48 \times 10^5$ | $3.48 \times 10^5$ | $3.49 \times 10^5$ |
| $T_{\text{ex}}$ | $1.43 \times 10^5$ | $9.55 \times 10^4$ | $9.57 \times 10^4$ | $9.56 \times 10^4$ | $9.64 \times 10^4$ | $1.02 \times 10^5$ | $1.06 \times 10^5$ |
| $T_0^4$ | $6.66 \times 10^6$ | $6.71 \times 10^6$ | $6.71 \times 10^6$ | $6.71 \times 10^6$ | $6.71 \times 10^6$ | $6.71 \times 10^6$ | $6.71 \times 10^6$ |
| $T_A^1$ | $6.61 \times 10^6$ | $5.65 \times 10^6$ | $5.65 \times 10^6$ | $5.65 \times 10^6$ | $5.67 \times 10^6$ | $5.84 \times 10^6$ | $5.98 \times 10^6$ |
| $T_1$ | $1.09 \times 10^5$ | $2.61 \times 10^5$ | $2.61 \times 10^5$ | $2.61 \times 10^5$ | $2.61 \times 10^5$ | $2.62 \times 10^5$ | $2.62 \times 10^5$ |
| $T_0^r$ | $3.56 \times 10^5$ | $1.81 \times 10^6$ | $1.80 \times 10^6$ | $1.80 \times 10^6$ | $1.79 \times 10^6$ | $1.71 \times 10^6$ | $1.65 \times 10^6$ |
| $T_A^r$ | $1.46 \times 10^6$ | $1.40 \times 10^6$ | $1.40 \times 10^6$ | $1.40 \times 10^6$ | $1.40 \times 10^6$ | $1.40 \times 10^6$ | $1.41 \times 10^6$ |
| $T_r$ | $2.85 \times 10^5$ | $7.48 \times 10^5$ | $7.46 \times 10^5$ | $7.47 \times 10^5$ | $7.45 \times 10^5$ | $7.26 \times 10^5$ | $7.10 \times 10^5$ |
| $M_0$ | $7.93 \times 10^5$ | $8.13 \times 10^5$ | $8.13 \times 10^5$ | $8.13 \times 10^5$ | $8.12 \times 10^5$ | $8.11 \times 10^5$ | $8.10 \times 10^5$ |
| $M_1$ | $3.31 \times 10^5$ | $3.88 \times 10^5$ | $3.88 \times 10^5$ | $3.88 \times 10^5$ | $3.88 \times 10^5$ | $3.88 \times 10^5$ | $3.87 \times 10^5$ |
| $M_2$ | $1.27 \times 10^6$ | $7.58 \times 10^5$ | $7.58 \times 10^5$ | $7.58 \times 10^5$ | $7.59 \times 10^5$ | $7.68 \times 10^5$ | $7.77 \times 10^5$ |
| $K_0$ | $3.89 \times 10^6$ | $3.83 \times 10^6$ | $3.83 \times 10^6$ | $3.83 \times 10^6$ | $3.83 \times 10^6$ | $3.83 \times 10^6$ | $3.83 \times 10^6$ |
| $K$ | $2.02 \times 10^5$ | $2.57 \times 10^5$ | $2.57 \times 10^5$ | $2.57 \times 10^5$ | $2.57 \times 10^5$ | $2.57 \times 10^5$ | $2.57 \times 10^5$ |
| $I_2$ | $2.05 \times 10^{-12}$ | $4.56 \times 10^{-12}$ | $4.56 \times 10^{-12}$ | $4.56 \times 10^{-12}$ | $4.56 \times 10^{-12}$ | $4.57 \times 10^{-12}$ | $4.58 \times 10^{-12}$ |
| $I_\gamma$ | $5.12 \times 10^{-11}$ | $6.48 \times 10^{-11}$ | $6.48 \times 10^{-11}$ | $6.48 \times 10^{-11}$ | $6.48 \times 10^{-11}$ | $6.48 \times 10^{-11}$ | $6.49 \times 10^{-11}$ |
| $I_\alpha$ | $9.25 \times 10^{-11}$ | $1.48 \times 10^{-10}$ | $1.48 \times 10^{-10}$ | $1.48 \times 10^{-10}$ | $1.48 \times 10^{-10}$ | $1.48 \times 10^{-10}$ | $1.48 \times 10^{-10}$ |
| $I_\beta$ | $1.46 \times 10^{-6}$ | $6.55 \times 10^{-7}$ | $6.55 \times 10^{-7}$ | $6.55 \times 10^{-7}$ | $6.56 \times 10^{-7}$ | $6.68 \times 10^{-7}$ | $6.78 \times 10^{-7}$ |
| $I_{10}$ | $1.78 \times 10^{-10}$ | $7.00 \times 10^{-11}$ | $7.00 \times 10^{-11}$ | $7.00 \times 10^{-11}$ | $7.02 \times 10^{-11}$ | $7.19 \times 10^{-11}$ | $7.33 \times 10^{-11}$ |

**Figure 10:**
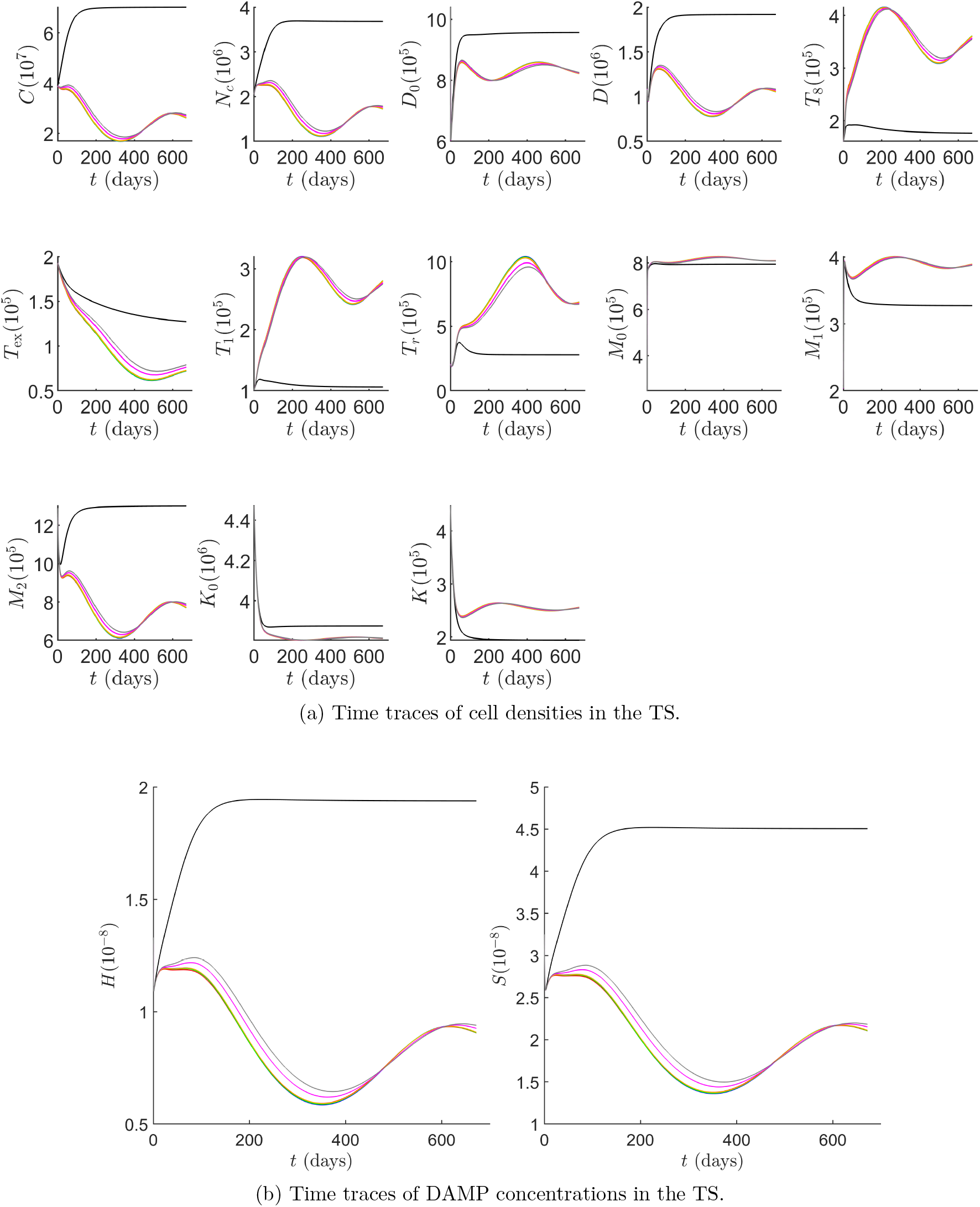
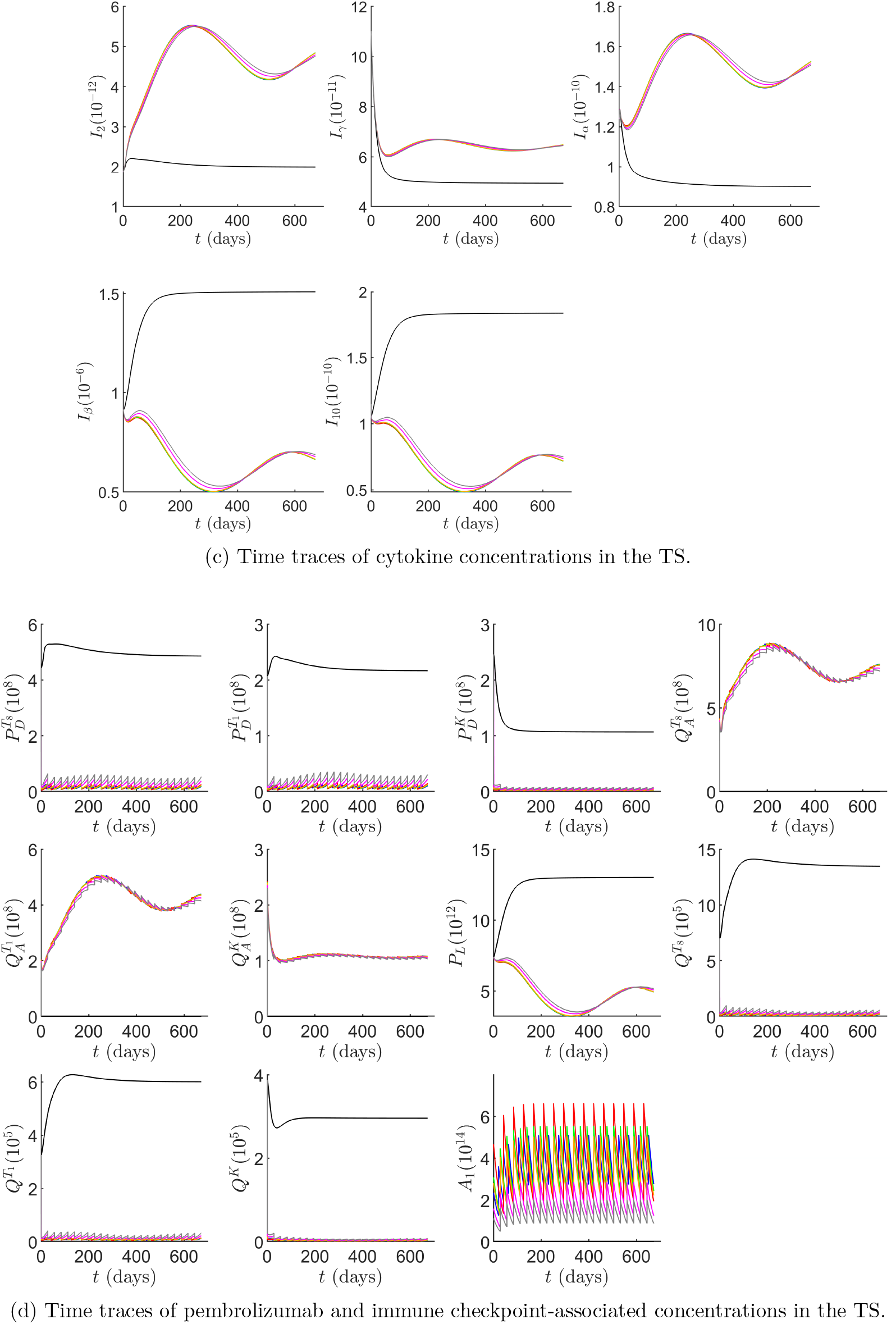
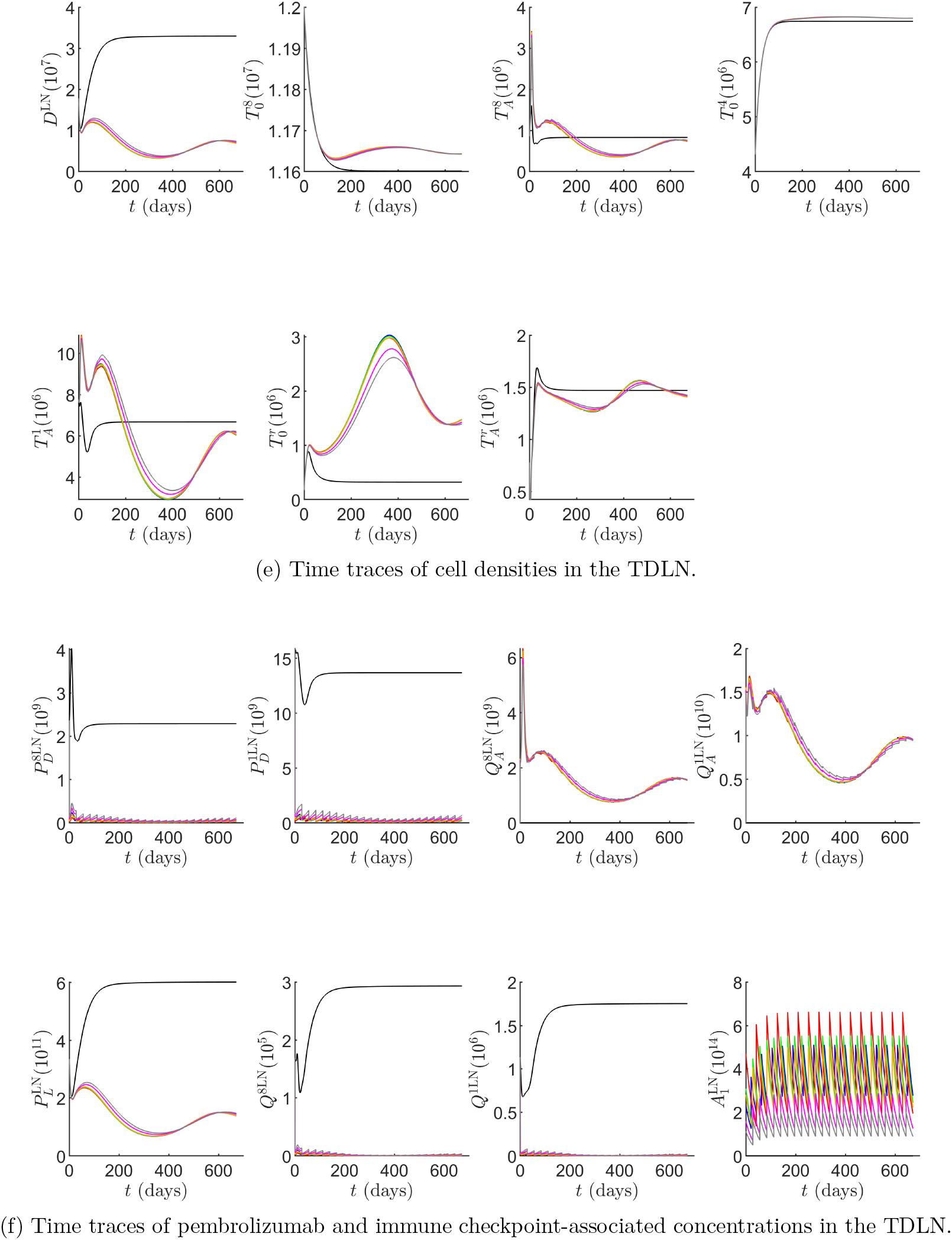
Time traces of variables in the model, with the units of the variables as in Table 1. Time traces with no treatment are in black, and Treatments 1–6 in blue, red, green, orange, magenta, and grey, respectively.

## 5 Discussion

We proceed to analyse the results from Figure 7, Figure 8, Figure 9, Figure 10, and Table 12 will be discussed in detail, and compare them to findings from laMCRC reported in [36]. We first analyse Figure 9, Figure 10, and Table 12 for the FDA-approved treatment regimens, Treatment 1 and Treatment 2.

We can see from Figure 9 that the FDA-approved treatments are effective during the first half of the treatment period at eradicating cancer cells, with the primary tumour volume reaching a minimum of approximately 14.59 cm^3^ at 330 days. This corresponds to a 56.1% decrease from the initial volume and a 75.6% reduction compared to the tumour volume without treatment at that time. We note that it takes a couple of months of pembrolizumab therapy for consistent tumour reductions to be observed, in agreement with experimental findings for mMCRC [11]. However, the treatment becomes insufficiently effective thereafter, leading to an increase in *V*_TS_, which peaks at 24.14 cm^3^ at 591 days before beginning to decline again, albeit at a slower rate. This is in contrast to the case of laMCRC, where treatment leads to a rapid and sustained decrease in total cancer concentration, and equivalently the tumour volume, even after treatment ends at 12 weeks, with a reduction of more than 85% by 18 weeks.

To justify this behaviour, we first examine the time traces of the model components shown in Figure 10, which allow us to identify key factors that contribute to maximising cancer reduction. One immediate point of note is that the TME is significantly more immunogenic and pro-inflammatory in laMCRC, with the initial concentrations of pro-inflammatory effector CD8+ T cells, effector Th1 cells, activated NK cells and M1 macrophages being 2.43 × 10^5^ cell/cm^3^, 1.04 × 10^5^ cell/cm^3^, 5.20 ×10^6^ cell/cm^3^, and 6.61 × 10^5^ cell/cm^3^, respectively, compared to 1.61 ×10^5^ cell/cm^3^, 1.01 ×10^5^ cell/cm^3^, 4.47 ×10^5^ cell/cm^3^, and 2.09 ×10^5^ cell/cm^3^ in dnmMCRC. Similarly, the concentrations of antiinflammatory cells, particularly viable cancer cells, effector Tregs, and M2 macrophages, are initially 1.79 × 10^5^ cell/cm^3^, 2.12 × 10^5^ cell/cm^3^, and 1.23 × 10^6^ cell/cm^3^ in laMCRC, respectively, compared to the generally larger values of 3.90 × 10^7^ cell/cm^3^, 2.02 × 10^5^ cell/cm^3^, and 1.29 × 10^6^ cell/cm^3^ in dnmMCRC. As such, in dnmMCRC, the TME is so strongly anti-inflammatory that a greater degree of stimulation is required to shift the system toward the more pro-inflammatory state as observed in laMCRC, with this transition inherently requiring more time.

Another important observation is that pembrolizumab therapy leads to a substantial increase in the concentration of activated and effector pro-inflammatory immune cells within the TS, compared to without treatment. Specifically, the concentrations of effector CD8+ T cells and effector Th1 cells increase significantly and monotonically, reaching peak values of 4.16 × 10^5^ cell/cm^3^ and 3.20 × 10^5^ cell/cm^3^ at 214 and 251 days, respectively, compared to 1.84 × 10^5^ cell/cm^3^ and 1.08 × 10^5^ cell/cm^3^ without treatment. Similarly, after an initial transient decrease, the concentrations of pro-inflammatory activated NK cells and M1 macrophages increase, reaching peaks of 2.64 × 10^5^ cell/cm^3^ and 4.01 × 10^5^ cell/cm^3^, respectively, at 245 and 269 days. This trend is reflected in the concentrations of pro-inflammatory cytokines IL-2, IFN-γ, and TNF which, after transient decreases, rise to 5.53 × 10^−12^ molec/cm^3^, 6.71 × 10^−11^ molec/cm^3^, and 1.67 × 10^−10^ molec/cm^3^ by 241, 224, and 239 days, respectively, compared to 2.05 × 10^−12^ molec/cm^3^, 4.97 × 10^−11^ molec/cm^3^ and 9.13 × 10^−11^ molec/cm^3^ in the absence of therapy. However, following their peaks, all these concentrations decline markedly. The concentrations of effector CD8+ T cells, effector Th1 cells, activated NK cells, M1 macrophages, IL-2, IFN-γ, and TNF reach their minimum values at approximately 498, 515, 525, 537, 510, 496, and 510 days, respectively, with corresponding concentrations of 3.09 × 10^5^ cell/cm^3^, 2.42 × 10^5^ cell/cm^3^, 2.49 × 10^5^ cell/cm^3^, 3.83 × 10^5^ cell/cm^3^, 4.17 × 10^−12^ molec/cm^3^, 6.24 × 10^−11^ molec/cm^3^, and 1.39 × 10^−10^ molec/cm^3^, before beginning to increase again. Nonetheless, the concentration of pro-inflammatory cells and cytokines in the TS remains significantly higher than without treatment, with the average concentrations of effector CD8+ T cells, effector Th1 cells, activated NK cells, M1 macrophages, IL-2, IFN-γ, and TNF under treatment being 3.48 × 10^5^ cell/cm^3^, 2.61 × 10^5^ cell/cm^3^, 2.57 × 10^5^ cell/cm^3^, 3.88 × 10^5^ cell/cm^3^, 4.56 × 10^−12^ molec/cm^3^, 6.48 × 10^−11^ molec/cm^3^, and 1.48 × 10^−10^ molec/cm^3^, respectively, all notably higher than their counterparts without treatment, which are 1.82× 10^5^ cell/cm^3^, 1.09 × 10^5^ cell/cm^3^, 2.02 × 10^5^ cell/cm^3^, 3.31 × 10^5^ cell/cm^3^, 2.05 × 10^−12^ molec/cm^3^, 5.12 × 10^−11^ molec/cm^3^, and 9.25 × 10^−11^ molec/cm^3^, without, using data from Table 12.

This behaviour is mirrored in the anti-inflammatory cells and cytokines within the TS, including viable cancer cells, M2 macrophages, TGF-β, and IL-10, whose concentrations decrease significantly up to approximately 329, 324, 317, and 327 days under treatment, respectively, reaching values of 1.68 × 10^7^ cell/cm^3^, 6.11 × 10^5^ cell/cm^3^, 5.00 × 10^−7^ molec/cm^3^, and 4.84 × 10^−11^ molec/cm^3^—compared to 7.01 × 10^7^ cell/cm^3^, 1.30 × 10^6^ cell/cm^3^, 1.51 × 10^−6^ molec/cm^3^, and 1.84 × 10^−10^ molec/cm^3^ without treatment. Following this decline, their concentrations rise again, peaking at around 589 days with corresponding concentrations of 2.80 × 10^7^ cell/cm^3^, 8.01 × 10^5^ cell/cm^3^, 7.02 × 10^−7^ molec/cm^3^, and 7.64 × 10^−11^ molec/cm^3^, before decreasing once more. Nonetheless, the concentrations of these components remains significantly lower than without treatment, with the average concentrations of viable cancer cells, M2 macrophages, TGF-β, and IL-10 under treatment being 2.54 × 10^7^ cell/cm^3^, 7.58 × 10^5^ cell/cm^3^, 6.55 × 10^−7^ molec/cm^3^, and 7.00 × 10^−11^ molec/cm^3^, respectively, which are all lower than their counterparts without treatment, which are 6.79 × 10^7^ cell/cm^3^, 1.27 × 10^6^ cell/cm^3^, 1.46 × 10^−6^ molec/cm^3^, and 1.78 × 10^−10^ molec/cm^3^, using data from Table 12.

These oscillations in concentration are perpetuated by several positive feedback loops within the TME. Firstly, increases in effector CD8+ and Th1 cell concentration lead to an increased concentration of IL-2, a key growth factor for these cells and an activator of resting NK cells. This, in turn, enhances IL-2 production, further stimulating effector CD8+ and Th1 cell proliferation, reinforcing the loop. Furthermore, elevated levels of pro-inflammatory cells enhance tumour cell lysis and increase production of pro-inflammatory cytokines, such as TNF and IFN-γ, which drive cancer cell necrosis and macrophage polarisation into the pro-inflammatory M1 phenotype, resulting in another positive feedback loop. Moreover, decreased concentrations of anti-inflammatory cells lead to diminished production of anti-inflammatory cytokines, which decreases polarisation of macrophages to the M2 phenotype, further decreasing anti-inflammatory cytokine production and reinforcing a positive feedback loop. A particularly potent example of this occurs with respect to TGF-β: as the levels of anti-inflammatory cells decline, TGF-β concentrations decrease, which reduces the inhibition of cancer cell lysis by effector CD8+ T cells and activated NK cells, diminishes suppression of NK cell activation, and reduces M2 macrophage polarisation. This further lowers the number of immunosuppressive cells, which perpetuates the decline in TGF-β concentration and amplifies the anti-tumour response. Furthermore, another important positive feedback loop involves PD-L1 concentrations in the TS. As anti-inflammatory cell concentrations, in particular viable cancer cells and M2 macrophages, decrease, there is a decrease in the total PD-L1 concentration in the TS, resulting in reduced PD-1/PD-L1 complex formation, and enhanced lysis of cancer cells by effector CD8+ T cells and activated NK cells. This further lowers cancer cell concentrations and, consequently, PD-L1 concentration. However, these feedback mechanisms are bidirectional: a decline in pro-inflammatory cell concentrations leads to reduced levels of pro-inflammatory cytokines, and vice versa. Similarly, increases in anti-inflammatory cell concentrations promote greater production of anti-inflammatory cytokines, and vice versa.

We now aim to provide an explanation for the sudden change in dynamics observed, focusing on three key aspects: DC maturation and migration to the TDLN, T cell proliferation and activation within the TDLN, and Treg hyperproliferation at the TS. Up to 330 days, as treatment progresses, tumour burden decreases, resulting in a decrease in the magnitude of necrotic cancer cells and, consequently, lower DAMP release and reduced DC maturation. This leads to a substantial decline in the concentration of mature DCs, reaching minima of 7.78 × 10^5^ cell/cm^3^ and 3.25 × 10^6^ cell/cm^3^ in the TS and TDLN, respectively, compared to 1.92 × 10^6^ cell/cm^3^ and 3.29 × 10^7^ cell/cm^3^ without treatment at the same time point. However, the concentration of mature DCs peaks to 1.09 × 10^6^ cell/cm^3^ in the TS and 7.52 × 10^6^ cell/cm^3^ in the TDLN at approximately 600 days, before declining again, closely mirroring the behaviour of anti-inflammatory cell populations.

Decreased DCs in the TDLN lead to decreased activation and proliferation of T cells in the TDLN; however, this is not the only contributing factor. Another important factor is the concentration of effector Tregs in the TDLN, which directly inhibits T cell proliferation and activation. Although their concentration initially decreases compared to without treatment, the reduction is insufficient to relieve this inhibition. Specifically, the effector Treg population in the TDLN exhibits a pattern similar to other anti-inflammatory cells, showing a pronounced dip to 1.27 × 10^6^ cell/cm^3^ at 268 days, followed by a peak of 1.57 × 10^6^ cell/cm^3^ at 468 days, before declining again, compared to a steady-state value of 1.47 × 10^6^ cell/cm^3^ without treatment. This, in conjunction with the decreased concentration of mature DCs in the TDLN, explains why the concentrations of effector CD8+ T cells and effector Th1 cells, after an initial transient phase, peak at 73 and 94 days, respectively, with concentrations reaching 1.22 × 10^6^ cell/cm^3^ and 9.50 × 10^6^ cell/cm^3^. These concentrations then fall to their minimum at 371 and 382 days, respectively, with values of 3.56 × 10^5^ cell/cm^3^ and 2.92 × 10^6^ cell/cm^3^, respectively. Notably, after 173 and 186 days, respectively, the concentrations of these effector cells become less than without treatment, leading to reduced cancer cell killing and decreased concentrations of proinflammatory cells in the TS. This initiates a cascade of effects that contribute to the later observed change in dynamics at 330 days, triggering the feedback loops described earlier.

This leads into the final critical aspect, which causes the treatment to fail: the extensive hyperproliferation of effector Tregs in the TS as a consequence of prolonged treatment. This population swells to a peak concentration of 1.04 × 10^6^ cell/cm^3^ at 391 days, far exceeding the value of 2.78 × 10^5^ cell/cm^3^ observed without treatment, before subsequently declining. This 3.74-fold increase results in strong inhibition of IL-2-mediated pro-inflammatory T cell growth, reduced IFN-γ production by T cells, and increased secretion of TGF-β, thereby exacerbating the previously mentioned TGF-β-driven feedback loop. Clinically, similar phenomena have been observed in cases of hyperprogressive disease following ICI therapy [98] and could represent a potential mechanism of acquired resistance to ICI therapy [99], both of which have been associated with treatment failure. However, given the significantly elevated effector CD8+ T cell and Th1 cell populations observed in the TS following immunotherapy, the hyperproliferation of Tregs is not entirely unexpected. A key distinction between this phenomenon in laMCRC and dnmMCRC is that Tregs may act as a regulatory mechanism to prevent excessive immune activation once the tumour burden is sufficiently reduced. Nevertheless, in dnmMCRC, this regulatory mechanism is triggered prematurely, resulting in a dampened immune response that ultimately fails to achieve complete cancer eradication.

This motivates the introduction of an additional immunotherapeutic agent aimed at depleting Tregs— an objective fulfilled by anti-CTLA-4 antibodies, which target the CTLA-4 immune checkpoint. A successful example of this approach is seen in the extended CheckMate 142 study, which evaluated the efficacy of combination treatment with nivolumab, a fully human IgG4 PD-1 antibody, and ipilimumab, a CTLA-4 antibody, for pretreated mMCRC [100]. In this study, nivolumab was administered at 3 mg/kg every three weeks, while ipilimumab was given at 1 mg/kg every three weeks for the first four doses, followed by maintenance therapy with nivolumab (3 mg/kg) every two weeks. Tumour burden reduction from baseline was observed in 77% of patients, and the 9-month and 12-month overall survival rates were 87% (95% CI 80.0–92.2%) and 85% (95% CI 77.0-90.2%), respectively [101]. These results prompted the FDA to approve combination nivolumab and ipilimumab therapy for pretreated mMCRC on July 10, 2018.

However, ipilimumab is not the only anti-CTLA-4 antibody, with promising results being reported from a phase I trial evaluating botensilimab, a fragment crystallisable (Fc)-enhanced anti-CTLA-4 antibody, with and without balstilimab, a fully human PD-1 antibody, in metastatic relapsed/refractory MSS CRC [102]. In particular, botensilimab has been demonstrated to be more potent in depleting Tregs than ipilimumab [103], largely due to enhanced antibody-dependent cellular cytotoxicity and phagocytosis mechanisms. The optimal timing, dosage, and scheduling of combination PD-1 and CTLA-4 inhibitor therapies remain important areas for future investigation. It is important to note, however, that prolonged use of anti-CTLA-4 antibodies carries a risk of autoimmune and immune-related adverse events [104], which must be carefully balanced against therapeutic benefit.

Another key observation is the substantial decrease in exhausted CD8+ T cells with pembrolizumab treatment. The average concentration of exhausted CD8+ T cells across the treatment period is 9.56 × 10^4^ cell/cm^3^, which is more than 33% lower than the 1.43 × 10^5^ cell/cm^3^ observed without treatment. We thus see that the concentration of exhausted CD8+ T cells plays a major role in treatment efficacy, as reducing the population of exhausted T cells leads to an increased concentration of effector cytotoxic CD8+ T cells, thereby enhancing cancer cell eradication.

We can also analyse the impact of therapy on the concentration of PD-1, PD-1/PD-L1, and PD-1/PD-L1 complex in the TS and the TDLN. As expected, pembrolizumab therapy significantly reduces the concentration of unbound PD-1 receptors on PD-1-expressing cells in both the TS and TDLN, with reductions of approximately 97.7% at trough and 98.7% at peak, very similar to results observed in laMCRC. Additionally, the concentration of PD-1/PD-L1 complexes on cells in the TS and TDLN decreases by more than 99% throughout treatment, as nearly all PD-1 receptors become bound by pembrolizumab, forming PD-1/pembrolizumab complexes. Consequently, there is enhanced lysis of cancer cells by effector CD8+ T cells and activated NK cells, reduced inhibition of pro-inflammatory T cell proliferation and activation, and a decrease in the number of activated and effector Tregs. Thus, we see that the extent of PD-1 receptor engagement and reduction in PD-1/PD-L1 complex concentration are critical factors in influencing treatment efficacy and success.

It is also beneficial for us to compare and analyse the time traces of TVR, efficacy, efficiency, and toxicity of Treatments 1–6 as shown in Figure 8. As expected, the TVRs and efficacies of Treatments 1 and 2 are similar to those of Treatments 3–4 throughout the treatment period, with efficacy being a monotonically increasing function of time. We note that the TVRs and efficacies of Treatments 5 and 6 are slightly lower than those of the other treatments. However, it is important to note that PD-1 receptor engagement by pembrolizumab saturates at relatively low doses, with the KEYNOTE-001 study demonstrating that 2 mg/kg of pembrolizumab is sufficient to saturate unbound PD-1 receptors and achieve maximal anti-tumour activity [105]. Therefore, despite the lower dosages and extended dosing intervals of the optimal regimens, the optimal treatments are significantly more efficient than Treatments 1 and 2, while maintaining comparable efficacy. As a result, the optimal dosing regimens are more efficient and exhibit lower overall dosing than the FDA-approved regimens while still achieving comparable efficacy and TVR. Finally, as expected, the toxicity of all treatments is generally a non-increasing function of time; however, if pembrolizumab concentrations are sufficiently high, small spikes in toxicity may occur following dose administration.

We now shift our focus to Figure 7. We see that TVR increases as the dosing increases and spacing decreases, though with diminishing returns at higher doses or shorter intervals. In particular, the TVRs and efficacies of all optimal regimens are high, with minimal deviations near these regions.

In the spirit of completeness, we verify that Treatments 1 and 2, the FDA-approved regimens, are non-toxic and compare their toxicity to that of the optimal regimens found. As expected, Treatments 1 and 2 are non-toxic, with toxicities of 1.83 × 10^−1^ and 2.37 × 10^−1^, respectively, whilst the optimal regimens have lower or comparable toxicity. Treatments 5 and 6, consisting of 136 mg and 96 mg of pembrolizumab administered every four weeks, are of particular interest, as they achieve comparable TVR and efficacy to Treatments 1 and 2 while being significantly less toxic, making them potentially better options for vulnerable patient populations.

Unsurprisingly, the FDA-approved treatments are very efficient, with the efficiency of Treatment 1 and 2 being approximately 9.71 × 10^−3^%/mg by 96 weeks. However, these pale in comparison to the other optimal regimens, particularly Treatment 6, which has an efficiency of 2.61 × 10^−2^%/mg — more than twice that of Treatments 1 and 2. There is also a clear transition between efficient and inefficient treatments, marked by the rapid shift in efficiency as one deviates from local optima. A treatment is inefficient if its TVR is low, regardless of the dosing and spacing (corresponding to the top left inefficient region in Figure 7), or if an excessive amount of pembrolizumab is administered, regardless of the TVR (corresponding to the bottom right inefficient region in Figure 7).

Striking a balance between TVR, efficiency, and toxicity is challenging, and the current FDA-approved regimens for mMCRC achieve this balance reasonably well. Nonetheless, the optimal regimens defined by Treatments 3–6 in Table 11 are more efficient, lead to comparable TVR, and are more cost-effective and convenient than current regimens, all while maintaining practicality and safety.

It should be noted that the model has several limitations, many of which exist for simplicity, but addressing these issues offers exciting avenues for future research.

- We ignored spatial effects in the model, however, their resolution can provide information about the distribution and clustering of different immune cell types in the TME and their clinical implications [106, 107].
- We assumed that the death rates were constant throughout the T cell proliferation program; however, linear death rates were shown to markedly improve the quality of fit of Deenick et al.’s model [108] to experimental data [109].
- We considered only the M1/M2 macrophage dichotomy, however, their plasticity motivates the description of their phenotypes as a continuum, giving them the ability to adapt their functions to achieve mixtures of M1/M2 responses and functions [49].
- In the optimisation of pembrolizumab therapy, we restricted ourselves to treatments with constant dosing and spacing as is common in the literature; however, varying dosages and dosing frequencies may result in improved regimens.
- We did not consider T cell avidity, the overall strength of a TCR-pMHC interaction, which governs whether a cancer cell will be successfully killed [65]. In particular, high-avidity T cells are necessary for lysing cancer cells and durable tumour eradication, while low-avidity T cells are ineffective and may inhibit high-avidity T cells [110, 111].
- We also did not consider the influence of cytokines in the TDLN for T cell activation and proliferation, which are important in influencing effector T cell differentiation [112, 113].
- The definition of toxicity does not account for its potential origins in autoimmunity, which is a crucial component of certain adverse effects [93].
- The model does not explicitly account for additional anatomical compartments such as the spleen, nor does it directly model metastasis, which may impact the accuracy of systemic immune dynamics and tumour-specific responses.
- The dnmMCRC model does not apply in many cases of recurrent mMCRC, as these cases often lack a primary tumour.

In this work, we have extended our model of laMCRC to mathematically model many immune cell types in the TME of dnmMCRC, using experimental data to govern parameter estimation, and finally analysing and optimising pembrolizumab therapy for TVR, efficiency, and toxicity. We conclude that, although pembrolizumab monotherapy leads to partial clinical response, it is insufficient for complete tumour eradication in dnmMCRC, highlighting the need for additional therapeutic agents such as anti-CTLA-4 antibodies for Treg depletion. This is in contrast to laMCRC, where pembrolizumab monotherapy is sufficient. Furthermore, we demonstrate that administering low to medium doses of pembrolizumab every four weeks achieves comparable efficacy to FDA-approved regimens while offering reduced toxicity and improved treatment efficiency. Overall, this work lays the foundation for deeper insights into tumour-immune dynamics and offers a flexible framework that can be expanded to advance cancer research and improve therapeutic outcomes.

## 6 CRediT authorship contribution statement

**Georgio Hawi:** conceptualisation, data curation, formal analysis, funding acquisition, investigation, methodology, project administration, resources, software, validation, visualisation, writing — original draft, writing — review & editing.

**Peter S. Kim:** conceptualisation, formal analysis, funding acquisition, investigation, methodology, project administration, resources, supervision, validation, visualisation, writing — original draft, writing — review & editing.

**Peter P. Lee:** conceptualisation, formal analysis, investigation, methodology, project administration, resources, supervision, validation, visualisation, writing — original draft, writing — review & editing.

## 8 Data availability

All data and procedures are available within the manuscript and its Supporting Information file.

## 9 Acknowledgements

This work was supported by an Australian Government Research Training Program Scholarship. PSK gratefully acknowledges support from the Australian Research Council Discovery Project (DP230100485).

