## Appendix A for "Optimisation of pembrolizumab therapy for de novo metastatic MSI-H/dMMR colorectal cancer using data-driven delay integro-differential equations"

### A Mathematical Model Derivation

The derivation of the equations for the full model follows an almost identical approach to [1], but is repeated here for the sake of completeness.

#### A.1 Equations for Cancer Cells, DAMPs, and DCs

##### A.1.1 Equations for Cancer Cells ($C$ and $N_c$)

Viable cancer cells are killed by effector CD8+ T cells [2] and activated NK cells [3] through direct contact, whilst TNF and IFN-$\gamma$ indirectly eliminate cancer cells via activating cell death pathways [4–6]. In particular, TNF and IFN-$\gamma$ induce the necroptosis, programmed necrotic cell death, of cancer cells [5, 7]. We note that TGF-$\beta$ and the PD-1/PD-L1 complex inhibit cancer cell lysis by CD8+ T cells [8–10], and that TGF-$\beta$ and PD-1/PD-L1 have been shown to inhibit NK cell cytotoxicity [11–16]. We assume that viable cancer cells grow logistically, as is done in many CRC models [17–19], due to space and resource competition in the TME. Combining these, we have

$$\frac{dC}{dt} = \underbrace{\lambda_C C\left(1-\frac{C}{C_0}\right)}_{\text{growth}} - \underbrace{\lambda_{CT_8}T_8\frac{1}{1+I_\beta/K_{CI_\beta}}\frac{1}{1+Q^{T_8}/K_{CQ^{T_8}}}C}_{\substack{\text{elimination by } T_8\\ \text{inhibited by } I_\beta \text{ and } Q^{T_8}}} - \underbrace{\lambda_{CK}K\frac{1}{1+I_\beta/K_{CI_\beta}}\frac{1}{1+Q^{K}/K_{CQ^{K}}}C}_{\substack{\text{elimination by } K\\ \text{inhibited by } I_\beta \text{ and } Q^{K}}}$$
$$- \underbrace{\lambda_{CI_\alpha}\frac{I_\alpha}{K_{CI_\alpha}+I_\alpha}C}_{\text{elimination by } I_\alpha} - \underbrace{\lambda_{CI_\gamma}\frac{I_\gamma}{K_{CI_\gamma}+I_\gamma}C}_{\text{elimination by } I_\gamma}, \tag{A.1}$$

$$\frac{dN_c}{dt} = \underbrace{\lambda_{CI_\alpha}\frac{I_\alpha}{K_{CI_\alpha}+I_\alpha}C}_{\text{Elimination by } I_\alpha} + \underbrace{\lambda_{CI_\gamma}\frac{I_\gamma}{K_{CI_\gamma}+I_\gamma}C}_{\text{Elimination by } I_\gamma} - \underbrace{d_{N_c}N_c}_{\text{Removal}}. \tag{A.2}$$

##### A.1.2 Equation for Primary Tumour Volume ($V_{\text{TS}}$)

However, it is natural to consider the primary tumour volume, $V_{\text{TS}}$, as it is experimentally easier to measure directly. To convert between $C$, $N_c$, and $V_{\text{TS}}$, we introduce scaling factors $f_C$ and $f_{N_c}$, so that $C(t) = f_C V_{\text{TS}}(t)$ and $N_c(t) = f_{N_c} V_{\text{TS}}(t)$. Hence $C(t) + N_c(t) = (f_C + f_{N_c})\, V_{\text{TS}}(t) \implies V_{\text{TS}}(t) = \frac{C(t)+N_c(t)}{f_C+f_{N_c}}$. Thus, substituting these into the sum of (A.1) and (A.2) leads to

$$\begin{aligned}\frac{dV_{\text{TS}}}{dt} = \frac{1}{f_C+f_{N_c}}\Bigg[&\lambda_C f_C V_{\text{TS}}\left(1-\frac{f_C V_{\text{TS}}}{C_0}\right) - \lambda_{CT_8}T_8\frac{1}{1+I_\beta/K_{CI_\beta}}\frac{1}{1+Q^{T_8}/K_{CQ^{T_8}}}f_C V_{\text{TS}} \\ &-\lambda_{CK}K\frac{1}{1+I_\beta/K_{CI_\beta}}\frac{1}{1+Q^{K}/K_{CQ^{K}}}f_C V_{\text{TS}} - d_{N_c}f_{N_c}V_{\text{TS}}\Bigg].\end{aligned} \tag{A.3}$$

##### A.1.3 Equation for HMGB1 ($H$)

The molecule HMGB1 is released by necrotic cancer cells [20] so that

$$\frac{dH}{dt} = \underbrace{\lambda_{HN_c}N_c}_{\text{production by } N_c} - \underbrace{d_H H}_{\text{degradation}}. \tag{A.4}$$

##### A.1.4 Equation for Calreticulin ($S$)

Necrotic cancer cells release calreticulin [21] so that

$$\frac{dS}{dt} = \underbrace{\lambda_{SN_c}N_c}_{\text{production by } N_c} - \underbrace{d_S S}_{\text{degradation}}. \tag{A.5}$$

##### A.1.5 Equations for Immature and Mature DCs in the TS ($D_0$ and $D$)

Immature DCs are stimulated to mature via DAMPs such as HMGB1 and calreticulin [22]; however, we employ Michaelis-Menten kinetics to account for the limited rate of receptor recycling time [23]. In addition, activated NK cells have been shown to efficiently kill immature DCs but not mature DCs; however, this is inhibited by TGF-β [24–26]. We also need to consider that some mature DCs migrate into the T cell zone of the TDLN and stimulate naive T cells, causing them to be activated [27, 28]. Assuming that immature DCs are supplied at a rate $\mathcal{A}_{D_0}$, we have that

$$\frac{dD_0}{dt} = \underbrace{\mathcal{A}_{D_0}}_{\text{source}} - \underbrace{\lambda_{DH}D_0\frac{H}{K_{DH}+H}}_{D_0\to D \text{ by } H} - \underbrace{\lambda_{DS}D_0\frac{S}{K_{DS}+S}}_{D_0\to D \text{ by } S} - \underbrace{\lambda_{D_0K}D_0K\frac{1}{1+I_\beta/K_{D_0I_\beta}}}_{\substack{\text{elimination by } K\\ \text{inhibited by } I_\beta}} - \underbrace{d_{D_0}D_0}_{\text{death}}, \tag{A.6}$$

$$\frac{dD}{dt} = \underbrace{\lambda_{DH}D_0\frac{H}{K_{DH}+H}}_{D_0\to D \text{ by } H} + \underbrace{\lambda_{DS}D_0\frac{S}{K_{DS}+S}}_{D_0\to D \text{ by } S} - \underbrace{\lambda_{DD^{\text{LN}}}D}_{\substack{D \text{ migration}\\ \text{to TDLN}}} - \underbrace{d_D D}_{\text{death}}. \tag{A.7}$$

##### A.1.6 Equation for Mature DCs in the TDLN ($D^{\mathrm{LN}}$)

We assume a fixed DC migration time of $\tau_m$ and also assume that only $\exp(-d_D\tau_m)$ of the mature DCs that leave the TS survive migration. Taking into account the volume change between the TS and the TDLN, we have that

$$\frac{dD^{\mathrm{LN}}}{dt} = \frac{V_{\mathrm{TS}}}{V_{\mathrm{LN}}} \underbrace{\lambda_{DD^{\mathrm{LN}}} \exp(-d_D\tau_m)\, D(t-\tau_m)}_{D \text{ migration to TDLN}} - \underbrace{d_D D^{\mathrm{LN}}}_{\text{death}}. \tag{A.8}$$

#### A.2 Equations for T Cells

##### A.2.1 Equation for Naive CD8+ T Cells in the TDLN ($T_0^8$)

We assume that naive CD8+ T cells come into the TDLN at a constant rate and that they have not undergone cell division, nor will they until their activation. For simplicity, we do not consider cytokines in the TDLN, absorbing their influence into $\lambda_{T_0^8 T_A^8}$. We do, however, explicitly take into account the influence of effector Tregs and the PD-1/PD-L1 complex in the TDLN, which have been shown to inhibit T cell activation via mechanisms including limiting naive T cells from binding to mature DCs [29–37]. Recalling that T cells that have become activated by mature DCs are no longer naive, and taking this all into account, leads to

$$\frac{dT_0^8}{dt} = \underbrace{\mathcal{A}_{T_0^8}}_{\text{source}} - \underbrace{R^8(t)}_{\substack{\text{CD8+ T cell}\\ \text{activation}}} - \underbrace{d_{T_0^8} T_0^8}_{\text{death}}, \tag{A.9}$$

where $R^8(t)$ is defined as

$$R^8(t) := \underbrace{\frac{\lambda_{T_0^8 T_A^8} \exp\left(-d_{T_0^8}\tau_8^{\mathrm{act}}\right) D^{\mathrm{LN}}(t-\tau_8^{\mathrm{act}}) T_0^8(t-\tau_8^{\mathrm{act}})}{\left(1+\int_{t-\tau_8^{\mathrm{act}}}^t T_A^r(s)\, ds / K_{T_0^8 T_A^r}\right)\left(1+\int_{t-\tau_8^{\mathrm{act}}}^t Q^{8\mathrm{LN}}(s)\, ds / K_{T_0^8 Q^{8\mathrm{LN}}}\right)}}_{\text{CD8+ T cell activation inhibited by } T_A^r \text{ and } Q^{8\mathrm{LN}}}. \tag{A.10}$$

In particular, since effector Tregs and the PD-1/PD-L1 complex inhibit T cell activation during the whole activation process, it is not sufficient to consider point estimates of effector Treg and PD-1/PD-L1 concentration. Instead, we resort to considering the integrals of the concentrations of the relevant species throughout the entire $\tau_8^{\mathrm{act}}$ time that the CD8+ T cell takes to complete activation. This is because these integrals are proportional (with a proportionality constant of $1/\tau_8^{\mathrm{act}}$) to the average concentration of these species throughout activation, allowing us to properly incorporate their inhibition by effector Tregs and the PD-1/PD-L1 complex.

##### A.2.2 Equation for Effector CD8+ T Cells in the TDLN ($T_A^8$)

It is known that activated CD8+ T cells undergo clonal expansion in the TDLN and differentiate before they stop proliferating and migrate to the TS [38, 39].

We assume that activated CD8+ T cells proliferate up to $n_{\max}^8$ times, upon which they stop dividing. For simplicity, we assume that the death rate of CD8+ T cells that have not completed their division program is equal to $d_{T_0^8}$, the death rate of naive CD8+ T cells, regardless of the number of cell divisions previously undergone. We also assume that only activated CD8+ T cells that have undergone $n^8_{\max}$ divisions become effector CD8+ T cells, which will leave the TDLN and migrate to the TS. Furthermore, we assume a constant cell cycle time of $\Delta_8$, except for the first cell division, which has a cycle time of $\Delta_8^0$. Thus, the duration of the activated CD8+ T cell division program to $n^8_{\max}$ divisions is given by

$$\tau_{T_A^8} := \Delta_8^0 + (n^8_{\max} - 1)\Delta_8. \tag{A.11}$$

In particular, we must take into account that some T cells will die before the division program is complete, so we must introduce a shrinkage factor of $\exp\left(-d_{T_0^8}\tau_{T_A^8}\right)$. Furthermore, we must also take into account that effector Tregs and the PD-1/PD-L1 complex inhibit CD8+ T cell proliferation throughout the program [30–32, 40, 41]. We must also consider that some of these effector CD8+ T cells will migrate to the TS to perform effector functions. We finally assume that the death rate of CD8+ T cells that have completed their division program is equal to the death rate of CD8+ T cells in the TS. Taking this all into account leads to

$$\frac{dT_A^8}{dt} = \underbrace{\frac{2^{n^8_{\max}} \exp\left(-d_{T_0^8}\tau_{T_A^8}\right) R^8(t-\tau_{T_A^8})}{\left(1+\int_{t-\tau_{T_A^8}}^t T_A^r(s)\,ds/K_{T_A^8T_A^r}\right)\left(1+\int_{t-\tau_{T_A^8}}^t Q^{8\mathrm{LN}}(s)\,ds/K_{T_A^8Q^{8\mathrm{LN}}}\right)}}_{\text{CD8+ T cell proliferation inhibited by } T_A^r \text{ and } Q^{8\mathrm{LN}}} - \underbrace{\lambda_{T_A^8T_8}T_A^8}_{\substack{T_A^8 \text{ migration}\\ \text{to the TS}}} - \underbrace{d_{T_8}T_A^8}_{\text{death}}. \tag{A.12}$$

##### A.2.3 Equations for Effector and Exhausted CD8+ T Cells in the TS ($T_8$ and $T_{\mathrm{ex}}$)

We assume that it takes $\tau_a$ amount of time for effector CD8+ T cells in the TDLN to migrate to the TS. We must also account for CTL expansion due to IL-2 [42], noting that this proliferation is inhibited by effector Tregs [30–32]. Furthermore, the death of CD8+ T cells is resisted by IL-10 [43, 44].

However, chronic antigen exposure can cause effector CD8+ T cells to enter a state of exhaustion, where they lose their ability to kill cancer cells, and the rate of cytokine secretion significantly decreases [45–47]. We denote this exhausted CD8+ T cell population as $T_{\mathrm{ex}}(t)$. It has also been shown that pembrolizumab can "reinvigorate" these cells back into the effector state [48, 49]. We model the re-invigoration and exhaustion using Michaelis-Menten terms in $A_1$ and $\int_{t-\tau_l}^t C(s)\,ds$ respectively, where $\tau_l$ is the median time that CD8+ T cells take to become exhausted after entering the TS. In particular, this has been shown to be more appropriate than simple mass-action kinetics as it accounts for extended antigen exposure [50].

As such, remembering to take the volume change between the TDLN and the TS into account, this implies that

$$\begin{aligned}\frac{dT_8}{dt} = \frac{V_{\mathrm{LN}}}{V_{\mathrm{TS}}}\underbrace{\lambda_{T_A^8T_8}\exp\left(-d_{T_8}\tau_a\right)T_A^8(t-\tau_a)}_{T_A^8 \text{ migration to the TS}} + \underbrace{\lambda_{T_8I_2}\frac{T_8I_2}{K_{T_8I_2}+I_2}\frac{1}{1+T_r/K_{T_8T_r}}}_{\text{growth by } I_2 \text{ inhibited by } T_r} \\ - \underbrace{\lambda_{T_8C}\frac{T_8\int_{t-\tau_l}^t C(s)\,ds}{K_{T_8C}+\int_{t-\tau_l}^t C(s)\,ds}}_{T_8\to T_{\mathrm{ex}} \text{ from } C \text{ exposure}} + \underbrace{\lambda_{T_{\mathrm{ex}}A_1}\frac{T_{\mathrm{ex}}A_1}{K_{T_{\mathrm{ex}}A_1}+A_1}}_{T_{\mathrm{ex}}\to T_8 \text{ by } A_1} - \underbrace{\frac{d_{T_8}T_8}{1+I_{10}/K_{T_8I_{10}}}}_{\substack{\text{death}\\ \text{inhibited by } I_{10}}},\end{aligned} \tag{A.13}$$

$$\frac{dT_{\text{ex}}}{dt} = \underbrace{\lambda_{T_8C}\frac{T_8\int_{t-\tau_l}^{t}C(s)\ ds}{K_{T_8C}+\int_{t-\tau_l}^{t}C(s)\ ds}}_{T_8\to T_{\text{ex}}\text{ from } C \text{ exposure}} - \underbrace{\lambda_{T_{\text{ex}}A_1}\frac{T_{\text{ex}}A_1}{K_{T_{\text{ex}}A_1}+A_1}}_{T_{\text{ex}}\to T_8 \text{ by } A_1} - \underbrace{\frac{d_{T_{\text{ex}}}T_{\text{ex}}}{1+I_{10}/K_{T_{\text{ex}}I_{10}}}}_{\substack{\text{death}\\ \text{inhibited by } I_{10}}}. \tag{A.14}$$

##### A.2.4 Equation for Naive CD4+ T Cells in the TDLN ($T_0^4$)

For simplicity, we consider only the Th1 subtype that naive CD4+ T cells differentiate into upon activation, absorbing the influence of cytokines via the kinetic rate constant $\lambda_{T_0^4T_A^1}$. Taking into account that effector Tregs and the PD-1/PD-L1 complex inhibit Th1 cell activation and some mature DCs migrate into the TDLN and activate naive CD4+ T cells, causing them to no longer be naive, and assuming that naive CD4+ T cells come into the TDLN at a rate $\mathcal{A}_{T_0^4}$, we can write a similar equation to (A.9):

$$\frac{dT_0^4}{dt} = \underbrace{\mathcal{A}_{T_0^4}}_{\text{source}} - \underbrace{R^1(t)}_{\text{Th1 cell activation}} - \underbrace{d_{T_0^4}T_0^4}_{\text{death}}, \tag{A.15}$$

where $R^1(t)$ is defined as

$$R^1(t) := \underbrace{\frac{\lambda_{T_0^4T_A^1}\exp\left(-d_{T_0^4}\tau_{\text{act}}^4\right)D^{\text{LN}}(t-\tau_{\text{act}}^4)T_0^4(t-\tau_{\text{act}}^4)}{\left(1+\int_{t-\tau_4^{\text{act}}}^{t}T_r^A(s)\ ds/K_{T_0^4T_A^r}\right)\left(1+\int_{t-\tau_4^{\text{act}}}^{t}Q^{1\text{LN}}(s)\ ds/K_{T_0^4Q^{1\text{LN}}}\right)}}_{\text{Th1 cell activation inhibited by } T_A^r \text{ and } Q^{1\text{LN}}}. \tag{A.16}$$

##### A.2.5 Equation for Effector Th1 Cells in the TDLN ($T_A^1$)

We assume that Th1 cells proliferate up to $n_{\text{max}}^1$ times, upon which they stop dividing and become effector cells. As before, we assume that the death rate of Th1 cells that have not completed their division program is equal to $d_{T_0^4}$, the death rate of naive CD4+ T cells, regardless of the number of cell divisions previously undergone. We assume a constant cell cycle time of $\Delta_1$, except for the first cell division, which has a cycle time of $\Delta_1^0$. Thus, the duration of the Th1 cell division program to $n_{\text{max}}^1$ divisions is given by

$$\tau_{T_A^1} := \Delta_1^0 + (n_{\text{max}}^1 - 1)\Delta_1. \tag{A.17}$$

In particular, we must take into account that some Th1 cells will die before the division program is complete, so we must introduce a shrinkage factor of $\exp\left(-d_{T_0^4}\tau_{T_A^1}\right)$. Furthermore, we must also take into account that effector Tregs and the PD-1/PD-L1 complex inhibit Th1 cell proliferation throughout their program. We also assume that the death rate of Th1 cells that have completed their division program is equal to the corresponding degradation rate in the TS. Taking this all into account, and incorporating effector Th1 cell migration to the TS, leads to

$$\frac{dT_A^1}{dt} = \underbrace{\frac{2^{n_{\text{max}}^1}\exp\left(-d_{T_0^4}\tau_{T_A^1}\right)R^1(t-\tau_{T_A^1})}{\left(1+\int_{t-\tau_{T_A^1}}^{t}Q^{1\text{LN}}(s)\ ds/K_{T_A^1Q^{1\text{LN}}}\right)\left(1+\int_{t-\tau_{T_A^1}}^{t}T_A^r(s)\ ds/K_{T_A^1T_A^r}\right)}}_{\text{Th1 cell proliferation inhibited by } T_A^r \text{ and } Q^{1\text{LN}}} - \underbrace{\lambda_{T_A^1T_1}T_A^1}_{\substack{T_A^1 \text{ migration}\\ \text{to the TS}}} - \underbrace{d_{T_1}T_A^1}_{\text{death}}. \tag{A.18}$$

##### A.2.6 Equation for Effector Th1 Cells in the TS ($T_1$)

We assume that it takes $\tau_a$ amount of time for these cells to migrate to the TS. We take into account the fact that IL-2 induces the growth of effector Th1 cells [51], noting that this proliferation is inhibited by effector Tregs [30–32]. Furthermore, the PD-1/PD-L1 axis converts Th1 cells to Tregs [52, 53], a process we consider to be mediated by the PD-1/PD-L1 complex on Th1 cells. Thus, we have that

$$\frac{dT_1}{dt} = \frac{V_{\text{LN}}}{V_{\text{TS}}} \underbrace{\lambda_{T_A^1 T_1} \exp\left(-d_{T_1}\tau_a\right) T_A^1(t-\tau_a)}_{T_A^1 \text{ migration to the TS}} + \underbrace{\lambda_{T_1 I_2} \frac{T_1 I_2}{K_{T_1 I_2} + I_2} \frac{1}{1 + T_r/K_{T_1 T_r}}}_{\text{growth by } I_2 \text{ inhibited by } T_r} - \underbrace{\lambda_{T_1 T_r} T_1 \frac{Q^{T_1}}{K_{T_1 Q^{T_1}} + Q^{T_1}}}_{T_1 \to T_r \text{ by } Q^{T_1}} - \underbrace{d_{T_1} T_1}_{\text{death}}. \tag{A.19}$$

##### A.2.7 Equation for Naive Tregs in the TDLN ($T_0^r$)

Finally, we consider the concentration of naive Tregs in the TDLN, following the same procedure as for CD8+ T cells and Th1 cells. We absorb the influence of cytokines on Treg activation via the kinetic rate constant $\lambda_{T_0^r T_A^r}$. We also take into account that some mature DCs migrate into the TDLN and activate naive Tregs, causing them to no longer be naive. Assuming that naive Tregs come into the TDLN at a rate $\mathcal{A}_{T_0^r}$, we can write a similar equation to (A.9) and (A.15):

$$\frac{dT_0^r}{dt} = \underbrace{\mathcal{A}_{T_0^r}}_{\text{source}} - \underbrace{R^r(t)}_{\text{Treg activation}} - \underbrace{d_{T_0^r} T_0^r}_{\text{death}}, \tag{A.20}$$

where $R^r(t)$ is defined as

$$R^r(t) := \underbrace{\lambda_{T_0^r T_A^r} \exp\left(-d_{T_0^r}\tau_{\text{act}}^r\right) D^{\text{LN}}(t-\tau_{\text{act}}^r) T_0^r(t-\tau_{\text{act}}^r)}_{\text{Treg activation}}. \tag{A.21}$$

##### A.2.8 Equation for Effector Tregs in the TDLN ($T_A^r$)

We assume that activated Tregs proliferate up to $n_{\text{max}}^r$ times, upon which they stop dividing and become effector Tregs. As before, we assume that the death rate of Tregs that have not completed their division program is equal to $d_{T_0^r}$, the death rate of naive Tregs. We assume a constant cell cycle time of $\Delta_r$, except for the first cell division, which has a cycle time of $\Delta_r^0$. Thus, the duration of the activated Treg division program to $n_{\text{max}}^r$ divisions is given by

$$\tau_{T_A^r} := \Delta_r^0 + (n_{\text{max}}^r - 1)\Delta_r. \tag{A.22}$$

In particular, we must take into account that some T cells will die before the division program is complete, so we must introduce a shrinkage factor of $\exp\left(-d_{T_0^r}\tau_{T_A^r}\right)$. We also assume that the death rate of effector Tregs in the TDLN is equal to the corresponding degradation rate in the TS. Taking this all into account, and incorporating effector Treg migration to the TS, leads to

$$\frac{dT_A^r}{dt} = \underbrace{2^{n_{\text{max}}^r} \exp\left(-d_{T_0^r}\tau_{T_A^r}\right) R^r(t-\tau_{T_A^r})}_{\text{Treg proliferation}} - \underbrace{\lambda_{T_A^r T_r} T_A^r}_{T_A^r \text{ migration to the TS}} - \underbrace{d_{T_r} T_A^r}_{\text{death}}. \tag{A.23}$$

##### A.2.9 Equation for Effector Tregs in the TS ($T_r$)

Assuming that it also takes $\tau_a$ amount of time for Tregs to migrate to the TS, we have that

$$\frac{dT_r}{dt} = \frac{V_{\mathrm{LN}}}{V_{\mathrm{TS}}} \underbrace{\lambda_{T_A^r T_r} \exp\left(-d_{T_r}\tau_a\right) T_A^r(t-\tau_a)}_{T_A^r \text{ migration to the TS}} + \underbrace{\lambda_{T_1 T_r} T_1 \frac{Q^{T_1}}{K_{T_1 Q^{T_1}} + Q^{T_1}}}_{T_1 \to T_r \text{ by } Q^{T_1}} - \underbrace{d_{T_r} T_r}_{\text{death}}. \tag{A.24}$$

#### A.3 Equations for Other Immune Cells in the TS

##### A.3.1 Equations for Naive, M1, and M2 Macrophages ($M_0$, $M_1$, and $M_2$)

TNF and IFN-γ polarise naive macrophages into M1 macrophages [54–57], whilst IL-10 and TGF-β polarise naive macrophages into the M2 phenotype [58–60]. In addition, TGF-β induces M1 macrophages to convert into M2 macrophages [60]. Furthermore, M2 macrophages change phenotype to M1 under the influence of TNF [54] and IFN-γ [61]. Assuming a production rate $\mathcal{A}_{M_0}$ of naive macrophages, we thus have that

$$\begin{aligned}\frac{dM_0}{dt} = {} & \underbrace{\mathcal{A}_{M_0}}_{\text{source}} - \underbrace{\lambda_{M_1 I_\alpha} M_0 \frac{I_\alpha}{K_{M_1 I_\alpha} + I_\alpha}}_{M_0 \to M_1 \text{ by } I_\alpha} - \underbrace{\lambda_{M_1 I_\gamma} M_0 \frac{I_\gamma}{K_{M_1 I_\gamma} + I_\gamma}}_{M_0 \to M_1 \text{ by } I_\gamma} - \underbrace{\lambda_{M_2 I_{10}} M_0 \frac{I_{10}}{K_{M_2 I_{10}} + I_{10}}}_{M_0 \to M_2 \text{ by } I_{10}} \\ & - \underbrace{\lambda_{M_2 I_\beta} M_0 \frac{I_\beta}{K_{M_2 I_\beta} + I_\beta}}_{M_0 \to M_2 \text{ by } I_\beta} - \underbrace{d_{M_0} M_0}_{\text{degradation}},\end{aligned} \tag{A.25}$$

$$\begin{aligned}\frac{dM_1}{dt} = {} & \underbrace{\lambda_{M_1 I_\alpha} M_0 \frac{I_\alpha}{K_{M_1 I_\alpha} + I_\alpha}}_{M_0 \to M_1 \text{ by } I_\alpha} + \underbrace{\lambda_{M_1 I_\gamma} M_0 \frac{I_\gamma}{K_{M_1 I_\gamma} + I_\gamma}}_{M_0 \to M_1 \text{ by } I_\gamma} + \underbrace{\lambda_{M I_\gamma} M_2 \frac{I_\gamma}{K_{M I_\gamma} + I_\gamma}}_{M_2 \to M_1 \text{ by } I_\gamma} + \underbrace{\lambda_{M I_\alpha} M_2 \frac{I_\alpha}{K_{M I_\alpha} + I_\alpha}}_{M_2 \to M_1 \text{ by } I_\alpha} \\ & - \underbrace{\lambda_{M I_\beta} M_1 \frac{I_\beta}{K_{M I_\beta} + I_\beta}}_{M_1 \to M_2 \text{ by } I_\beta} - \underbrace{d_{M_1} M_1}_{\text{degradation}},\end{aligned} \tag{A.26}$$

$$\begin{aligned}\frac{dM_2}{dt} = {} & \underbrace{\lambda_{M_2 I_{10}} M_0 \frac{I_{10}}{K_{M_2 I_{10}} + I_{10}}}_{M_0 \to M_2 \text{ by } I_{10}} + \underbrace{\lambda_{M_2 I_\beta} M_0 \frac{I_\beta}{K_{M_2 I_\beta} + I_\beta}}_{M_0 \to M_2 \text{ by } I_\beta} - \underbrace{\lambda_{M I_\gamma} M_2 \frac{I_\gamma}{K_{M I_\gamma} + I_\gamma}}_{M_2 \to M_1 \text{ by } I_\gamma} - \underbrace{\lambda_{M I_\alpha} M_2 \frac{I_\alpha}{K_{M I_\alpha} + I_\alpha}}_{M_2 \to M_1 \text{ by } I_\alpha} \\ & + \underbrace{\lambda_{M I_\beta} M_1 \frac{I_\beta}{K_{M I_\beta} + I_\beta}}_{M_1 \to M_2 \text{ by } I_\beta} - \underbrace{d_{M_2} M_2}_{\text{degradation}}.\end{aligned} \tag{A.27}$$

##### A.3.2 Equations for Resting and Activated NK Cells ($K_0$ and $K$)

Resting NK cells are activated by IL-2 [62, 63] and immature and mature DCs [64]. However, NK cell activation is inhibited by TGF-β [65]. Thus, assuming a supply rate $\mathcal{A}_{K_0}$ of resting NK cells, we have that

$$\frac{dK_0}{dt} = \underbrace{\mathcal{A}_{K_0}}_{\text{source}} - \left( \underbrace{\lambda_{KI_2} K_0 \frac{I_2}{K_{KI_2} + I_2}}_{K_0 \to K \text{ by } I_2} + \underbrace{\lambda_{KD_0} K_0 \frac{D_0}{K_{KD_0} + D_0}}_{K_0 \to K \text{ by } D_0} + \underbrace{\lambda_{KD} K_0 \frac{D}{K_{KD} + D}}_{K_0 \to K \text{ by } D} \right) \underbrace{\frac{1}{1 + I_\beta / K_{KI_\beta}}}_{\substack{\text{activation} \\ \text{inhibited by } I_\beta}} - \underbrace{d_{K_0} K_0}_{\text{degradation}} , \tag{A.28}$$

$$\frac{dK}{dt} = \left( \underbrace{\lambda_{KI_2} K_0 \frac{I_2}{K_{KI_2} + I_2}}_{K_0 \to K \text{ by } I_2} + \underbrace{\lambda_{KD_0} K_0 \frac{D_0}{K_{KD_0} + D_0}}_{K_0 \to K \text{ by } D_0} + \underbrace{\lambda_{KD} K_0 \frac{D}{K_{KD} + D}}_{K_0 \to K \text{ by } D} \right) \underbrace{\frac{1}{1 + I_\beta / K_{KI_\beta}}}_{\substack{\text{activation} \\ \text{inhibited by } I_\beta}} - \underbrace{d_K K}_{\text{degradation}} . \tag{A.29}$$

#### A.4 Equations for Cytokines

##### A.4.1 Equation for IL-2 ($I_2$)

IL-2 is produced by effector CD8+ T cells [66, 67] and Th1 cells [68], so that

$$\frac{dI_2}{dt} = \underbrace{\lambda_{I_2 T_8} T_8}_{\text{production by } T_8} + \underbrace{\lambda_{I_2 T_1} T_1}_{\text{production by } T_1} - \underbrace{d_{I_2} I_2}_{\text{degradation}} . \tag{A.30}$$

##### A.4.2 Equation for IFN-γ ($I_\gamma$)

IFN-γ is produced by effector CD8+ T cells [69] and Th1 cells [7, 70], with both expressions being inhibited by Tregs [71]. Furthermore, activated NK cells also produce IFN-γ [72]. Thus,

$$\frac{dI_\gamma}{dt} = \left( \underbrace{\lambda_{I_\gamma T_8} T_8}_{\text{production by } T_8} + \underbrace{\lambda_{I_\gamma T_1} T_1}_{\text{production by } T_1} \right) \underbrace{\frac{1}{1 + T_r / K_{I_\gamma T_r}}}_{\text{inhibition by } T_r} + \underbrace{\lambda_{I_\gamma K} K}_{\text{production by } K} - \underbrace{d_{I_\gamma} I_\gamma}_{\text{degradation}} . \tag{A.31}$$

##### A.4.3 Equation for TNF ($I_\alpha$)

TNF is produced by effector CD8+ T cells [73, 74] and Th1 cells [75, 76], M1 macrophages [77], and activated NK cells [78, 79]. Hence,

$$\frac{dI_\alpha}{dt} = \underbrace{\lambda_{I_\alpha T_8} T_8}_{\text{production by } T_8} + \underbrace{\lambda_{I_\alpha T_1} T_1}_{\text{production by } T_1} + \underbrace{\lambda_{I_\alpha M_1} M_1}_{\text{production by } M_1} + \underbrace{\lambda_{I_\alpha K} K}_{\text{production by } K} - \underbrace{d_{I_\alpha} I_\alpha}_{\text{degradation}} . \tag{A.32}$$

##### A.4.4 Equation for TGF-β ($I_\beta$)

TGF-β is produced by viable cancer cells [80], effector Tregs [81] and M2 macrophages [59, 82]. Thus,

$$\frac{dI_\beta}{dt} = \underbrace{\lambda_{I_\beta C} C}_{\text{production by } C} + \underbrace{\lambda_{I_\beta T_r} T_r}_{\text{production by } T_r} + \underbrace{\lambda_{I_\beta M_2} M_2}_{\text{production by } M_2} - \underbrace{d_{I_\beta} I_\beta}_{\text{degradation}} . \tag{A.33}$$

##### A.4.5 Equation for IL-10 ($I_{10}$)

IL-10 is produced by viable cancer cells [83, 84] and M2 macrophages [85, 86]. Additionally, effector Tregs secrete IL-10 [87] with IL-2 enhancing this production [88, 89]. Hence,

$$\frac{dI_{10}}{dt} = \underbrace{\lambda_{I_{10}C} C}_{\text{production by } C} + \underbrace{\lambda_{I_{10}M_2} M_2}_{\text{production by } M_2} + \underbrace{\lambda_{I_{10}T_r} T_r \left(1 + \lambda_{I_{10}I_2} \frac{I_2}{K_{I_{10}I_2} + I_2}\right)}_{\text{production by } T_r \text{ enhanced by } I_2} - \underbrace{d_{I_{10}} I_{10}}_{\text{degradation}} . \tag{A.34}$$

#### A.5 Equations for Immune Checkpoint-Associated Components in the TS

##### A.5.1 Equations for Unbound PD-1 receptors on Cells in the TS ($P_D^{T_8}$, $P_D^{T_1}$, $P_D^K$)

It is known that PD-1 is expressed on the surface of effector CD8+ T cells [90–92], effector Th1 cells [93] and activated NK cells [13, 16, 94]. We assume that the rate of PD-1 synthesis is proportional to the concentration of the cell expressing it. However, unbound PD-1 receptors on these PD-1-expressing cells can bind to either pembrolizumab or PD-L1, forming the PD-1/pembrolizumab and PD-1/PD-L1 complexes, respectively, resulting in the depletion of unbound PD-1 molecules [95, 96]. For simplicity, we assume that the formation and dissociation rates of the PD-1/PD-L1 and PD-1/pembrolizumab complexes are invariant of the type of cell expressing PD-1. Considering unbound PD-1 receptors on effector CD8+ T cells in the TS at first, and taking into account the degradation of PD-1 receptors, this motivates the equation for $P_D^{T_8}$ to be

$$\frac{dP_D^{T_8}}{dt} = \underbrace{\lambda_{P_D^{T_8}} T_8}_{\text{synthesis}} + \underbrace{\lambda_{Q_A} Q_A^{T_8}}_{\text{dissociation of } Q_A^{T_8}} + \underbrace{\lambda_Q Q^{T_8}}_{\text{dissociation of } Q^{T_8}} - \underbrace{\lambda_{P_D A_1} P_D^{T_8} A_1}_{\text{binding to } A_1} - \underbrace{\lambda_{P_D P_L} P_D^{T_8} P_L}_{\text{binding to } P_L} - \underbrace{d_{P_D} P_D^{T_8}}_{\text{degradation}} . \tag{A.35}$$

Similarly, we have that

$$\frac{dP_D^{T_1}}{dt} = \underbrace{\lambda_{P_D^{T_1}} T_1}_{\text{synthesis}} + \underbrace{\lambda_{Q_A} Q_A^{T_1}}_{\text{dissociation of } Q_A^{T_1}} + \underbrace{\lambda_Q Q^{T_1}}_{\text{dissociation of } Q^{T_1}} - \underbrace{\lambda_{P_D A_1} P_D^{T_1} A_1}_{\text{binding to } A_1} - \underbrace{\lambda_{P_D P_L} P_D^{T_1} P_L}_{\text{binding to } P_L} - \underbrace{d_{P_D} P_D^{T_1}}_{\text{degradation}} , \tag{A.36}$$

$$\frac{dP_D^{K}}{dt} = \underbrace{\lambda_{P_D^{K}} K}_{\text{synthesis}} + \underbrace{\lambda_{Q_A} Q_A^{K}}_{\text{dissociation of } Q_A^{K}} + \underbrace{\lambda_Q Q^{K}}_{\text{dissociation of } Q^{K}} - \underbrace{\lambda_{P_D A_1} P_D^{K} A_1}_{\text{binding to } A_1} - \underbrace{\lambda_{P_D P_L} P_D^{K} P_L}_{\text{binding to } P_L} - \underbrace{d_{P_D} P_D^{K}}_{\text{degradation}} . \tag{A.37}$$

##### A.5.2 Equations for the PD-1/pembrolizumab Complex on Cells in the TS ($Q_A^{T_8}$, $Q_A^{T_1}$, $Q_A^K$)

Pembrolizumab binds to unbound PD-1 on the surfaces of PD-1-expressing cells in a 1:1 ratio [97], forming the PD-1/pembrolizumab complex in a reversible chemical process [98, 99]. We must also account for loss due to the endocytosis and internalisation of the PD-1/pembrolizumab complex from the surface of cells [100, 101]. We assume that the rates of PD-1/pembrolizumab complex internalisation and dissociation are invariant of the type of cell expressing PD-1, so that

$$\frac{dQ_A^{T_8}}{dt} = \underbrace{\lambda_{P_D A_1} P_D^{T_8} A_1}_{\text{formation of } Q_A^{T_8}} - \underbrace{\lambda_{Q_A} Q_A^{T_8}}_{\text{dissociation of } Q_A^{T_8}} - \underbrace{d_{Q_A} Q_A^{T_8}}_{\text{internalisation}}, \tag{A.38}$$

$$\frac{dQ_A^{T_1}}{dt} = \underbrace{\lambda_{P_D A_1} P_D^{T_1} A_1}_{\text{formation of } Q_A^{T_1}} - \underbrace{\lambda_{Q_A} Q_A^{T_1}}_{\text{dissociation of } Q_A^{T_1}} - \underbrace{d_{Q_A} Q_A^{T_1}}_{\text{internalisation}}, \tag{A.39}$$

$$\frac{dQ_A^{K}}{dt} = \underbrace{\lambda_{P_D A_1} P_D^{K} A_1}_{\text{formation of } Q_A^{K}} - \underbrace{\lambda_{Q_A} Q_A^{K}}_{\text{dissociation of } Q_A^{K}} - \underbrace{d_{Q_A} Q_A^{K}}_{\text{internalisation}}. \tag{A.40}$$

##### A.5.3 Equation for Pembrolizumab in the TS ($A_1$)

We assume that pembrolizumab is administered intravenously at times $t_1$, $t_2$, ..., $t_n$ with doses $\xi_1$, $\xi_2$, ..., $\xi_n$ respectively, assuming that the duration of infusion is negligible in comparison to the time period of interest. We also account for pembrolizumab depletion due to binding to unbound PD-1, replenishment due to PD-1/pembrolizumab complex dissociation, and elimination of pembrolizumab. It is important to note that the administered dose is not equal to the corresponding change in concentration in the TS. For simplicity, we assume linear pharmacokinetics so that, for some scaling factor $f_{\text{pembro}}$, we have that

$$\frac{dA_1}{dt} = \underbrace{\sum_{j=1}^{n} \xi_j f_{\text{pembro}} \delta\left(t - t_j\right)}_{\text{infusion}} + \underbrace{\lambda_{Q_A}\left(Q_A^{T_8} + Q_A^{T_1} + Q_A^{K}\right)}_{\text{dissociation of } Q_A^{T_8},\ Q_A^{T_1},\text{ and } Q_A^{K}} - \underbrace{\lambda_{P_D A_1}\left(P_D^{T_8} + P_D^{T_1} + P_D^{K}\right) A_1}_{\text{formation of } Q_A^{T_8},\ Q_A^{T_1},\text{ and } Q_A^{K}} - \underbrace{d_{A_1} A_1}_{\text{elimination}}. \tag{A.41}$$

##### A.5.4 Equation for Unbound PD-L1 in the TS ($P_L$)

We also know that PD-L1 is expressed on the surface of viable cancer cells [102], mature DCs [103], effector CD8+ T cells [104, 105], effector Th1 cells [106], effector Tregs [107], and M2 macrophages [108]. For brevity, we denote $\mathcal{X}$ as the set of PD-L1-expressing cells in the TS, so that $\mathcal{X} := \{C, D, T_8, T_1, T_r, M_2\}$. Furthermore, $\lambda_{P_L X}$ denotes the synthesis rate of unbound PD-L1 on the surface of $X \in \mathcal{X}$. We must take into account the synthesis of PD-L1, its depletion due to binding to unbound PD-1, replenishment due to PD-1/PD-L1 complex dissociation, and the degradation of PD-L1. Hence,

$$\frac{dP_L}{dt} = \underbrace{\sum_{X \in \mathcal{X}} \lambda_{P_L X} X}_{\text{synthesis}} + \underbrace{\lambda_Q\left(Q^{T_8} + Q^{T_1} + Q^{K}\right)}_{\text{dissociation of } Q^{T_8},\ Q^{T_1} \text{ and } Q^{K}} - \underbrace{\lambda_{P_D P_L}\left(P_D^{T_8} + P_D^{T_1} + P_D^{K}\right) P_L}_{\text{formation of } Q^{T_8},\ Q^{T_1} \text{ and } Q^{K}} - \underbrace{d_{P_L} P_L}_{\text{degradation}}. \tag{A.42}$$

##### A.5.5 Equations for the PD-1/PD-L1 Complex in the TS ($Q^{T_8}$, $Q^{T_1}$, and $Q^K$)

PD-L1 binds to unbound PD-1 receptors on the surfaces of PD-1-expressing cells in a 1:1 ratio [109], forming the PD-1/PD-L1 complex in a reversible chemical process. Considering $Q^{T_8}$ as an example, we can express its formation and dissociation via the reaction $P_D^{T_8} + P_L \underset{\lambda_Q}{\overset{\lambda_{P_D P_L}}{\rightleftharpoons}} Q^{T_8}$. We assume that the degradation is negligible relative to the dissociation, so that

$$\frac{dQ^{T_8}}{dt} = \underbrace{\lambda_{P_D P_L} P_D^{T_8} P_L}_{\text{formation}} - \underbrace{\lambda_Q Q^{T_8}}_{\text{dissociation}}. \tag{A.43}$$

However, the dissociation rate constant of the PD-1/PD-L1 complex is 1.44 $s^{-1}$, corresponding to a mean lifetime of less than 1 second [109]. As such, we employ a quasi-steady-state approximation (QSSA) for $Q^{T_8}$, so that $\frac{dQ^{T_8}}{dt} = 0$, so that

$$Q^{T_8} = \frac{\lambda_{P_D P_L}}{\lambda_Q} P_D^{T_8} P_L. \tag{A.44}$$

Similarly,

$$Q^{T_1} = \frac{\lambda_{P_D P_L}}{\lambda_Q} P_D^{T_1} P_L, \tag{A.45}$$

$$Q^{K} = \frac{\lambda_{P_D P_L}}{\lambda_Q} P_D^{K} P_L. \tag{A.46}$$

Furthermore, we can simplify (A.35) – (A.37) and (A.42) by substituting in (A.44) – (A.46) so that

$$\frac{dP_D^{T_8}}{dt} = \underbrace{\lambda_{P_D^{T_8}} T_8}_{\text{synthesis}} + \underbrace{\lambda_{Q_A} Q_A^{T_8}}_{\substack{\text{dissociation}\\ \text{of } Q_A^{T_8}}} - \underbrace{\lambda_{P_D A_1} P_D^{T_8} A_1}_{\text{binding to } A_1} - \underbrace{d_{P_D} P_D^{T_8}}_{\text{degradation}}, \tag{A.47}$$

$$\frac{dP_D^{T_1}}{dt} = \underbrace{\lambda_{P_D^{T_1}} T_1}_{\text{synthesis}} + \underbrace{\lambda_{Q_A} Q_A^{T_1}}_{\substack{\text{dissociation}\\ \text{of } Q_A^{T_1}}} - \underbrace{\lambda_{P_D A_1} P_D^{T_1} A_1}_{\text{binding to } A_1} - \underbrace{d_{P_D} P_D^{T_1}}_{\text{degradation}}, \tag{A.48}$$

$$\frac{dP_D^{K}}{dt} = \underbrace{\lambda_{P_D^{K}} K}_{\text{synthesis}} + \underbrace{\lambda_{Q_A} Q_A^{K}}_{\substack{\text{dissociation}\\ \text{of } Q_A^{K}}} - \underbrace{\lambda_{P_D A_1} P_D^{K} A_1}_{\text{binding to } A_1} - \underbrace{d_{P_D} P_D^{K}}_{\text{degradation}}, \tag{A.49}$$

$$\frac{dP_L}{dt} = \underbrace{\sum_{X \in \mathcal{X}} \lambda_{P_L X} X}_{\text{synthesis}} - \underbrace{d_{P_L} P_L}_{\text{degradation}}. \tag{A.50}$$

#### A.6 Equations for Immune Checkpoint-Associated Components in the TDLN

##### A.6.1 Equations for Unbound PD-1 receptors on cells in the TDLN ($P_D^{8\text{LN}}$ and $P_D^{1\text{LN}}$)

The equations for $P_D^{8\text{LN}}$ and $P_D^{1\text{LN}}$ follow identically to that of (A.47) – (A.48). For simplicity, we assume that the formation and dissociation rates of the PD-1/pembrolizumab complex are identical in the TDLN and the TS, so that

$$\frac{dP_D^{8\text{LN}}}{dt} = \underbrace{\lambda_{P_D^{8\text{LN}}} T_A^8}_{\text{synthesis}} + \underbrace{\lambda_{Q_A} Q_A^{8\text{LN}}}_{\substack{\text{dissociation}\\ \text{of } Q_A^{8\text{LN}}}} - \underbrace{\lambda_{P_D A_1} P_D^{8\text{LN}} A_1^{\text{LN}}}_{\text{binding to } A_1^{\text{LN}}} - \underbrace{d_{P_D} P_D^{8\text{LN}}}_{\text{degradation}}, \tag{A.51}$$

$$\frac{dP_D^{1\text{LN}}}{dt} = \underbrace{\lambda_{P_D^{1\text{LN}}} T_A^1}_{\text{synthesis}} + \underbrace{\lambda_{Q_A} Q_A^{1\text{LN}}}_{\substack{\text{dissociation}\\ \text{of } Q_A^{1\text{LN}}}} - \underbrace{\lambda_{P_D A_1} P_D^{1\text{LN}} A_1^{\text{LN}}}_{\text{binding to } A_1^{\text{LN}}} - \underbrace{d_{P_D} P_D^{1\text{LN}}}_{\text{degradation}}. \tag{A.52}$$

##### A.6.2 Equations for the PD-1/pembrolizumab Complex on Cells in the TDLN ($Q_A^{8\text{LN}}$ and $Q_A^{1\text{LN}}$)

The equations for $Q_A^{8\text{LN}}$ and $Q_A^{1\text{LN}}$ follow identically to that of (A.38) – (A.39). For simplicity, we assume that the rates of PD-1 receptor internalisation are identical in the TDLN and the TS, so that

$$\frac{dQ_A^{8\text{LN}}}{dt} = \underbrace{\lambda_{P_D A_1} P_D^{8\text{LN}} A_1^{\text{LN}}}_{\text{formation of } Q_A^{8\text{LN}}} - \underbrace{\lambda_{Q_A} Q_A^{8\text{LN}}}_{\text{dissociation of } Q_A^{8\text{LN}}} - \underbrace{d_{Q_A} Q_A^{8\text{LN}}}_{\text{internalisation}}, \tag{A.53}$$

$$\frac{dQ_A^{1\text{LN}}}{dt} = \underbrace{\lambda_{P_D A_1} P_D^{1\text{LN}} A_1^{\text{LN}}}_{\text{formation of } Q_A^{1\text{LN}}} - \underbrace{\lambda_{Q_A} Q_A^{1\text{LN}}}_{\text{dissociation of } Q_A^{1\text{LN}}} - \underbrace{d_{Q_A} Q_A^{1\text{LN}}}_{\text{internalisation}}. \tag{A.54}$$

##### A.6.3 Equation for Pembrolizumab in the TDLN ($A_1^{\text{LN}}$)

The equation for $A_1^{\text{LN}}$ follows identically to that of (A.41) so that

$$\frac{dA_1^{\text{LN}}}{dt} = \underbrace{\sum_{j=1}^{n} \xi_j f_{\text{pembro}} \delta(t - t_j)}_{\text{infusion}} + \underbrace{\lambda_{Q_A}\left(Q_A^{8\text{LN}} + Q_A^{1\text{LN}}\right)}_{\text{dissociation of } Q_A^{8\text{LN}} \text{ and } Q_A^{1\text{LN}}} - \underbrace{\lambda_{P_D A_1}\left(P_D^{8\text{LN}} + P_D^{1\text{LN}}\right) A_1^{\text{LN}}}_{\text{formation of } Q_A^{8\text{LN}} \text{ and } Q_A^{1\text{LN}}} - \underbrace{d_{A_1} A_1^{\text{LN}}}_{\text{elimination}}. \tag{A.55}$$

##### A.6.4 Equation for Unbound PD-L1 in the TDLN ($P_L^{\text{LN}}$)

We recall that PD-L1 is expressed on the surface of mature DCs, effector CD8+ T cells, effector Th1 cells, and effector Tregs. We denote $\mathcal{Y}$ as the set of PD-L1-expressing cells in the TDLN, so that $\mathcal{Y} := \{D^{\text{LN}}, T_A^8, T_A^1, T_A^r\}$, with $\lambda_{P_L^{\text{LN}} Y}$ denoting the synthesis rate of unbound PD-L1 on the surface of $Y \in \mathcal{Y}$. The equation for $P_L^{\text{LN}}$ follows identically to (A.50) so that

$$\frac{dP_L^{\text{LN}}}{dt} = \underbrace{\sum_{Y \in \mathcal{Y}} \lambda_{P_L^{\text{LN}} Y} Y}_{\text{synthesis}} - \underbrace{d_{P_L} P_L^{\text{LN}}}_{\text{degradation}}. \tag{A.56}$$

##### A.6.5 Equations for the PD-1/PD-L1 Complex in the TDLN ($Q^{8\text{LN}}$ and $Q^{1\text{LN}}$)

For simplicity, we assume that the formation and dissociation rates of the PD-1/PD-L1 complex are identical in the TDLN and the TS. The equations for $Q^{8\text{LN}}$ and $Q^{1\text{LN}}$ follow identically from (A.44) – (A.45) so that

$$Q^{8\text{LN}} = \frac{\lambda_{P_D P_L}}{\lambda_Q} P_D^{8\text{LN}} P_L^{\text{LN}}, \tag{A.57}$$

$$Q^{1\text{LN}} = \frac{\lambda_{P_D P_L}}{\lambda_Q} P_D^{1\text{LN}} P_L^{\text{LN}}. \tag{A.58}$$

We note that throughout the model, the PD-1/PD-L1 complex appears only within an inhibition constant, making its absolute magnitude less important since it always appears as a ratio. One thing to note is that activated CD8+ T cells and Th1 cells also express PD-1 receptors and PD-L1 ligands, and we assume that effector and activated cells express these in equal amounts. However, as discussed in Appendix B.2, the ratio between effector and activated T cells can be assumed to remain roughly constant. Since the PD-1/PD-L1 complex concentration is linearly proportional to the product of PD-L1 concentration and unbound PD-1 receptor concentration, and PD-1/PD-L1-mediated inhibition of T cell proliferation in the TDLN appears only as a ratio, it is sufficient to consider only PD-1, PD-L1, and PD-1/PD-L1 concentrations on effector cells, as this will be appropriately scaled by the corresponding inhibition constants. Furthermore, this also justifies using the PD-1/PD-L1 complex concentration on effector T cells as a proxy for its concentration on activated T cells that have not yet undergone division, given that their ratio to effector cells remains roughly constant and that PD-1/PD-L1-mediated inhibition of T cell activation in the TDLN appears only as a ratio.

#### A.7 Model Reduction via QSSA

We observe from Table 2 that the degradation rates of cytokines and DAMPs are, in general, orders of magnitude larger than those of immune and cancer cells. In particular, IL-2, IFN-$\gamma$, TNF, and TGF-$\beta$ evolve on a very fast timescale, with degradation rates significantly higher than all other species in the model, causing them to equilibrate much more rapidly. As such, we perform a QSSA and reduce the model by setting (A.30) – (2.36) to 0 and solving for $I_2$, $I_\gamma$, $I_\alpha$, and $I_\beta$ in terms of the other parameters and variables in the model. This minimally affects the system's evolution after a very short period of transient behaviour [110], and we justify this by observing that, empirically, the deviation in system trajectories remains negligible for nearby parameter choices. Performing the QSSA leads to

$$\frac{dI_2}{dt} = 0 \implies I_2 = \frac{1}{d_{I_2}}\left(\lambda_{I_2T_8}T_8 + \lambda_{I_2T_1}T_1\right), \tag{A.59}$$

$$\frac{dI_\gamma}{dt} = 0 \implies I_\gamma = \frac{1}{d_{I_\gamma}}\left[\left(\lambda_{I_\gamma T_8}T_8 + \lambda_{I_\gamma T_1}T_1\right)\frac{1}{1+T_r/K_{I_\gamma T_r}} + \lambda_{I_\gamma K}K\right], \tag{A.60}$$

$$\frac{dI_\alpha}{dt} = 0 \implies I_\alpha = \frac{1}{d_{I_\alpha}}\left(\lambda_{I_\alpha T_8}T_8 + \lambda_{I_\alpha T_1}T_1 + \lambda_{I_\alpha M_1}M_1 + \lambda_{I_\alpha K}K\right), \tag{A.61}$$

$$\frac{dI_\beta}{dt} = 0 \implies I_\beta = \frac{1}{d_{I_\beta}}\left(\lambda_{I_\beta C}C + \lambda_{I_\beta T_r}T_r + \lambda_{I_\beta M_2}M_2\right). \tag{A.62}$$

We note that this reduction is valid since the timescale of IFN-$\gamma$, the slowest of the "fast" species, is significantly shorter than the timescales of all "slow" species in the model.
