## Appendix B for "Optimisation of pembrolizumab therapy for de novo metastatic MSI-H/dMMR colorectal cancer using data-driven delay integro-differential equations"

### B Steady States and Initial Conditions

We estimated all steady states and initial conditions under the assumption that pembrolizumab has not been and will not be administered.

#### B.1 Steady States and Initial Conditions for Cells in the TS

Digital cytometry has proved itself to be a powerful technique in characterising immune cell populations from individual patients' bulk tissue transcriptomes without requiring physical cell isolation [1–5]. In particular, RNA-sequencing (RNA-seq) deconvolution of tumour gene expressions has been very useful in determining immune profiles and adjusting treatment accordingly. For all algorithms outlined in the sequel, we aggregate the estimates by taking the median of the relevant non-zero values elementwise and then normalising such that their sums become 1.

To estimate immune cell population proportions in dnmMCRC, we applied multiple algorithms and then synthesised their results to obtain estimates for all cell types in the model. We first used the ImmuCellAI algorithm [6], which estimates the abundance of 24 immune cell types from gene expression data and has also been shown to be highly accurate in predicting immunotherapy response. These immune cell types include 18 T cell subsets, including CD4+ T cells which incorporate T helper cells (namely Th1 cells, Th2 cells, Th17 cells, and T follicular helper cells), regulatory T cells (including natural Tregs (nTregs), induced Tregs (iTregs), and type 1 regulatory T cells (Tr1s)), naive CD4+ T cells (CD4_naive) and other CD4+ T cells (CD4_T). In addition, they include naive CD8+ T cells (CD8_T), CTLs (Tc), exhausted CD8+ T cells (Tex), central memory T cells (Tcm), effector memory T cells (Tem), natural killer T cells (NKT), $\gamma\delta$ T cells (Tgd), and mucosal-associated invariant T cells (MAIT). ImmuCellAI also estimates the abundance of DCs, B cells, monocytes, macrophages, and

NK cells. Direct correspondences between state variables in the model and ImmuCellAI cell types are shown in Table B.1.

Table B.1: Mappings between state variables of the model and ImmuCellAI immune cell types.

| State Variable | ImmuCellAI Cell Type |
|---|---|
| $T_8$ | Tc |
| $T_{\text{ex}}$ | Tex |
| $T_r$ | nTreg |
| $D_0$, $D$ | DC |
| $T_1$ | Th1 |
| $M_0$, $M_1$, $M_2$ | Macrophage |
| $K_0$, $K$ | NK |

Using the UCSC Xena web portal [7], RSEM normalised RNA-seq gene expression profiles of patients from the TCGA COAD and TCGA READ projects [8] were acquired, featuring patients with colorectal adenocarcinomas. Corresponding clinical and biospecimen data were downloaded from the GDC portal [9] and included tumour dimensions, necrotic cell percentage, AJCC TNM stage, and MSIsensor and MANTIS MSI statuses. We filtered for samples from primary tumours and with non-empty necrosis percentage data from patients with AJCC stage IV CRC and at least one of MANTIS score $> 0.4$ or MSIsensor score $> 3.5\%$, as these are the default thresholds for MSI-H [10]. We performed 2-means clustering on the estimated cell proportions generated by ImmuCellAI to categorise samples into the stage IVA and stage IVB/IVC TNM stages. We considered only the 'NK', 'DC', nTreg', 'Th1', 'Cytotoxic', and 'Exhausted' cell types as part of the clustering, excluding macrophages because they consist of M0, M1, and M2 phenotypes, which are not differentiated by ImmuCellAI. To properly assign and label each cluster, we compared the individual coordinates of each cluster's centroid. Notably, stage IVB/IVC samples, corresponding to further CRC progression, are characterised by a higher proportion of DCs and nTregs, along with a lower proportion of Th1 cells, cytotoxic T cells, and NK cells compared to stage IVA samples. In particular, to infer steady states, we used stage IVB/IVC samples, whilst stage IVA samples were used to determine initial conditions. We also used the manually curated TIMEDB cell composition database [11] to source tumour deconvolution estimates for each relevant individual sample.

Additionally, the aggregated estimated cell proportions generated by ImmuCellAI for steady states and initial conditions, after normalisation, are shown in Table B.2 and Table B.3.

Table B.2: TS steady-state cell proportions for the model, derived using RNA-sequencing deconvolution via ImmuCellAI. Values for italicised cell types are used in estimating TS cell populations in the model.

| **Cell Type** | **Proportion** | **Cell Type** | **Proportion** |
|---|---|---|---|
| *DC* | 0.082580 | *nTreg* | 0.007285 |
| B_cell | 0.091844 | iTreg | 0.004596 |
| Monocyte | 0.083978 | *Th1* | 0.002784 |
| *Macrophage* | 0.069823 | Th2 | 0.004640 |
| *NK* | 0.106673 | Th17 | 0.003709 |
| Neutrophil | 0.119640 | Tfh | 0.007416 |
| CD4_T | 0.069093 | CD8_naive | 0.003222 |
| CD8_T | 0.064491 | *Tc* | 0.004640 |
| NKT | 0.153019 | *Tex* | 0.003248 |
| Tgd | 0.097433 | MAIT | 0.004640 |
| CD4_naive | 0.001375 | Tcm | 0.003669 |
| Tr1 | 0.006492 | Tem | 0.003712 |

Table B.3: TS initial condition cell proportions for the model, derived using RNA-sequencing deconvolution via ImmuCellAI. Values for italicised cell types are used in estimating TS cell populations in the model.

| **Cell Type** | **Proportion** | **Cell Type** | **Proportion** |
|---|---|---|---|
| *DC* | 0.059843 | *nTreg* | 0.005280 |
| B_cell | 0.144175 | iTreg | 0.005280 |
| Monocyte | 0.111765 | *Th1* | 0.002640 |
| *Macrophage* | 0.050461 | Th2 | 0.002703 |
| *NK* | 0.128857 | Th17 | 0.002703 |
| Neutrophil | 0.116214 | Tfh | 0.009680 |
| CD4_T | 0.075684 | CD8_naive | 0.005054 |
| CD8_T | 0.081099 | *Tc* | 0.004211 |
| NKT | 0.100022 | *Tex* | 0.005054 |
| Tgd | 0.070285 | MAIT | 0.007580 |
| CD4_naive | 0.002546 | Tcm | 0.002703 |
| Tr1 | 0.006160 | Tem | 0.000000 |

To determine the proportions of $M_0$, $M_1$ and $M_2$, we used the CIBERSORTx algorithm [1], due to its high accuracy [12]. We followed a similar approach to [13] and [14] and applied CIBERSORTx B-mode on the refined gene expression data, using the validated LM22 signature matrix [5], which gave relative immune cell proportions of 22 immune cell types using 547 signature genes derived from microarray data. Direct correspondences between state variables in the model and keys of the LM22 signature matrix are shown in Table B.4.

Table B.4: Mappings between state variables of the model and keys of the LM22 signature matrix.

| **State Variable** | **LM22 key** |
|---|---|
| $D_0$ | Dendritic cells resting |
| $D$ | Dendritic cells activated |
| $M_0$ | Macrophages M0 |
| $M_1$ | Macrophages M1 |
| $M_2$ | Macrophages M2 |
| $K_0$ | NK cells resting |
| $K$ | NK cells activated |

The aggregated estimated cell proportions generated by CIBERSORTx for steady states and initial conditions, after normalisation, are shown in Table B.5 and Table B.6.

Table B.5: TS steady-state cell proportions for the model, derived using RNA-sequencing deconvolution via CIBERSORTx. Values for italicised cell types are used in estimating TS cell populations in the model.

| **Cell Type** | **Proportion** | **Cell Type** | **Proportion** |
|---|---|---|---|
| B cells naive | 0.021581 | NK cells activated | 0.068787 |
| B cells memory | 0.010581 | Monocytes | 0.010005 |
| Plasma cells | 0.012526 | *Macrophages M0* | 0.107696 |
| T cells CD8 | 0.080354 | *Macrophages M1* | 0.044431 |
| T cells CD4 naive | 0.109475 | *Macrophages M2* | 0.176747 |
| T cells CD4 memory resting | 0.146694 | Dendritic cells resting | 0.011121 |
| T cells CD4 memory activated | 0.041974 | Dendritic cells activated | 0.000000 |
| T cells follicular helper | 0.004463 | Mast cells resting | 0.000000 |
| T cells regulatory (Tregs) | 0.030406 | Mast cells activated | 0.088533 |
| T cells gamma delta | 0.000000 | Eosinophils | 0.000000 |
| NK cells resting | 0.000000 | Neutrophils | 0.034625 |

Table B.6: TS initial condition cell proportions for the model, derived using RNA-sequencing deconvolution via CIBERSORTx. Values for italicised cell types are used in estimating TS cell populations in the model.

| **Cell Type** | **Proportion** | **Cell Type** | **Proportion** |
|---|---|---|---|
| B cells naive | 0.043647 | NK cells activated | 0.074405 |
| B cells memory | 0.020216 | Monocytes | 0.033754 |
| Plasma cells | 0.019842 | *Macrophages M0* | 0.047465 |
| T cells CD8 | 0.197846 | *Macrophages M1* | 0.039707 |
| T cells CD4 naive | 0.000000 | *Macrophages M2* | 0.245338 |
| T cells CD4 memory resting | 0.070216 | Dendritic cells resting | 0.038535 |
| T cells CD4 memory activated | 0.000000 | Dendritic cells activated | 0.000000 |
| T cells follicular helper | 0.008138 | Mast cells resting | 0.048794 |
| T cells regulatory (Tregs) | 0.068395 | Mast cells activated | 0.032746 |
| T cells gamma delta | 0.000000 | Eosinophils | 0.010958 |
| NK cells resting | 0.000000 | Neutrophils | 0.000000 |

However, to determine the proportions of $D_0$ and $D$, and $K_0$ and $K$, we could not use CIBERSORTx due to its nil results. Instead, we used a combination of biologically informed assumptions and data from physical experiments. It was determined in [15] that the ratio of the proportions of cytotoxic, activated NK cells to resting NK cells decreases as CRC progresses. We thus assume that, $K_0(0) = 10K(0)$, and that $\overline{K_0} = 20\overline{K}$. We also estimate that $D_0(0) \approx D(0)$, using [16] as a guide, and noting that cancer progression leads to increased DAMP production and DC maturation, assume that $\overline{D} = 2\overline{D_0}$.

We integrated the relative proportions within cell types for DCs, NK cells, and macrophages outputted by CIBERSORTx into the ImmuCellAI abundance estimates. We note that the density of immune cells in a healthy adult colon is approximately $3.37 \times 10^7$ cell/g [17], which assuming a tissue density of 1.03 g/cm$^3$, results in a total immune cell density of $3.47 \times 10^7$ cell/cm$^3$. However, advanced cancer induces lymphadenopathy [18, 19], which [17] estimates results in an increase in the total number of lymphocytes of at most 10%. As such, we assume that there is a 10% increase in lymphocyte concentration in dnmMCRC.

Accounting for the low immunogenicity of tumours in dnmMCRC, we followed [14] and assumed at steady state that the density of cancer cells is equal to double the total immune cell density. Taking into account lymphadenopathy and using data from [17], we assumed that the total immune cell density in dnmMCRC initially is approximately $3.70 \times 10^7$ cell/cm$^3$ and at steady state is approximately $3.69 \times 10^7$ cell/cm$^3$. As such, we are justified in assuming that the total immune cell density remains constant throughout the cancer's progression. From the TCGA biospecimen data, the median necrotic cancer cell percentage for stage IV MSI-H CRC samples is 5%. As such, denoting $\overline{\text{TIC}}$ as the total immune cell density at steady state, and $N_p$ as the necrotic cell percentage, we have that

$$\overline{C} + \overline{N_c} = 2 \times \overline{\text{TIC}}, \tag{B.1}$$

$$\frac{\overline{N_c}}{N_p} = \frac{\overline{C}}{1 - N_p}, \quad \overline{C} = 2 \times \overline{\text{TIC}} \times (1 - N_p) \implies \overline{N_c} = 2 \times \overline{\text{TIC}} \times N_p. \tag{B.2}$$

Thus, at steady state, $C \approx 7.02 \times 10^7$ cell/cm$^3$ and $N_c \approx 3.69 \times 10^6$ cell/cm$^3$.

A retrospective cohort study by Burke et al. considered CRC patients at Leeds Teaching Hospitals NHS Trust over a 2-year interval who received no treatment and who underwent computed tomography (CT) scans twice, more than 5 weeks apart. It was found that in patients whose M category remained stable at M1, the median tumour doubling time for these patients was 213 days [20]. Furthermore, the mean tumour volume in stage IVA and stage IVB CRC patients was found to be 33.23 cm$^3$ and 59.87 cm$^3$, respectively [21]. Noting that fewer than 0.1% of disseminated cancer cells successfully form distant metastases [22], we assumed that the primary tumour volume in dnmMCRC remains close, if not slightly smaller, to these values. This corresponds to $V_{\text{TS}}$ taking approximately 180.9 days to reach its steady-state value, and we assume this to be the case for $C$ and $N_c$ as well. This also corresponds to an initial condition for $C$ being $C(0) = 3.90 \times 10^7$ cell/cm$^3$ and thus $N_c(0) \approx 2.05 \times 10^6$ cell/cm$^3$.

Combining everything, the resultant steady states and initial conditions for the model are shown in Table B.7 and Table B.8.

Table B.7: TS steady-state cell densities for the model, combining estimates derived from ImmuCellAI and CIBERSORTx. All values are in cell/cm$^3$.

| $C$ | $N_c$ | $D_0$ | $D$ | $T_8$ | $T_{\text{ex}}$ | $T_1$ |
|---|---|---|---|---|---|---|
| $7.02 \times 10^7$ | $3.69 \times 10^6$ | $9.55 \times 10^5$ | $1.91 \times 10^6$ | $1.77 \times 10^5$ | $1.24 \times 10^5$ | $1.06 \times 10^5$ |
| $T_r$ | $M_0$ | $M_1$ | $M_2$ | $K_0$ | $K$ | |
| $2.78 \times 10^5$ | $7.93 \times 10^5$ | $3.27 \times 10^5$ | $1.30 \times 10^6$ | $3.88 \times 10^6$ | $1.94 \times 10^5$ | |

Table B.8: TS initial condition cell densities for the model, combining estimates derived from ImmuCellAI and CIBERSORTx. All values are in cell/cm$^3$.

| $C$ | $N_c$ | $D_0$ | $D$ | $T_8$ | $T_{\text{ex}}$ | $T_1$ |
|---|---|---|---|---|---|---|
| $3.90 \times 10^7$ | $2.05 \times 10^6$ | $1.04 \times 10^6$ | $1.04 \times 10^6$ | $1.61 \times 10^5$ | $1.93 \times 10^5$ | $1.01 \times 10^5$ |
| $T_r$ | $M_0$ | $M_1$ | $M_2$ | $K_0$ | $K$ | |
| $2.02 \times 10^5$ | $2.50 \times 10^5$ | $2.09 \times 10^5$ | $1.29 \times 10^6$ | $4.47 \times 10^6$ | $4.47 \times 10^5$ | |

We note that, technically, ImmuCellAI is an enrichment-based method that does not provide absolute immune cell proportions but rather estimates abundances across various immune cell subtypes not reported by CIBERSORTx. However, normalising these abundances provides a good approximation of the true immune cell proportions, thereby allowing ImmuCellAI to be justifiably employed to estimate immune cell steady states and initial conditions.

#### B.2 Steady States and Initial Conditions for Cells in the TDLN

To determine the initial conditions and steady-state values for $T_0^8$, $T_A^8$, $T_0^4$, $T_A^1$, $T_0^r$, and $T_A^r$, we used ImmuCellAI on the GSE26571 dataset from the NCBI Gene Expression Omnibus repository [23, 24], obtaining deconvolution results from TIMEDB. This contains 9 samples of lymph node metastases from 7 patients with colon adenocarcinoma, with data from [25]. However, the dataset's metadata does not contain AJCC TNM stages for patients. To estimate the TNM stages of the patients with lymph node metastases, we considered the samples of lymph node metastases for these patients and applied the ImmuCellAI algorithm to estimate their immune cell abundances, ignoring Tcm and Tem cell subtypes. Furthermore, we denote $T_A^{8\text{LN}}$ and $T_A^{1\text{LN}}$ as the total number of activated CD8+ T cells and activated Th1 cells in the TDLN, respectively, and let $T_r^{\text{LN}}$ denote the total number of Tregs in the TDLN. Mappings between ImmuCellAI immune cell types and TDLN cell types in the model are shown in Table B.9.

Table B.9: Mappings between TDLN cell types in the model and ImmuCellAI immune cell types.

| **State Variable** | **ImmuCellAI Cell Type** |
|---|---|
| $T_0^8$ | CD8_naive |
| $T_A^{8\text{LN}}$ | Tc |
| $T_0^4$ | CD4_naive |
| $T_A^{1\text{LN}}$ | Th1 |
| $T_0^r$, $T_r^{\text{LN}}$ | nTreg |

To distinguish lymph node metastases from stage IVA patients, with those with more advanced disease, we performed 2-means clustering on the estimated cell proportions generated by ImmuCellAI, following a similar approach to Appendix B.1. In particular, we considered the 'nTreg', 'Th1', 'Th2', and 'Cytotoxic' cell types as part of the clustering. We, again, compare the individual coordinates of each cluster's centroid and note that lymph node metastases from stage IVB/IVC samples, which correspond to more advanced CRC progression, exhibit a higher proportion of Th2 cells and nTregs, alongside a lower proportion of Th1 cells and cytotoxic T cells compared to stage IVA samples. Like before, we used lymph node metastases from stage IVB/IVC patients to infer TDLN steady states and those from stage IVA patients to infer initial conditions. Aggregating the estimates, as before, and then normalising such that their sums become 1, results in the proportions as shown in Table B.10 and Table B.11.

Table B.10: TDLN steady-state cell proportions for the model, derived using RNA-sequencing deconvolution via ImmuCellAI. Values for italicised cell types are used in estimating TDLN cell populations in the model.

| **Cell Type** | **Proportion** | **Cell Type** | **Proportion** |
|---|---|---|---|
| DC | 0.145811 | Tr1 | 0.000876 |
| B_cell | 0.107580 | *nTreg* | 0.001742 |
| Monocyte | 0.025104 | iTreg | 0.002588 |
| Macrophage | 0.086242 | *Th1* | 0.007173 |
| NK | 0.028137 | Th2 | 0.023289 |
| Neutrophil | 0.078899 | Th17 | 0.006353 |
| CD4_T | 0.141528 | Tfh | 0.002723 |
| CD8_T | 0.065793 | *CD8_naive* | 0.006270 |
| NKT | 0.153813 | *Tc* | 0.000896 |
| Tgd | 0.105321 | Tex | 0.004445 |
| *CD4_naive* | 0.003631 | MAIT | 0.001786 |

Table B.11: TDLN initial condition cell proportions for the model, derived using RNA-sequencing deconvolution via ImmuCellAI. Values for italicised cell types are used in estimating TDLN cell populations in the model.

| **Cell Type** | **Proportion** | **Cell Type** | **Proportion** |
|---|---|---|---|
| DC | 0.176871 | Tr1 | 0.000930 |
| B_cell | 0.116990 | *nTreg* | 0.000929 |
| Monocyte | 0.056462 | iTreg | 0.003714 |
| Macrophage | 0.035633 | *Th1* | 0.008357 |
| NK | 0.037605 | Th2 | 0.026991 |
| Neutrophil | 0.087545 | Th17 | 0.004162 |
| CD4_T | 0.134744 | Tfh | 0.002777 |
| CD8_T | 0.056912 | *CD8_naive* | 0.006467 |
| NKT | 0.130453 | *Tc* | 0.000927 |
| Tgd | 0.105489 | Tex | 0.003719 |
| *CD4_naive* | 0.002324 | MAIT | 0.000000 |

The density of immune cells in the lymph nodes of an adult is approximately $1.8 \times 10^9$ cell/g [17], which assuming a tissue density of 1.03 g/cm$^3$, results in a total immune cell density of $1.854 \times 10^9$ cell/cm$^3$. We also assumed that in the TDLN, the number of activated CD8+ T cells having undergone $n^8_{\max}$

divisions is roughly half the number that has only undergone $n^8_{\max} - 1$ divisions and so forth, and similarly for Th1 cells. Furthermore, we assumed that initially, and at steady state, 10% of all Tregs are naive. Thus, we assumed that for $i = 1, 8$,

$$T^i_A = \frac{2^{n^i_{\max}}}{2^{n^i_{\max}+1} - 1} T^{i\mathrm{LN}}_A,$$

and that

$$T^r_0 = \frac{T^{\mathrm{LN}}_r}{10}, \quad T^r_A = \frac{9}{10} \frac{2^{n^r_{\max}}}{2^{n^r_{\max}+1} - 1} T^{\mathrm{LN}}_r.$$

Combining everything, and incorporating T cell division numbers as justified in Appendix C.1, the resultant steady-states and initial conditions for the model are shown in Table B.12 and Table B.13, respectively.

Table B.12: TDLN steady-state cell densities for the model, using estimates derived from ImmuCellAI. All values are in cell/cm$^3$.

| $T^8_0$ | $T^8_A$ | $T^4_0$ | $T^1_A$ | $T^r_0$ | $T^r_A$ |
|---|---|---|---|---|---|
| $1.16 \times 10^7$ | $8.31 \times 10^5$ | $6.73 \times 10^6$ | $6.66 \times 10^6$ | $3.23 \times 10^5$ | $1.47 \times 10^6$ |

Table B.13: TDLN initial condition cell densities for the model, using estimates derived from ImmuCellAI. All values are in cell/cm$^3$.

| $T^8_0$ | $T^8_A$ | $T^4_0$ | $T^1_A$ | $T^r_0$ | $T^r_A$ |
|---|---|---|---|---|---|
| $1.20 \times 10^7$ | $8.60 \times 10^5$ | $4.31 \times 10^6$ | $7.76 \times 10^6$ | $1.72 \times 10^5$ | $7.81 \times 10^5$ |

#### B.3 Steady States and Initial Conditions for DAMPs

We note that 1 cm$^3$ = 1 mL for all DAMP measurements. To estimate DAMP steady states and initial conditions, we look at the respective experimental tissue concentration data, noting that this is more accurate than the more widely available serum/plasma concentration data. Nonetheless, we use serum/plasma concentration data, where relevant, to guide estimates if the corresponding tissue concentration data is limited. We note that DAMPs only appear in the model within an inhibition or half-saturation constant, making their absolute magnitude less important since they always appear as a ratio.

##### B.3.1 Estimates for $H$

In [26], a study of blood samples from 144 patients with CRC was conducted, with the serum HMGB1 levels of patients with distant metastasis being $13.32 \pm 6.12$ $\mu$g/L, which was significantly higher than those with only lymphatic metastasis at $10.14 \pm 4.38$ $\mu$g/L. We assumed that the serum concentrations and tissue concentrations of HMGB1 are similar, so we took the initial condition for $H$ to be $1.33 \times 10^{-8}$ g/cm$^3$ and the steady state to be $1.94 \times 10^{-8}$ g/cm$^3$.

##### B.3.2 Estimates for $S$

In epithelial ovarian cancer (EOC), calreticulin concentrations when no drugs are introduced were approximately $2 \times 10^{-2} \pm 2.5 \times 10^{-2}$ $\mu$g/mL [27]. Since surface calreticulin is produced by necrotic cancer cells, which have a larger population at steady state compared to initially, we assume that there is more surface calreticulin at steady state. We assumed that calreticulin concentrations in EOC are similar to those in MSI-H/dMMR CRC, so we assumed an initial condition for $S$ of $3.25 \times 10^{-8}$ g/cm$^3$ and a steady state of $4.5 \times 10^{-8}$ g/cm$^3$.

#### B.4 Steady States and Initial Conditions for Cytokines

To estimate cytokine steady states and initial conditions, we look at the respective experimental tissue concentration data, noting that 1 cm$^3$ = 1 mL for all cytokine measurements. We note that cytokines only appear in the model within an inhibition or half-saturation constant, making their absolute magnitude less important since they always appear as a ratio.

##### B.4.1 Estimates for $I_2$

The tissue concentration of IL-2 in CRC is very low and was found to be below the lower limit of quantification in various experiments [28, 29]. In tumour supernatants of invasive ductal cancer, the median IL-2 concentration was found to be 2.1 pg/mL with the interquartile range being 2.0 pg/mL − 4.9 pg/mL [30]. We assumed similar concentrations of IL-2 in the tissue of CRC patients.

Taking into account the well-documented anti-tumour properties of IL-2 [31, 32] and decreased IL-2 serum concentration in metastatic CRC patients compared to those without distant metastasis [33], we assume that $I_2$ has a steady-state value of $2.00 \times 10^{-12}$ g/cm$^3$.

##### B.4.2 Estimates for $I_\gamma$

It was found in [28] that the maximum tissue concentration of IFN-γ in CRC patients was 49.3 pg/mL, with the upper quartile concentration being approximately 16.9 pg/mL. It was found in [34] that the serum concentration of IFN-γ in stage IV CRC patients (median ≈ 20.75 pg/mL) is significantly higher than that of stage I-III patients (median ≈ 1 pg/mL). We thus set the steady state of $I_\gamma$ to $4.93 \times 10^{-11}$ g/cm$^3$.

##### B.4.3 Estimates for $I_\alpha$

It was found in [34] that in advanced CRC patients, i.e those with stage III or stage IV disease, the mean TNF tissue concentration was ≈ 53 pg/mL, with the concentration one standard deviation above the mean being approximately 90 pg/mL. Furthermore, the serum TNF concentration in stage IV CRC patients (median 20.3 pg/mL) is significantly higher than in stage III CRC patients (median 16.0 pg/mL) [35]. We thus set the steady state of $I_\alpha$ to $9.00 \times 10^{-11}$ g/cm$^3$.

##### B.4.4 Estimates for $I_\beta$

It was found in [36] that in CRC patients, the mean TGF-β tissue concentration was 1311.5 pg/mg, with the concentration one standard error above the mean being 1469.1 pg/mg. Assuming a tissue density of 1.03 g/mL, these correspond to tissue concentrations of $1.35 \times 10^6$ pg/mL and $1.51 \times 10^6$ pg/mL, respectively. Furthermore, the serum TGF-β concentration in stage IV CRC patients (mean 55 pg/mL) is significantly higher than in stage III CRC patients (mean 45 pg/mL) [37]. We thus set the steady state of $I_\beta$ to $1.51 \times 10^{-6}$ g/cm$^3$.

##### B.4.5 Estimates for $I_{10}$

It was found in [34] that in advanced CRC patients, i.e those with stage III or stage IV disease, the mean IL-10 tissue concentration was 115 pg/mL, with the concentration one standard deviation above the mean being approximately 184 pg/mL. Furthermore, the serum IL-10 concentration in stage IV CRC patients (mean 36.02 pg/mL) is significantly higher than in stage III CRC patients (mean 17.07 pg/mL) [38]. We thus set the steady state of $I_{10}$ to $1.84 \times 10^{-10}$ g/cm$^3$, with an initial condition of $1.15 \times 10^{-10}$ g/cm$^3$.
