## Appendix C for "Optimisation of pembrolizumab therapy for de novo metastatic MSI-H/dMMR colorectal cancer using data-driven delay integro-differential equations"

### C Parameter Estimation

We estimate all parameters, where possible, under the assumption that no pembrolizumab has/will be administered. The exception to this is the parameters directly related to pembrolizumab treatment, for which the assumptions are explicitly stated during estimation. Many of the assumptions and techniques in this section are adopted from [1].

#### C.1 TDLN Parameters

##### C.1.1 Estimate for $V_{\mathrm{LN}}$

The mean diameter of lymph nodes in CRC patients where cancer has metastasised was found to be 5.6 mm in [2], with the diameter one standard deviation above the mean being 7.5 mm. Assuming a spherical lymph node, we take the radius of the TDLN to be 3.275 mm, corresponding to $V_{\mathrm{LN}} = \frac{4}{3} \times 3.275^3 \times \pi \text{ mm}^3 = 1.47 \times 10^{-1} \text{ cm}^3$.

##### C.1.2 Estimate for $\tau_m$

In [3], it took 18 hours for DCs, which acquired antigen from a site of subcutaneous injection, to arrive at the lymph node. We assume that this migration time is the same for DCs acquiring cancer antigens from the TS so that $\tau_m = 18 \text{ hr} = 0.75$ day.

##### C.1.3 Estimate for $\tau_a$

To estimate $\tau_a$, we note that T cells in the TDLN travel at speeds of 11 – 14 $\mu$m/min, in comparison to DCs which migrate at speeds of 3 – 6 $\mu$m/min [4]. We thus have that $\tau_a = \frac{4.5}{12.5}\tau_m \approx 0.27$ day.

##### C.1.4 Estimates for CD8+ T cells

Using data from [5], we estimate that naive CD8+ T cells take 2 days to activate, and so set $\tau^8_{\text{act}} = 2$ day. It was found in [6] that activated CD8+ T cells required 39 hours on average to complete their first cell division, and so we set $\Delta^0_8 = 39$ hr $= 1.63$ day. Furthermore, the average division time for subsequent cell cycles is 8.6 hours [6]; however, it can vary between $5-28$ hours. Thus, we set $\Delta_8 = 8.6$ hr $= 0.36$ day. It was shown in [7] that fully activated CD8+ T cells divide a minimum of $7-10$ times; however, they can divide more if persistent antigen exposure is present. Indeed, in Lymphocytic Choriomeningitis Virus (LCV), CD8+ T cells can divide more than 15 times [8]. We perform a compromise and set $n^8_{\max} = 10$. We thus have that $\tau_{T^8_A} = 4.87$ day. Finally, it is widely accepted that T cell exhaustion can arise only days to weeks from the initial antigen exposure in the case of chronic antigen stimulation [9, 10], so that we take $\tau_l = 10$ day.

##### C.1.5 Estimates for Th1 cells

We first note that Th1 cells are phenotypes of CD4+ T helper cells. It was found in [11] that CD4+ T cell priming takes between $1-2$ days, and so we set $\tau^{\text{act}}_4 = 1.5$ day. Compared to CD8+ T cells, CD4+ T cells appear to divide less, with only approximately nine cell divisions as in LCV [12]. We assume this is similar in MSI-H/dMMR CRC, and so set $n^1_{\max} = 9$. It takes between 12 and 24 hours for the first CD4+ T cell division to occur, with subsequent divisions occurring at a rate of approximately 10 hours per cell division [13]. We thus set $\Delta^0_1 = 18.5$ hr $= 0.77$ day, and $\Delta_1 = 10$ hr $= 0.42$ day. This leads to $\tau_{T^1_A} = 4.13$ day.

##### C.1.6 Estimates for Tregs

We assume that the activation of Tregs takes the same amount of time as that of CD4+ T helper cells, so that $\tau^{\text{act}}_r = 1.5$ day. It was found in [14] that in mice, 6 days after tumour implantation, 45% of Tregs in the TDLN had undergone at least 1 division, and 14% had undergone more than six divisions. We thus set $n^r_{\max} = 6$ and assume that the cell division rates of Tregs and CD4+ T helper cells are the same, so that $\Delta^0_r = 0.77$ day and $\Delta_r = 0.42$ day. We thus have that $\tau_{T^r_A} = 2.87$ day.

#### C.2 Half-Saturation Constants

We recall that for some species $X$, $K_X$ is denoted the half-saturation constant of $X$ in a term of the form

$$\frac{X}{K_X + X}.$$

For simplicity, we assume that if $\overline{X}$ denotes the steady-state value of $X$, then

$$\frac{\overline{X}}{K_X + \overline{X}} = \frac{1}{2} \implies K_X = \overline{X}. \tag{C.1}$$

This implies that

$$\begin{aligned} K_{T_8C} &= \overline{C}\tau_l = 7.02 \times 10^8 \ \left(\text{cell/cm}^3\right) \text{ day}, \\ K_{KD_0} &= \overline{D_0} = 9.55 \times 10^5 \text{ cell/cm}^3, \\ K_{KD} &= \overline{D} = 1.91 \times 10^6 \text{ cell/cm}^3, \\ K_{DH} &= \overline{H} = 1.94 \times 10^{-8} \text{ g/cm}^3, \end{aligned}$$

$$K_{DS} = \overline{S} = 4.50 \times 10^{-8}\ \text{g/cm}^3,$$
$$K_{T_8 I_2} = K_{T_1 I_2} = K_{K I_2} = K_{I_{10} I_2} = \overline{I_2} = 2.00 \times 10^{-12}\ \text{g/cm}^3,$$
$$K_{C I_\gamma} = K_{M_1 I_\gamma} = K_{M I_\gamma} = \overline{I_\gamma} = 4.93 \times 10^{-11}\ \text{g/cm}^3,$$
$$K_{C I_\alpha} = K_{M_1 I_\alpha} = K_{M I_\alpha} = \overline{I_\alpha} = 9.00 \times 10^{-11}\ \text{g/cm}^3,$$
$$K_{M_2 I_\beta} = K_{M I_\beta} = \overline{I_\beta} = 1.51 \times 10^{-6}\ \text{g/cm}^3,$$
$$K_{M_2 I_{10}} = \overline{I_{10}} = 1.84 \times 10^{-10}\ \text{g/cm}^3,$$
$$K_{T_1 Q^{T_1}} = \overline{Q^{T_1}} = 6.01 \times 10^{5}\ \text{molec/cm}^3.$$

To estimate $K_{T_{\text{ex}} A_1}$, we note that the value of the geometric mean $C_{\text{avg}}$ of pembrolizumab in serum at steady state varied minimally regardless of whether pembrolizumab was administered at 200 mg every 3 weeks, or 400 mg every 6 weeks [15]. This was equal to approximately 50.8 $\mu$g/mL, and we assumed this to be the same in tissue, so we take $C_{\text{avg}} = 5.08 \times 10^{-5}$ g/cm$^3$ $= 2.05 \times 10^{14}$ molec/cm$^3$, noting that the molecular mass of pembrolizumab is approximately $149,000$ g/mol [16]. Thus, we assume that $K_{T_{\text{ex}} A_1} = 2.05 \times 10^{14}$ molec/cm$^3$.

#### C.3 Inhibition Constants

We recall that for some species $X$, $K_X$ is denoted as the inhibition constant of $X$ in a term of the form

$$\frac{1}{1 + X/K_X}.$$

For simplicity, we assume that if $\overline{X}$ denotes the steady-state value of $X$, then

$$\frac{1}{1 + \overline{X}/K_X} = \frac{1}{2} \implies K_X = \overline{X}. \tag{C.2}$$

This implies that

$$K_{T_8 T_r} = K_{T_1 T_r} = K_{I_\gamma T_r} = \overline{T_r} = 2.78 \times 10^{5}\ \text{cell/cm}^3,$$
$$K_{C I_\beta} = K_{D_0 I_\beta} = K_{K I_\beta} = \overline{I_\beta} = 1.51 \times 10^{-6}\ \text{g/cm}^3,$$
$$K_{T_8 I_{10}} = K_{T_{\text{ex}} I_{10}} = \overline{I_{10}} = 1.84 \times 10^{-10}\ \text{g/cm}^3,$$
$$K_{C Q^{T_8}} = \overline{Q^{T_8}} = 1.35 \times 10^{6}\ \text{molec/cm}^3,$$
$$K_{C Q^{K}} = \overline{Q^{K}} = 2.96 \times 10^{5}\ \text{molec/cm}^3,$$
$$K_{T_0^8 T_A^r} = \tau_8^{\text{act}} \overline{T_A^r} = 2.94 \times 10^{6}\ (\text{cell/cm}^3)\ \text{day},$$
$$K_{T_0^8 Q^{8\text{LN}}} = \tau_8^{\text{act}} \overline{Q^{8\text{LN}}} = 5.84 \times 10^{5}\ (\text{molec/cm}^3)\ \text{day},$$
$$K_{T_A^8 T_A^r} = \tau_{T_A^8} \overline{T_A^r} = 7.16 \times 10^{6}\ (\text{molec/cm}^3)\ \text{day},$$
$$K_{T_A^8 Q^{8\text{LN}}} = \tau_{T_A^8} \overline{Q^{8\text{LN}}} = 1.42 \times 10^{6}\ (\text{molec/cm}^3)\ \text{day},$$
$$K_{T_0^4 T_A^r} = \tau_4^{\text{act}} \overline{T_A^r} = 2.21 \times 10^{6}\ (\text{cell/cm}^3)\ \text{day},$$
$$K_{T_0^4 Q^{1\text{LN}}} = \tau_4^{\text{act}} \overline{Q^{1\text{LN}}} = 2.61 \times 10^{6}\ (\text{molec/cm}^3)\ \text{day},$$
$$K_{T_A^1 T_A^r} = \tau_{T_A^1} \overline{T_A^r} = 6.07 \times 10^{6}\ (\text{molec/cm}^3)\ \text{day},$$
$$K_{T_A^1 Q^{1\text{LN}}} = \tau_{T_A^1} \overline{Q^{1\text{LN}}} = 7.19 \times 10^{6}\ (\text{molec/cm}^3)\ \text{day}.$$

#### C.4 Degradation Rates

We recall the formula that the degradation rate of some species, $X$, is given by

$$d_X = \frac{\ln 2}{t^X_{1/2}} \tag{C.3}$$

where $t^X_{1/2}$ is the half-life of $X$.

##### C.4.1 Estimate for $d_H$

The half-life of HMGB1 was found to be approximately 3 hours in the context of prostate cancer [17]. We assume a similar value for MSI-H/dMMR CRC, and so

$$d_H = \frac{\ln 2}{3 \text{ hr}} = 5.55 \text{ day}^{-1}.$$

##### C.4.2 Estimate for $d_S$

Surface calreticulin has a half-life of approximately 12 hours [18, 19]. Thus, we have that

$$d_S = \frac{\ln 2}{12 \text{ hr}} = 1.39 \text{ day}^{-1}.$$

##### C.4.3 Estimate for $d_{D_0}$

The time taken for immature DCs to degrade is estimated to be 28 days in mice [20]. We assume that this is similarly the case for humans, so that this corresponds to

$$d_{D_0} = \frac{1}{28 \text{ day}} = 3.57 \times 10^{-2} \text{ day}^{-1}.$$

##### C.4.4 Estimate for $d_D$

Mature DCs have a half-life of 1.5 – 2.9 days in mice [21]. We assume that this is similarly the case for humans, and take $t^D_{1/2} = 2.2$ day so that

$$d_D = \frac{\ln 2}{2.2 \text{ day}} = 3.15 \times 10^{-1} \text{ day}^{-1}.$$

##### C.4.5 Estimate for $d_{T_0^8}$

The half-life of naive CD8+ T cells in the lymph node was estimated to be 21.5 days in [22] so that

$$d_{T_0^8} = \frac{\ln 2}{21.5 \text{ day}} = 3.22 \times 10^{-2} \text{ day}^{-1}.$$

##### C.4.6 Estimate for $d_{T_8}$ and $d_{T_{\text{ex}}}$

It was measured in [23] that the mean death rate of circulating CD8+ T cells in HIV seronegative patients was 0.009 day$^{-1}$. We assume that this is the case for MSI-H/dMMR CRC, and so we set $d_{T_8} = d_{T_{\text{ex}}} = 0.009$ day$^{-1}$.

##### C.4.7 Estimate for $d_{T_0^4}$

The half-life of naive CD4+ T cells in the lymph node was estimated to be 17.2 days in [22] so that

$$d_{T_0^4} = \frac{\ln 2}{17.2 \text{ day}} = 4.03 \times 10^{-2} \text{ day}^{-1}.$$

##### C.4.8 Estimate for $d_{T_1}$

It was measured in [23] that the mean death rate of circulating CD4+ T cells in HIV seronegative patients was 0.008 day$^{-1}$. We assume that this is the case for Th1 cells in MSI-H/dMMR CRC, and so we set $d_{T_1} = 0.008$ day$^{-1}$.

##### C.4.9 Estimate for $d_{T_0^r}$

The death rate of naive Tregs in the lymph node was estimated to be $2.2 \times 10^{-3}$ day$^{-1}$ in [24], and we assume that the death rate in MSI-H/dMMR CRC is similar, so that

$$d_{T_0^r} = 2.2 \times 10^{-3} \text{ day}^{-1}.$$

##### C.4.10 Estimate for $d_{T_r}$

The mean half-life of Tregs in healthy adults was measured to be approximately 11 days in [25]. We assume that this is similarly the case for MSI-H/dMMR CRC and that this corresponds to

$$d_{T_r} = \frac{\ln 2}{11 \text{ day}} = 6.30 \times 10^{-2} \text{ day}^{-1}.$$

##### C.4.11 Estimate for $d_{M_0}$

The lifespan for naive macrophages was found in humans to be approximately 1.37 days on average [26]. This corresponds to

$$d_{M_0} = \frac{1}{1.37 \text{ day}} = 0.73 \text{ day}^{-1}.$$

##### C.4.12 Estimate for $d_{M_1}$

The lifespan for M1 macrophages was found in humans to be approximately 1.01 days on average [26]. This corresponds to

$$d_{M_1} = \frac{1}{1.01 \text{ day}} = 0.99 \text{ day}^{-1}.$$

##### C.4.13 Estimate for $d_{M_2}$

The lifespan for M2 macrophages was found in humans to be approximately 7.41 days on average [26]. This corresponds to

$$d_{M_2} = \frac{1}{7.41 \text{ day}} = 1.35 \times 10^{-1} \text{ day}^{-1}.$$

##### C.4.14 Estimate for $d_{K_0}$ and $d_K$

The half-life of human NK cells varies between 1 – 2 weeks [27–29]. We assume that the half-lives of resting and activated NK cells are both equal to 10 days, so that

$$d_{K_0} = d_K = \frac{\ln 2}{10 \text{ day}} = 6.93 \times 10^{-2} \text{ day}^{-1}.$$

##### C.4.15 Estimate for $d_{I_2}$

The half-life of IL-2 varies between 5 – 7 minutes [30]. We take $t_{1/2}^{I_2} = 6.9$ min so that

$$d_{I_2} = \frac{\ln 2}{6.9 \text{ min}} = 1.45 \times 10^{2} \text{ day}^{-1}.$$

##### C.4.16 Estimate for $d_{I_\gamma}$

The half-life of IFN-$\gamma$ varies between 25 – 35 minutes [31]. We take $t_{1/2}^{I_\gamma}$ to be 30 minutes so that

$$d_{I_\gamma} = \frac{\ln 2}{30 \text{ min}} = 3.33 \times 10^{1} \text{ day}^{-1}.$$

##### C.4.17 Estimate for $d_{I_\alpha}$

The half-life of TNF varies between 15 – 30 minutes [32, 33]. We take $t_{1/2}^{I_\alpha}$ to be 18.2 minutes, so that

$$d_{I_\alpha} = \frac{\ln 2}{18.2 \text{ min}} = 5.48 \times 10^{1} \text{ day}^{-1}.$$

##### C.4.18 Estimate for $d_{I_\beta}$

The half-life of active TGF-$\beta$ is approximately 2 – 3 minutes [34]. We take $t_{1/2}^{I_\beta} = 2.5$ min, so that

$$d_{I_\beta} = \frac{\ln 2}{2.5 \text{ min}} = 3.99 \times 10^{2} \text{ day}^{-1}.$$

##### C.4.19 Estimate for $d_{I_{10}}$

The half-life of IL-10 varies between 2.7 – 4.5 hours [35]. We take $t_{1/2}^{I_{10}} = 2.7$ hr so that

$$d_{I_{10}} = \frac{\ln 2}{2.7 \text{ hr}} = 6.16 \text{ day}^{-1}.$$

##### C.4.20 Estimate for $d_{P_D}$

The median lower bound on PD-1 half-life on human peripheral blood mononuclear cells was found to be 49.5 hours based on leucine enrichment in [36]. Hence, we take $t_{1/2}^{P_D} = 49.5$hr so that

$$d_{P_D} = \frac{\ln 2}{49.5 \text{ hr}} = 3.36 \times 10^{-1} \text{ day}^{-1}.$$

##### C.4.21 Estimate for $d_{Q_A}$

The internalisation rate of the PD-1/pembrolizumab complex was estimated to be 0.43 $\text{day}^{-1}$ in [37], and so we estimate $d_{Q_A} = 0.43\ \text{day}^{-1}$.

##### C.4.22 Estimate for $d_{A_1}$

The half-life of pembrolizumab varies between 22 – 27 days [38–40]. We take it to be 23.7 days in consistency with models from Li et al. and Ahamadi et al. [41–43] so that

$$d_{A_1} = \frac{\ln 2}{23.7\ \text{day}} = 2.92 \times 10^{-2}\ \text{day}^{-1}.$$

##### C.4.23 Estimate for $d_{P_L}$

The half-life of fully glycosylated PD-L1 is approximately 12 hours [44], with PD-L1 on immune cells being heavily glycosylated [45]. Thus, we take $t_{1/2}^{P_L} = 12$ hr so that

$$d_{P_L} = \frac{\ln 2}{12\ \text{hr}} = 1.39\ \text{day}^{-1}.$$

#### C.5 DAMP Parameters

##### C.5.1 Estimates for $H$

Considering (2.4) at steady state, we have that

$$\lambda_{HN_c}\overline{N_c} - d_H\overline{H} = 0.$$

This leads to

$$\lambda_{HN_c} = 2.92 \times 10^{-14}\ \ (\text{g/cell})\,\text{day}^{-1}.$$

##### C.5.2 Estimates for $S$

Considering (2.5) at steady state leads to the equation

$$\lambda_{SN_c}\overline{N_c} - d_S\overline{S} = 0.$$

This leads to

$$\lambda_{SN_c} = 1.70 \times 10^{-14}\ \ (\text{g/cell})\,\text{day}^{-1}.$$

#### C.6 Cytokine Production Parameters

To estimate many of the cytokine production constants, we consider (2.30) – (2.38) at steady state and use the data from [46]. For each immune cell, we assume that each cytokine's corresponding gene expression is proportional to its production rate by that cell.

##### C.6.1 Estimates for $I_2$

Using values from [46] and considering (2.30) at steady state, or equivalently considering (2.31), leads to the equations

$$\frac{\lambda_{I_2T_8}}{0.114615876287774} = \frac{\lambda_{I_2T_1}}{0.335763693785869},$$

and

$$\lambda_{I_2T_8}\overline{T_8} + \lambda_{I_2T_1}\overline{T_1} - d_{I_2}\overline{I_2} = 0.$$

Solving these simultaneously leads to

$$\lambda_{I_2T_8} = 5.95 \times 10^{-16} \ \ (\text{g/cell})\,\text{day}^{-1},$$
$$\lambda_{I_2T_1} = 1.74 \times 10^{-15} \ \ (\text{g/cell})\,\text{day}^{-1}.$$

Consequently, considering (2.31), we have that

$$I_2(0) = \frac{1}{d_{I_2}}\left(\lambda_{I_2T_8}T_8(0) + \lambda_{I_2T_1}T_1(0)\right) = 1.87 \times 10^{-12} \ \text{g/cm}^3.$$

##### C.6.2 Estimates for $I_\gamma$

Using values from [46] and considering (2.32) at steady state, or equivalently considering (2.33), leads to the equations

$$\frac{\lambda_{I_\gamma T_8}}{2 \times 0.0539973307184416} = \frac{\lambda_{I_\gamma T_1}}{2 \times 0.0188926732394088} = \lambda_{I_\gamma K},$$

and

$$\left(\lambda_{I_\gamma T_8}\overline{T_8} + \lambda_{I_\gamma T_1}\overline{T_1}\right)\frac{1}{2} + \lambda_{I_\gamma K}\overline{K} - d_{I_\gamma}\overline{I_\gamma} = 0.$$

Solving these simultaneously leads to

$$\lambda_{I_\gamma T_8} = 8.62 \times 10^{-16} \ \ (\text{g/cell})\,\text{day}^{-1},$$
$$\lambda_{I_\gamma T_1} = 3.02 \times 10^{-16} \ \ (\text{g/cell})\,\text{day}^{-1},$$
$$\lambda_{I_\gamma K} = 7.99 \times 10^{-15} \ \ (\text{g/cell})\,\text{day}^{-1}.$$

Consequently, considering (2.33), we have that

$$I_\gamma(0) = \frac{1}{d_{I_\gamma}}\left[\left(\lambda_{I_\gamma T_8}T_8(0) + \lambda_{I_\gamma T_1}T_1(0)\right)\frac{1}{1 + T_r(0)/K_{I_\gamma T_r}} + \lambda_{I_\gamma K}K(0)\right] = 1.10 \times 10^{-10} \ \text{g/cm}^3.$$

##### C.6.3 Estimates for $I_\alpha$

Using values from [46] and considering (2.34) at steady state, or equivalently considering (2.35), leads to the equations

$$\frac{\lambda_{I_\alpha T_8}}{0.0654443776961264} = \frac{\lambda_{I_\alpha T_1}}{0.108187215112606} = \frac{\lambda_{I_\alpha M_1}}{0.0396575742078822} = \frac{\lambda_{I_\alpha K}}{0.114108294134927},$$

and

$$\lambda_{I_\alpha T_8}\overline{T_8} + \lambda_{I_\alpha T_1}\overline{T_1} + \lambda_{I_\alpha M_1}\overline{M_1} + \lambda_{I_\alpha K}\overline{K} - d_{I_\alpha}\overline{I_\alpha} = 0.$$

Solving these simultaneously leads to

$$\begin{aligned}
\lambda_{I_\alpha T_8} &= 5.55 \times 10^{-15} \ \ \text{(g/cell) day}^{-1}, \\
\lambda_{I_\alpha T_1} &= 9.17 \times 10^{-15} \ \ \text{(g/cell) day}^{-1}, \\
\lambda_{I_\alpha M_1} &= 3.36 \times 10^{-15} \ \ \text{(g/cell) day}^{-1}, \\
\lambda_{I_\alpha K} &= 9.68 \times 10^{-15} \ \ \text{(g/cell) day}^{-1}.
\end{aligned}$$

Consequently, considering (2.35), we have that

$$I_\alpha(0) = \frac{1}{d_{I_\alpha}} \left(\lambda_{I_\alpha T_8} T_8(0) + \lambda_{I_\alpha T_1} T_1(0) + \lambda_{I_\alpha M_1} M_1(0) + \lambda_{I_\alpha K} K(0)\right) = 1.25 \times 10^{-10} \ \text{g/cm}^3.$$

##### C.6.4 Estimates for $I_\beta$

Estimating the production constants for TGF-β is slightly more complicated compared to other cytokines. We assume that the results for fibroblastic reticular cells in [47] translate directly to results for cancer-associated fibroblasts (CAFs), which are considered to be all fibroblasts found in the TME [47]. We assume that at steady state, CAFs produce twice as much TGF-β as cancer cells in the TME, and denote the production rate of TGF-β by CAFs as $\lambda_{I_\beta C_F}$. This, in conjunction with values from [46], and considering (2.36) at steady state, or equivalently considering (2.37), leads to the equations

$$\frac{\lambda_{I_\beta C_F}}{2} = \frac{\lambda_{I_\beta C}}{1},$$

and

$$\frac{\lambda_{I_\beta C_F}}{0.175283003265127} = \frac{\lambda_{I_\beta T_r}}{0.507677682409403} = \frac{\lambda_{I_\beta} M_2}{0.63070357154901},$$

and

$$\lambda_{I_\beta C}\overline{C} + \lambda_{I_\beta T_r}\overline{T_r} + \lambda_{I_\beta M_2}\overline{M_2} - d_{I_\beta}\overline{I_\beta} = 0.$$

Solving these simultaneously leads to

$$\begin{aligned}
\lambda_{I_\beta C} &= 7.42 \times 10^{-12} \ \ \text{(g/cell) day}^{-1}, \\
\lambda_{I_\beta T_r} &= 4.30 \times 10^{-11} \ \ \text{(g/cell) day}^{-1}, \\
\lambda_{I_\beta M_2} &= 5.34 \times 10^{-11} \ \ \text{(g/cell) day}^{-1}.
\end{aligned}$$

Consequently, considering (2.37), we have that

$$I_\beta(0) = \frac{1}{d_{I_\beta}} \left(\lambda_{I_\beta C} C(0) + \lambda_{I_\beta T_r} T_r(0) + \lambda_{I_\beta M_2} M_2(0)\right) = 9.20 \times 10^{-7} \ \text{g/cm}^3.$$

##### C.6.5 Estimates for $I_{10}$

Amongst 48 different cell lines tested, it was found in [48] that cancer IL-10 production was maximised in cell lines derived from colon carcinomas. As such, we assume that at steady state, cancer production of IL-10 is equal to half of that by $M_2$ macrophages. We assume that the enhancement factor of IL-2 for IL-10 production by Tregs is similar in CRC to that of inflammatory bowel disease and use the estimate of $\lambda_{I_{10} I_2} = 3$ that was used in [49]. This, in conjunction with values from [46], and considering

(2.38) at steady state leads to the equations

$$\frac{\lambda_{I_{10}C}}{1} = \frac{\lambda_{I_{10}M_2}}{2},$$

and

$$\frac{\lambda_{I_{10}M_2}}{1} = \frac{\lambda_{I_{10}T_r}\left(1 + \frac{\lambda_{I_{10}I_2}}{2}\right)}{0.472157630570674},$$

and

$$\lambda_{I_{10}C}\overline{C} + \lambda_{I_{10}M_2}\overline{M_2} + \lambda_{I_{10}T_r}\left(1 + \frac{\lambda_{I_{10}I_2}}{2}\right)\overline{T_r} - d_{I_{10}}\overline{I_{10}} = 0.$$

Solving these simultaneously leads to

$$\begin{aligned}
\lambda_{I_{10}C} &= 1.55 \times 10^{-17} \ \ (\text{g/cell})\,\text{day}^{-1},\\
\lambda_{I_{10}M_2} &= 3.10 \times 10^{-17} \ \ (\text{g/cell})\,\text{day}^{-1},\\
\lambda_{I_{10}T_r} &= 5.86 \times 10^{-18} \ \ (\text{g/cell})\,\text{day}^{-1}.
\end{aligned}$$

#### C.7 Parameters for DCs, Macrophages, and NK Cells

##### C.7.1 Estimates for $D_0$ and $D$

Adding (2.6) and (2.7) at steady state leads to

$$\mathcal{A}_{D_0} - \frac{\lambda_{D_0K}\overline{D_0K}}{2} - d_{D_0}\overline{D_0} - \lambda_{DD^{\text{LN}}}\overline{D} - d_D\overline{D} = 0.$$

We assume that HMGB1 is the most potent inducer of DC maturation, and as such, at steady state, we assume that

$$\frac{\lambda_{DH}}{2 \times 10} = \frac{\lambda_{DS}}{2 \times 1}.$$

In [50], it was also shown that the percentage of immature DCs that were lysed as a result of NK cells is roughly linear in the ratio of NK cells to immature DCs. When a 1:1 ratio of activated NK cells to immature DCs is present, after 24 hours, roughly 35.5% of immature DCs are lysed, whereas if a 5:1 ratio is present, 85.5% of immature DCs are lysed. At steady state, the ratio of NK cells to immature DCs is $\approx 2.39 : 1$, corresponding to an approximate 52.85% being lysed. However, if we consider only immature DC loss due to degradation, after 24 hours, only $1 - \exp(-d_{D_0}) \approx 3.54\%$ are lost to it. Thus, we assume at steady state that

$$\frac{\lambda_{D_0K}\overline{D_0K}}{2 \times 0.5285} = \frac{d_{D_0}\overline{D_0}}{0.0354} \implies \lambda_{D_0K} = 5.49 \times 10^{-6} \ \left(\text{cell/cm}^3\right)^{-1}\text{day}^{-1}.$$

Considering (2.7) at steady state leads to

$$\frac{\lambda_{DH}\overline{D_0}}{2} + \frac{\lambda_{DS}\overline{D_0}}{2} - \lambda_{DD^{\text{LN}}}\overline{D} - d_D\overline{D} = 0.$$

Finally, it was found in [51] that only a limited number of DCs migrate up to the TDLN, with at most 4% of DCs reaching the TDLN in melanoma patients when DCs were injected intradermally. We assume at steady state that this holds, too, for MSI-H/dMMR CRC. Taking into account that only

$\exp(-d_D\tau_m)$ of mature DCs that leave the TS survive their migration to the TDLN, we have that

$$\frac{\lambda_{DD^{\mathrm{LN}}}}{0.04\exp(d_D\tau_m)} = \frac{d_D}{1-0.04\exp(d_D\tau_m)}.$$

Solving these simultaneously leads to

$$\begin{aligned}
\mathcal{A}_{D_0} &= 1.18\times 10^6\ \left(\text{cell/cm}^3\right)\text{day}^{-1},\\
\lambda_{DH} &= 1.21\ \text{day}^{-1},\\
\lambda_{DS} &= 1.21\times 10^{-1}\ \text{day}^{-1},\\
\lambda_{DD^{\mathrm{LN}}} &= 1.68\times 10^{-2}\ \text{day}^{-1}.
\end{aligned}$$

Considering (2.8) at steady state leads to

$$\frac{\overline{V_{\mathrm{TS}}}}{V_{\mathrm{LN}}}\lambda_{DD^{\mathrm{LN}}}\exp(-d_D\tau_m)\,\overline{D} - d_D\overline{D^{\mathrm{LN}}} = 0 \implies \overline{D^{\mathrm{LN}}} = \frac{\overline{V_{\mathrm{TS}}}\lambda_{DD^{\mathrm{LN}}}\exp(-d_D\tau_m)\,\overline{D}}{V_{\mathrm{LN}}d_D}.$$

We set the initial condition for $D^{\mathrm{LN}}$ to be such that

$$\frac{D^{\mathrm{LN}}(0)}{\overline{D^{\mathrm{LN}}}} = \frac{D(0)}{\overline{D}} \implies D^{\mathrm{LN}}(0) = \overline{D^{\mathrm{LN}}}\frac{D(0)}{\overline{D}}.$$

Thus, solving simultaneously,

$$\begin{aligned}
\overline{D^{\mathrm{LN}}} &= 3.28\times 10^7\ \text{cell/cm}^3,\\
D^{\mathrm{LN}}(0) &= 1.78\times 10^7\ \text{cell/cm}^3.
\end{aligned}$$

##### C.7.2 Estimates for $M_0$, $M_1$, and $M_2$

To estimate the macrophage production constants, we consider (2.25), (2.26), (2.27) at steady state, and use the data from [46]. We assume that the magnitude of response to a specific cytokine is proportional to its corresponding macrophage polarisation rate, where the response is defined as the Euclidean distance between the centroid vectors of cytokine-treated macrophages and phosphate-buffered saline (PBS)-treated macrophages.

Adding (2.25), (2.26), and (2.27) at steady state leads to

$$\mathcal{A}_{M_0} - d_{M_0}\overline{M_0} - d_{M_1}\overline{M_1} - d_{M_2}\overline{M_2} = 0 \implies \mathcal{A}_{M_0} = 1.08\times 10^6\ \left(\text{cell/cm}^3\right)\text{day}^{-1}.$$

Using values from [46], and considering (2.25) at steady state, leads to the equations

$$\begin{aligned}
&\mathcal{A}_{M_0} - \frac{\lambda_{M_1I_\alpha}\overline{M_0}}{2} - \frac{\lambda_{M_1I_\gamma}\overline{M_0}}{2} - \frac{\lambda_{M_2I_{10}}\overline{M_0}}{2} - \frac{\lambda_{M_2I_\beta}\overline{M_0}}{2} - d_{M_0}\overline{M_0} = 0,\\
&\frac{\lambda_{M_1I_\alpha}}{2\times 10.77} = \frac{\lambda_{M_1I_\gamma}}{2\times 12.54} = \frac{\lambda_{M_2I_{10}}}{2\times 6.81} = \frac{\lambda_{M_2I_\beta}}{2\times 7.63}.
\end{aligned}$$

We assume that IFN-$\gamma$ repolarises M2 macrophages to the M1 phenotype slightly more potently than TNF. Hence, at steady state

$$\frac{\lambda_{MI_\gamma}}{2\times 6} = \frac{\lambda_{MI_\alpha}}{2\times 5}.$$

At steady state, we also assume that

$$\frac{1}{\overline{M_2}}\left(\frac{\lambda_{MI_\gamma}}{2}+\frac{\lambda_{MI_\alpha}}{2}\right)=\frac{1}{\overline{M_1}}\frac{\lambda_{MI_\beta}}{2}.$$

Solving these simultaneously leads to

$$\begin{aligned}
\lambda_{M_1I_\alpha} &= 3.59\times 10^{-1}\ \text{day}^{-1},\\
\lambda_{M_1I_\gamma} &= 4.18\times 10^{-1}\ \text{day}^{-1},\\
\lambda_{M_2I_{10}} &= 2.27\times 10^{-1}\ \text{day}^{-1},\\
\lambda_{M_2I_\beta} &= 2.54\times 10^{-1}\ \text{day}^{-1},\\
\lambda_{MI_\gamma} &= 1.39\times 10^{-2}\ \text{day}^{-1},\\
\lambda_{MI_\alpha} &= 1.15\times 10^{-2}\ \text{day}^{-1},\\
\lambda_{MI_\beta} &= 6.39\times 10^{-3}\ \text{day}^{-1}.
\end{aligned}$$

##### C.7.3 Estimates for $K_0$ and $K$

To estimate NK cell production parameters, we do a similar process to macrophages. Adding (2.28) and (2.29) at steady state leads to

$$\mathcal{A}_{K_0}-d_{K_0}\overline{K_0}-d_K\overline{K}=0 \implies \mathcal{A}_{K_0}=2.82\times 10^5\ \left(\text{cell/cm}^3\right)\text{day}^{-1}.$$

Considering (2.29) at steady state leads to

$$\frac{1}{2}\left(\frac{\lambda_{KI_2}\overline{K_0}}{2}+\frac{\lambda_{KD_0}\overline{K_0}}{2}+\frac{\lambda_{KD}\overline{K_0}}{2}\right)-d_K\overline{K}=0.$$

We assume that mature DCs are more potent activators of NK cells than immature DCs, so that at steady state

$$\frac{\lambda_{KD}}{2\times 5}=\frac{\lambda_{KD_0}}{2\times 1}.$$

We finally assume that DC-mediated NK-cell activation is twice as potent as cytokine-induced activation at steady state, so that

$$\frac{\lambda_{KD_0}/2+\lambda_{KD}/2}{2}=\frac{\lambda_{KI_2}/2}{1}.$$

Solving these simultaneously leads to

$$\begin{aligned}
\lambda_{KI_2} &= 4.62\times 10^{-3}\ \text{day}^{-1},\\
\lambda_{KD_0} &= 1.54\times 10^{-3}\ \text{day}^{-1},\\
\lambda_{KD} &= 7.70\times 10^{-3}\ \text{day}^{-1}.
\end{aligned}$$

#### C.8 T Cell Parameters and Estimates

##### C.8.1 Estimates for $T_0^8$, $T_A^8$, $T_8$, and $T_{\text{ex}}$

Considering (2.9) at steady state leads to

$$\mathcal{A}_{T_0^8}-\overline{R^8}-d_{T_0^8}\overline{T_0^8}=0,$$

and in particular,

$$\overline{R^8} = \frac{\lambda_{T_0^8 T_A^8} \exp\left(-d_{T_0^8} \tau_8^{\mathrm{act}}\right) \overline{D^{\mathrm{LN}} T_0^8}}{4}.$$

Considering (2.11) at steady state leads to

$$\frac{2^{n_{\max}^8} \exp\left(-d_{T_0^8} \tau_{T_A^8}\right) \overline{R^8}}{4} - \lambda_{T_A^8 T_8} \overline{T_A^8} - d_{T_8} \overline{T_A^8} = 0.$$

We first consider the case where no pembrolizumab is present. Considering (2.13) and (2.14) at steady state leads to

$$\frac{V_{\mathrm{LN}}}{V_{\mathrm{TS}}} \lambda_{T_A^8 T_8} \exp\left(-d_{T_8} \tau_a\right) \overline{T_A^8} + \frac{\lambda_{T_8 I_2} \overline{T_8}}{4} - \frac{\lambda_{T_8 C} \overline{T_8}}{2} - \frac{d_{T_8} \overline{T_8}}{2} = 0,$$

$$\frac{\lambda_{T_8 C} \overline{T_8}}{2} - \frac{d_{T_{\mathrm{ex}}} \overline{T_{\mathrm{ex}}}}{2} = 0.$$

We assume that at steady state, 95% of positive $T_8$ growth is due to $T_A^8$ migration to the TS, and the other 5% is due to IL-2-induced proliferation. Thus, we have that

$$\frac{V_{\mathrm{LN}}}{V_{\mathrm{TS}}} \frac{\lambda_{T_A^8 T_8} \exp\left(-d_{T_8} \tau_a\right) \overline{T_A^8}}{0.95} = \frac{\lambda_{T_8 I_2} \overline{T_8}/4}{0.05}.$$

To determine $\lambda_{T_{\mathrm{ex}} A_1}$, we assume that when pembrolizumab is present, at steady state, 20% of exhausted CD8+ T cells are reinvigorated. That is, we assume that

$$\frac{\lambda_{T_{\mathrm{ex}} A_1} \overline{T_{\mathrm{ex}}}/2}{0.2} = \frac{d_{T_{\mathrm{ex}}} \overline{T_{\mathrm{ex}}}/2}{0.8}.$$

Solving these equations simultaneously leads to

$$\begin{aligned}
\mathcal{A}_{T_0^8} &= 3.76 \times 10^5 \ \left(\mathrm{cell/cm^3}\right) \mathrm{day}^{-1}, \\
\lambda_{T_0^8 T_A^8} &= 2.73 \times 10^{-11} \ \left(\mathrm{cell/cm^3}\right)^{-1} \mathrm{day}^{-1}, \\
\overline{R^8} &= 2.43 \times 10^3 \ \left(\mathrm{cell/cm^3}\right) \mathrm{day}^{-1}, \\
\lambda_{T_A^8 T_8} &= 6.32 \times 10^{-1} \ \mathrm{day}^{-1}, \\
\lambda_{T_8 I_2} &= 1.53 \times 10^{-3} \ \mathrm{day}^{-1}, \\
\lambda_{T_8 C} &= 6.31 \times 10^{-3} \ \mathrm{day}^{-1}, \\
\lambda_{T_{\mathrm{ex}} A_1} &= 2.25 \times 10^{-3} \ \mathrm{day}^{-1}.
\end{aligned}$$

##### C.8.2 Estimates for $T_0^4$, $T_A^1$, and $T_1$

Considering (2.15) at steady state leads to

$$\mathcal{A}_{T_0^4} - \overline{R^1} - d_{T_0^4} \overline{T_0^4} = 0,$$

where

$$\overline{R^1} = \frac{\lambda_{T_0^4 T_A^1} \exp\left(-d_{T_0^4} \tau_{\mathrm{act}}^4\right) \overline{D^{\mathrm{LN}} T_0^4}}{4}.$$

Considering (2.17) at steady state leads to

$$\frac{2^{n^1_{\max}} \exp\left(-d_{T_0^4}\tau_{T_A^1}\right)\overline{R^1}}{4} - \lambda_{T_A^1 T_1}\overline{T_A^1} - d_{T_1}\overline{T_A^1} = 0.$$

We assume, like for CD8+ T cells, that at steady state, 95% of positive $T_1$ growth is due to $T_A^1$ migration to the TS, and the other 5% is due to IL-2-induced proliferation. Thus, we have that

$$\frac{V_{\text{LN}}}{V_{\text{TS}}}\frac{\lambda_{T_A^8 T_1}\exp\left(-d_{T_1}\tau_a\right)\overline{T_A^1}}{0.95} = \frac{\lambda_{T_1 I_2}\overline{T_1}/4}{0.05}.$$

Based on murine data from [52], we assume that at steady state, 20% of Th1 cells are converted to Tregs. That is, we assume that

$$\frac{\lambda_{T_1 T_r}\overline{T_1}/2}{0.2} = \frac{d_{T_1}\overline{T_1}}{0.8}.$$

Finally, considering (2.19) at steady state leads to

$$\frac{V_{\text{LN}}}{V_{\text{TS}}}\lambda_{T_A^1 T_1}\exp\left(-d_{T_1}\tau_a\right)\overline{T_A^1} + \frac{\lambda_{T_1 I_2}\overline{T_1}}{4} - \frac{\lambda_{T_1 T_r}\overline{T_1}}{2} - d_{T_1}\overline{T_1} = 0.$$

Solving these equations simultaneously leads to

$$\begin{aligned}
\mathcal{A}_{T_0^4} &= 2.76 \times 10^5 \ \left(\text{cell/cm}^3\right)\text{day}^{-1},\\
\lambda_{T_0^4 T_A^1} &= 8.26 \times 10^{-11} \ \left(\text{cell/cm}^3\right)^{-1}\text{day}^{-1},\\
\overline{R^1} &= 4.28 \times 10^3 \ \left(\text{cell/cm}^3\right)\text{day}^{-1},\\
\lambda_{T_A^1 T_1} &= 6.17 \times 10^{-2}\ \text{day}^{-1},\\
\lambda_{T_1 I_2} &= 2.00 \times 10^{-3}\ \text{day}^{-1},\\
\lambda_{T_1 T_r} &= 4.00 \times 10^{-3}\ \text{day}^{-1}.
\end{aligned}$$

##### C.8.3 Estimates for $T_0^r$, $T_A^r$, and $T_r$

Considering (2.20) at steady state leads to

$$\mathcal{A}_{T_0^r} - \overline{R^r} - d_{T_0^r}\overline{T_0^r} = 0,$$

where

$$\overline{R^r} = \lambda_{T_0^r T_A^r}\exp\left(-d_{T_0^r}\tau_{\text{act}}^r\right)\overline{D^{\text{LN}}}\overline{T_0^r}.$$

Considering (2.22) at steady state leads to

$$2^{n^r_{\max}}\exp\left(-d_{T_0^r}\tau_{T_A^r}\right)\overline{R^r} - \lambda_{T_A^r T_r}\overline{T_A^r} - d_{T_r}\overline{T_A^r} = 0.$$

Finally, considering (2.24) at steady state leads to

$$\frac{V_{\text{LN}}}{V_{\text{TS}}}\lambda_{T_A^r T_r}\exp\left(-d_{T_r}\tau_a\right)\overline{T_A^r} + \frac{\lambda_{T_1 T_r}\overline{T_1}}{2} - d_{T_r}\overline{T_r} = 0.$$

Solving these equations simultaneously leads to

$$
\begin{aligned}
\mathcal{A}_{T_0^r} &= 1.15 \times 10^5 \ \left(\text{cell/cm}^3\right) \text{day}^{-1}, \\
\lambda_{T_0^r T_A^r} &= 1.08 \times 10^{-8} \ \left(\text{cell/cm}^3\right)^{-1} \text{day}^{-1}, \\
\overline{R^r} &= 1.14 \times 10^5 \ \left(\text{cell/cm}^3\right) \text{day}^{-1}, \\
\lambda_{T_A^r T_r} &= 4.88 \ \text{day}^{-1}.
\end{aligned}
$$

#### C.9 Cancer Cell Parameters

##### C.9.1 Estimates for $C$

Considering (2.1) at steady state leads to

$$
\lambda_C \left(1 - \frac{\overline{C}}{C_0}\right) - \frac{\lambda_{CT_8}}{4}\overline{T_8} - \frac{\lambda_{CK}}{4}\overline{K} - \frac{\lambda_{CI_\alpha}}{2} - \frac{\lambda_{CI_\gamma}}{2} = 0.
$$

We assume that CD8+ T cells and NK cells kill cancer cells with similar potency, so we approximate

$$
\lambda_{CK}/4 = \lambda_{CT_8}/4 \implies \lambda_{CK} = \lambda_{CT_8}.
$$

We also assume that the rate that TNF induces tumour necroptosis is larger than that for IFN-$\gamma$, so we approximate

$$
\frac{\lambda_{CI_\alpha}}{2 \times 5} = \frac{\lambda_{CI_\gamma}}{2}.
$$

Solving these simultaneously leads to

$$
\begin{aligned}
\lambda_{CK} &= \lambda_{CT_8}, \\
\lambda_{CI_\alpha} &= \left(\frac{5}{3} - \frac{1.17 \times 10^8}{C_0}\right)\lambda_C - \frac{463750}{3}\lambda_{CT_8}, \\
\lambda_{CI_\gamma} &= \left(\frac{1}{3} - \frac{2.34 \times 10^7}{C_0}\right)\lambda_C - \frac{92750}{3}\lambda_{CT_8}.
\end{aligned}
$$

##### C.9.2 Estimates for $N_c$

Considering (2.2) at steady state leads to the equation

$$
\frac{\lambda_{CI_\alpha}\overline{C}}{2} + \frac{\lambda_{CI_\gamma}\overline{C}}{2} - d_{N_c}\overline{N_c} = 0.
$$

This leads to

$$
d_{N_c} = \frac{1}{\overline{N_c}}\left[\left(1 - \frac{7.02 \times 10^7}{C_0}\right)\lambda_C - 92750\lambda_{CT_8}\right].
$$

##### C.9.3 Fitting $\lambda_C$, $\lambda_{CT_8}$, and $C_0$

We fit $\lambda_C$, $\lambda_{CT_8}$, and $C_0$ by choosing the values such that the steady-state value of $C$ and $N_c$ is reached at 180.9 days, in particular ensuring that $C$ and $N_c$ reach steady state at exactly 180.9 days. Furthermore, we expect monotonicity in the growth of the total cancer population ($C + N_c$) as the cancer progresses without treatment, and so we aim to minimise

$$\text{Objective} = \max\left(\frac{|C(180.9)-\overline{C}|}{\overline{C}}, \frac{|N_c(180.9)-\overline{N_c}|}{\overline{N_c}}, \frac{|C(180.9)+N_c(180.9)-(\overline{C}+\overline{N_c})|}{\overline{C}+\overline{N_c}}\right), \quad \text{(C.4)}$$

subject to

$$\max_{t\in[0,180.9]}(C(t)+N_c(t)) \leq \overline{C}+\overline{N_c}. \quad \text{(C.5)}$$

We perform a parameter sweep to minimise (C.4) subject to (C.5), and set the parameter space to be $\lambda_C \in (0 \text{ day}^{-1}, 2 \text{ day}^{-1}]$, $\lambda_{CT_8} \in (0 \text{ day}^{-1}, 1\times 10^{-6} \text{ day}^{-1}]$, and $C_0 \in (8\times 10^7 \text{ cell/cm}^3, 10^{11} \text{ cell/cm}^3]$, ensuring that all model parameters are positive. The optimal values of $\lambda_C$, $\lambda_{CT_8}$, and $C_0$ were found to be

$$\begin{aligned}\lambda_C &= 5.25\times 10^{-2} \text{ day}^{-1},\\ \lambda_{CT_8} &= 8.01\times 10^{-8} \left(\text{cell/cm}^3\right)^{-1} \text{day}^{-1},\\ C_0 &= 8.89\times 10^7 \text{ cell/cm}^3,\end{aligned}$$

which implies that

$$\begin{aligned}\lambda_{CK} &= 8.01\times 10^{-8} \left(\text{cell/cm}^3\right)^{-1} \text{day}^{-1},\\ \lambda_{CI_\alpha} &= 6.02\times 10^{-3} \text{ day}^{-1},\\ \lambda_{CI_\gamma} &= 1.20\times 10^{-3} \text{ day}^{-1},\\ d_{N_c} &= 6.88\times 10^{-2} \text{ day}^{-1}.\end{aligned}$$

##### C.9.4 Estimates for $V_{\text{TS}}$

We know that $C = f_C V_{\text{TS}}$, and $N_c = f_{N_c} V_{\text{TS}}$. Considering this at steady state leads to

$$\begin{aligned}f_C &= \frac{\overline{C}}{\overline{V_{\text{TS}}}},\\ f_{N_c} &= \frac{\overline{N_c}}{\overline{V_{\text{TS}}}}.\end{aligned}$$

Substituting in values from Appendix B.1 leads to

$$\begin{aligned}f_C &= 1.17\times 10^6 \text{ cell/(cm}^3)^2,\\ f_{N_c} &= 6.16\times 10^4 \text{ cell/(cm}^3)^2.\end{aligned}$$

#### C.10 Estimates for Immune Checkpoint-Associated Components in the TS

##### C.10.1 Estimate for $\lambda_Q$

The dissociation rate of the PD-1/PD-L1 complex was found to be 1.44 $\text{sec}^{-1}$ in [53]. Thus, we have that

$$\lambda_Q = 60\times 60\times 24\times 1.44 \text{ sec}^{-1} = 1.24\times 10^5 \text{ day}^{-1}.$$

##### C.10.2 Estimate for $\lambda_{P_D P_L}$

The formation rate of the PD-1/PD-L1 complex was found to be $1.84 \times 10^5$ $M^{-1}sec^{-1}$ in [53]. To convert this to units of $(\text{molec/cm}^3)^{-1}\text{day}^{-1}$, we recall that 1 M = 1 mol/L = $10^{-3}$ mol/cm$^3$ = $6.022 \times 10^{20}$ molec/cm$^3$. As such,

$$\lambda_{P_D P_L} = 60 \times 60 \times 24 \times 1.84 \times 10^5 \times (6.022 \times 10^{20})^{-1} = 2.64 \times 10^{-11}\ (\text{molec/cm}^3)^{-1}\text{day}^{-1}.$$

##### C.10.3 Estimates for Synthesis Rates and Steady States

We note that estimating parameters, steady states, and initial conditions for PD-1, PD-L1, and the PD-1/PD-L1 complex in the TS is more involved than the previous estimations and requires more information.

We first denote $\rho_{P_D^{T_8}}$, $\rho_{P_D^{T_1}}$, and $\rho_{P_D^K}$ as the number of PD-1 molecules expressed on the surface of CD8+ T cells, Th1 cells, and activated NK cells in the TS, respectively. To determine these parameters, we used the baseline data collected in [54] on 5 advanced cancer patients before their pembrolizumab infusions. The net number of PD-1 molecules on the surface of CD4+ T cells was 2053 molec/cell, and so we set $\rho_{P_D^{T_1}} = 2.05 \times 10^3$ molec/cell. The net number of PD-1 molecules on the surface of CD8+ T cells was 2761 molec/cell, and so we set $\rho_{P_D^{T_8}} = 2.76 \times 10^3$ molec/cell. Despite the net number of PD-1 molecules on the surface of NK cells being below the lower limit of quantification in [54], NK cells substantially express PD-1 [55] in CRC, and so we set $\rho_{P_D^K} = \rho_{P_D^{T_8}}/5 = 5.52 \times 10^2$ molec/cell.

We next denote $\rho_{P_L X}$ as the number of PD-L1 molecules expressed on $X \in \mathcal{X}$, recalling that $\mathcal{X} = \{C, D, T_8, T_1, T_r, M_2\}$. It was found in [53] that the PD-L1 expression on activated CD3+ PD-L1+ T cells was 9282 molec/cell, whilst the PD-L1 expression on mature DCs was 80,372 molec/cell. However, amongst advanced CRC patients, only 22.4% of CD4+ T cells were PD-L1+, and only 16.1% of CD8+ T cells were PD-L1+ [56]. Moreover, only 22% of colonic DCs were PD-L1+ in [57]. We thus assumed that $\rho_{P_L T_1} = \rho_{P_L T_r} = 2.08 \times 10^3$ molec/cell, $\rho_{P_L T_8} = 1.49 \times 10^3$ molec/cell, and $\rho_{P_L D} = 1.77 \times 10^4$ molec/cell. In their quantitative systems pharmacology model of colorectal cancer, Anbari et al. estimated the baseline numbers of PD-L1 molecules per cancer cell and per APC to be 180,000 molec/cell and 266,666 molec/cell, respectively [58]. This makes sense, noting that PDL1 expression in macrophages is stronger and more continuous than that in cancer cells [59]. As such, we set $\rho_{P_L C} = 1.8 \times 10^5$ molec/cell and $\rho_{P_L M_2} = 2.67 \times 10^5$ molec/cell.

Considering (2.39) – (2.41), and (2.46) – (2.49) at steady state in the absence of pembrolizumab leads to

$$\begin{aligned}
\lambda_{P_D^{T_8}} \overline{T_8} - d_{P_D} \overline{P_D^{T_8}} &= 0, \\
\lambda_{P_D^{T_1}} \overline{T_1} - d_{P_D} \overline{P_D^{T_1}} &= 0, \\
\lambda_{P_D^K} \overline{K} - d_{P_D} \overline{P_D^K} &= 0, \\
\sum_{X \in \mathcal{X}} \lambda_{P_L X} \overline{X} - d_{P_L} \overline{P_L} &= 0, \\
\overline{Q^{T_8}} - \frac{\lambda_{P_D P_L}}{\lambda_Q} \overline{P_D^{T_8}}\,\overline{P_L} &= 0,
\end{aligned}$$

$$\overline{Q^{T_1}} - \frac{\lambda_{P_D P_L}}{\lambda_Q}\overline{P_D^{T_1}}\,\overline{P_L} = 0,$$

$$\overline{Q^{K}} - \frac{\lambda_{P_D P_L}}{\lambda_Q}\overline{P_D^{K}}\,\overline{P_L} = 0.$$

By considering the total number of PD-1 receptors expressed on each PD-1-expressing cell at steady state, we expect in the absence of pembrolizumab that

$$\overline{P_D^{T_8}} + \overline{Q^{T_8}} = \rho_{P_D^{T_8}}\overline{T_8},$$

$$\overline{P_D^{T_1}} + \overline{Q^{T_1}} = \rho_{P_D^{T_1}}\overline{T_1},$$

$$\overline{P_D^{K}} + \overline{Q^{K}} = \rho_{P_D^{K}}\overline{K}.$$

We can also consider the total number of PD-L1 ligands at steady state so that

$$\overline{P_L} + \overline{Q^{T_8}} + \overline{Q^{T_1}} + \overline{Q^{K}} = \sum_{X\in\mathcal{X}} \rho_{P_L X}\overline{X}.$$

Finally, we expect the synthesis rates of PD-1 and PD-L1 to be proportional to the total number of PD-1 and PD-L1 molecules expressed per PD-1- and PD-L1-expressing cell, respectively, so that

$$\frac{\lambda_{P_D^{T_8}}}{\rho_{P_D^{T_8}}} = \frac{\lambda_{P_D^{T_1}}}{\rho_{P_D^{T_1}}} = \frac{\lambda_{P_D^{K}}}{\rho_{P_D^{K}}},$$

$$\frac{\lambda_{P_L C}}{\rho_{P_L C}} = \frac{\lambda_{P_L D}}{\rho_{P_L D}} = \frac{\lambda_{P_L T_8}}{\rho_{P_L T_8}} = \frac{\lambda_{P_L T_1}}{\rho_{P_L T_1}} = \frac{\lambda_{P_L T_r}}{\rho_{P_L T_r}} = \frac{\lambda_{P_L M_2}}{\rho_{P_L M_2}}.$$

Solving these simultaneously and ensuring all model parameters are positive leads to

$$\lambda_{P_D^{T_8}} = 9.25\times 10^{2}\ \text{(molec/cell) day}^{-1},$$

$$\lambda_{P_D^{T_1}} = 6.87\times 10^{2}\ \text{(molec/cell) day}^{-1},$$

$$\lambda_{P_D^{K}} = 1.85\times 10^{2}\ \text{(molec/cell) day}^{-1},$$

$$\lambda_{P_L C} = 2.50\times 10^{5}\ \text{(molec/cell) day}^{-1},$$

$$\lambda_{P_L D} = 2.46\times 10^{4}\ \text{(molec/cell) day}^{-1},$$

$$\lambda_{P_L T_8} = 2.07\times 10^{3}\ \text{(molec/cell) day}^{-1},$$

$$\lambda_{P_L T_1} = 2.89\times 10^{3}\ \text{(molec/cell) day}^{-1},$$

$$\lambda_{P_L T_r} = 2.89\times 10^{3}\ \text{(molec/cell) day}^{-1},$$

$$\lambda_{P_L M_2} = 3.71\times 10^{5}\ \text{(molec/cell) day}^{-1}.$$

This leads to

$$\overline{P_D^{T_8}} = 4.87\times 10^{8}\ \text{molec/cm}^3,$$

$$\overline{P_D^{T_1}} = 2.17\times 10^{8}\ \text{molec/cm}^3,$$

$$\overline{P_D^{K}} = 1.07\times 10^{8}\ \text{molec/cm}^3,$$

$$\overline{P_L} = 1.30\times 10^{13}\ \text{molec/cm}^3,$$

$$\overline{Q^{T_8}} = 1.35 \times 10^6 \text{ molec/cm}^3,$$
$$\overline{Q^{T_1}} = 6.01 \times 10^5 \text{ molec/cm}^3,$$
$$\overline{Q^{K}} = 2.96 \times 10^5 \text{ molec/cm}^3.$$

##### C.10.4 Estimates for Initial Conditions

To determine the relevant initial conditions, we can simply consider the total number of PD-1 receptors on each PD-1-expressing cell and PD-L1 ligands in the absence of pembrolizumab so that

$$\begin{aligned}
P_D^{T_8}(0) + Q^{T_8}(0) &= \rho_{P_D^{T_8}} T_8(0), \\
P_D^{T_1}(0) + Q^{T_1}(0) &= \rho_{P_D^{T_1}} T_1(0), \\
P_D^{K}(0) + Q^{K}(0) &= \rho_{P_D^{K}} K(0), \\
P_L(0) + Q^{T_8}(0) + Q^{T_1}(0) + Q^{K}(0) &= \sum_{X \in \mathcal{X}} \rho_{P_L X} X(0).
\end{aligned}$$

We can also consider (2.47) – (2.49) initially, so that

$$\begin{aligned}
Q^{T_8}(0) - \frac{\lambda_{P_D P_L}}{\lambda_Q} P_D^{T_8}(0) P_L(0) &= 0, \\
Q^{T_1}(0) - \frac{\lambda_{P_D P_L}}{\lambda_Q} P_D^{T_1}(0) P_L(0) &= 0, \\
Q^{K}(0) - \frac{\lambda_{P_D P_L}}{\lambda_Q} P_D^{K}(0) P_L(0) &= 0.
\end{aligned}$$

Solving these simultaneously leads to

$$\begin{aligned}
P_D^{T_8}(0) &= 4.44 \times 10^8 \text{ molec/cm}^3, \\
P_D^{T_1}(0) &= 2.07 \times 10^8 \text{ molec/cm}^3, \\
P_D^{K}(0) &= 2.46 \times 10^8 \text{ molec/cm}^3, \\
P_L(0) &= 7.40 \times 10^{12} \text{ molec/cm}^3, \\
Q^{T_8}(0) &= 6.99 \times 10^5 \text{ molec/cm}^3, \\
Q^{T_1}(0) &= 3.26 \times 10^5 \text{ molec/cm}^3, \\
Q^{K}(0) &= 3.88 \times 10^5 \text{ molec/cm}^3.
\end{aligned}$$

We note that excluding bound PD-1 receptors when considering the total number of PD-1 receptors on PD-1-expressing cells does not affect the parameter estimates, steady states, or initial conditions at this level of precision, since the number of unbound PD-1 receptors is several orders of magnitude larger than the number of bound PD-1 receptors on PD-1-expressing cells. Furthermore, this also applies when considering the total number of PD-L1 ligands.

#### C.11 Estimates for Immune Checkpoint-Associated Components in the TDLN

##### C.11.1 Estimates for Synthesis Rates and Steady States

For simplicity, we assume that the total number of PD-1 receptors and PD-L1 ligands on cells in the TDLN is equal to the number on the corresponding cells in the TS. Thus, denoting $\rho_{P_D^{8\text{LN}}}$ and $\rho_{P_D^{1\text{LN}}}$ as the number of PD-1 molecules expressed on the surface of CD8+ T cells and Th1 cells in the TDLN, respectively, we have that $\rho_{P_D^{8\text{LN}}} = \rho_{P_D^{T_8}}$ and $\rho_{P_D^{1\text{LN}}} = \rho_{P_D^{T_1}}$. Similarly, we have that $\lambda_{P_L^{\text{LN}} D^{\text{LN}}} = \lambda_{P_L D}$, $\lambda_{P_L^{\text{LN}} T_A^8} = \lambda_{P_L T_8}$, $\lambda_{P_L^{\text{LN}} T_A^1} = \lambda_{P_L T_1}$, and $\lambda_{P_L^{\text{LN}} T_A^r} = \lambda_{P_L T_r}$ where $\lambda_{P_L^{\text{LN}} D^{\text{LN}}}$, $\lambda_{P_L^{\text{LN}} T_A^8}$, $\lambda_{P_L^{\text{LN}} T_A^1}$, and $\lambda_{P_L^{\text{LN}} T_A^r}$ denote the number of PD-L1 ligands expressed on the surfaces of mature DCs, effector CD8$^+$ T cells, effector Th1 cells, and effector Tregs in the TDLN, respectively. We recall that the set of PD-L1-expressing cells in the TDLN is $\mathcal{Y} = \left\{D^{\text{LN}}, T_A^8, T_A^1, T_A^r\right\}$. The procedure for estimating parameters, steady states, and initial conditions for PD-1, PD-L1, and the PD-1/PD-L1 complex in the TDLN is the same as in the TS. Considering (2.50) – (2.51) and (2.55) – (2.57) at steady state in the absence of pembrolizumab, and making the same assumptions for estimation as in the TS, we obtain

$$
\begin{aligned}
\lambda_{P_D^{8\text{LN}}} \overline{P_D^{8\text{LN}}} - d_{P_D} \overline{P_D^{8\text{LN}}} &= 0, \\
\lambda_{P_D^{1\text{LN}}} \overline{P_D^{1\text{LN}}} - d_{P_D} \overline{P_D^{1\text{LN}}} &= 0, \\
\sum_{Y \in \mathcal{Y}} \lambda_{P_L^{\text{LN}} Y} \overline{Y} - d_{P_L} \overline{P_L^{\text{LN}}} &= 0, \\
\overline{Q^{8\text{LN}}} - \frac{\lambda_{P_D P_L}}{\lambda_Q} \overline{P_D^{8\text{LN}}}\, \overline{P_L^{\text{LN}}} &= 0, \\
\overline{Q^{1\text{LN}}} - \frac{\lambda_{P_D P_L}}{\lambda_Q} \overline{P_D^{1\text{LN}}}\, \overline{P_L^{\text{LN}}} &= 0, \\
\overline{P_D^{8\text{LN}}} + \overline{Q^{8\text{LN}}} &= \rho_{P_D^{8\text{LN}}} \overline{T_A^8}, \\
\overline{P_D^{1\text{LN}}} + \overline{Q^{1\text{LN}}} &= \rho_{P_D^{1\text{LN}}} \overline{T_A^1}, \\
\overline{P_L^{\text{LN}}} + \overline{Q^{8\text{LN}}} + \overline{Q^{1\text{LN}}} &= \sum_{Y \in \mathcal{Y}} \rho_{P_L^{\text{LN}} Y} \overline{Y}, \\
\frac{\lambda_{P_D^{8\text{LN}}}}{\rho_{P_D^{8\text{LN}}}} &= \frac{\lambda_{P_D^{1\text{LN}}}}{\rho_{P_D^{1\text{LN}}}}, \\
\frac{\lambda_{P_L D^{\text{LN}}}}{\rho_{P_L D^{\text{LN}}}} = \frac{\lambda_{P_L T_A^8}}{\rho_{P_L T_A^8}} = \frac{\lambda_{P_L T_A^1}}{\rho_{P_L T_A^1}} &= \frac{\lambda_{P_L T_A^r}}{\rho_{P_L T_A^r}}.
\end{aligned}
$$

Solving these simultaneously and ensuring all model parameters are positive leads to

$$
\begin{aligned}
\lambda_{P_D^{8\text{LN}}} &= 9.27 \times 10^2 \ \ (\text{molec/cell})\, \text{day}^{-1}, \\
\lambda_{P_D^{1\text{LN}}} &= 6.89 \times 10^2 \ \ (\text{molec/cell})\, \text{day}^{-1}, \\
\lambda_{P_L^{\text{LN}} D^{\text{LN}}} &= 2.46 \times 10^4 \ \ (\text{molec/cell})\, \text{day}^{-1}, \\
\lambda_{P_L^{\text{LN}} T_A^8} &= 2.07 \times 10^3 \ \ (\text{molec/cell})\, \text{day}^{-1}, \\
\lambda_{P_L^{\text{LN}} T_A^1} &= 2.89 \times 10^3 \ \ (\text{molec/cell})\, \text{day}^{-1}, \\
\lambda_{P_L^{\text{LN}} T_A^r} &= 2.89 \times 10^3 \ \ (\text{molec/cell})\, \text{day}^{-1}.
\end{aligned}
$$

This leads to

$$
\begin{aligned}
\overline{P_D^{8\mathrm{LN}}} &= 2.29 \times 10^{9} \text{ molec/cm}^3, \\
\overline{P_D^{1\mathrm{LN}}} &= 1.37 \times 10^{10} \text{ molec/cm}^3, \\
\overline{P_L^{\mathrm{LN}}} &= 5.99 \times 10^{11} \text{ molec/cm}^3, \\
\overline{Q^{8\mathrm{LN}}} &= 2.92 \times 10^{5} \text{ molec/cm}^3, \\
\overline{Q^{1\mathrm{LN}}} &= 1.74 \times 10^{6} \text{ molec/cm}^3.
\end{aligned}
$$

##### C.11.2 Estimates for Initial Conditions

To determine the relevant immune checkpoint initial conditions, we can simply consider the total number of PD-1 receptors on each PD-1-expressing cell and the PD-L1 ligands in the absence of pembrolizumab, so that

$$
\begin{aligned}
P_D^{8\mathrm{LN}}(0) + Q^{8\mathrm{LN}}(0) &= \rho_{P_D^{8\mathrm{LN}}} T_A^8(0), \\
P_D^{1\mathrm{LN}}(0) + Q^{1\mathrm{LN}}(0) &= \rho_{P_D^{1\mathrm{LN}}} T_A^1(0), \\
P_L^{\mathrm{LN}}(0) + Q^{8\mathrm{LN}}(0) + Q^{1\mathrm{LN}}(0) &= \sum_{Y \in \mathcal{Y}} \rho_{P_L Y} Y(0).
\end{aligned}
$$

We can also consider (2.56) and (2.57) initially, so that

$$
\begin{aligned}
Q^{8\mathrm{LN}}(0) - \frac{\lambda_{P_D P_L}}{\lambda_Q} P_D^{8\mathrm{LN}}(0) P_L^{\mathrm{LN}}(0) &= 0, \\
Q^{1\mathrm{LN}}(0) - \frac{\lambda_{P_D P_L}}{\lambda_Q} P_D^{1\mathrm{LN}}(0) P_L^{\mathrm{LN}}(0) &= 0.
\end{aligned}
$$

Solving these simultaneously leads to

$$
\begin{aligned}
P_D^{8\mathrm{LN}}(0) &= 2.37 \times 10^{9} \text{ molec/cm}^3, \\
P_D^{1\mathrm{LN}}(0) &= 1.59 \times 10^{10} \text{ molec/cm}^3, \\
P_L^{\mathrm{LN}}(0) &= 3.34 \times 10^{11} \text{ molec/cm}^3, \\
Q^{8\mathrm{LN}}(0) &= 1.69 \times 10^{5} \text{ molec/cm}^3, \\
Q^{1\mathrm{LN}}(0) &= 1.13 \times 10^{6} \text{ molec/cm}^3.
\end{aligned}
$$

We note again that excluding bound PD-1 receptors when considering the total number of PD-1 receptors on PD-1-expressing cells does not affect the parameter estimates, steady states, or initial conditions at this level of precision, since the number of unbound PD-1 receptors is several orders of magnitude larger than the number of bound PD-1 receptors on PD-1-expressing cells. Furthermore, this also applies when considering the total number of PD-L1 ligands.

#### C.12 Estimates for $A_1$ and $A_1^{\mathrm{LN}}$

##### C.12.1 Estimate for $f_{\mathrm{pembro}}$

To determine $f_{\mathrm{pembro}}$, we use the formula

$$f_{\mathrm{pembro}} = \frac{C_{\mathrm{max,ss}}(\xi_{\mathrm{pembro}}) - C_{\mathrm{min,ss}}(\xi_{\mathrm{pembro}})}{\xi_{\mathrm{pembro}}}, \tag{C.6}$$

where $C_{\mathrm{max,ss}}/C_{\mathrm{min,ss}}$ corresponds to the maximum and minimum serum concentration of pembrolizumab at steady state after a dose, $\xi_{\mathrm{pembro}}$, of pembrolizumab is administered, respectively.

For pembrolizumab, the mean $C_{\mathrm{min,ss}}/C_{\mathrm{max,ss}}$ was found to be approximately 32.6/85.8 $\mu$g/mL and 22.4/147.7 $\mu$g/mL for Treatment 1 and Treatment 2 respectively [15]. This results in $f_{\mathrm{pembro}} \approx 2.90 \times 10^{-7}$ $(\mathrm{g/cm^3})$/mg of pembrolizumab administered for all doses. To convert this into units of
$(\mathrm{molec/cm^3})$/mg, we note that the molecular mass of pembrolizumab is approximately 149,000 g/mol [16], which corresponds to $f_{\mathrm{pembro}} \approx 1.17 \times 10^{12}$ $(\mathrm{molec/cm^3})$/mg.

#### C.13 Estimates for PD-1/pembrolizumab Complex on Cells

##### C.13.1 Estimate for $\lambda_{Q_A}$

The dissociation rate of the PD-1/pembrolizumab complex was measured using biolayer interferometry to be 2.6 day$^{-1}$ in [37]. Thus, we take $\lambda_{Q_A} = 2.6$ day$^{-1}$.

##### C.13.2 Estimate for $\lambda_{P_D A_1}$

We estimate $\lambda_{P_D A_1}$ by fitting it to target engagement (TE) at trough for a triweekly regimen, specifically PD-1 receptor saturation by pembrolizumab, based on data from the 03TLC9 study [60]. For example, the TE of pembrolizumab on CD8+ T cells in the TS, which we denote $\mathrm{TE}^{\mathrm{T_8}}$ is mathematically defined as

$$\mathrm{TE}^{\mathrm{T_8}} = \frac{Q_A^{T_8}}{P_D^{T_8} + Q_A^{T_8} + Q^{T_8}} \times 100\%,$$

which is the percentage of all PD-1 receptors on CD8+ T cells at the TS that are bound to pembrolizumab. TE of pembrolizumab on other cells in the TS and TDLN are defined and notated similarly, with minimal deviation across all cell types. We define the overall TE as the average TE across all cell types in the TS and TDLN. The median TE at trough for a triweekly regimen at various doses is shown in Table C.1, noting that we assume a patient mass of 80 kg.

Table C.1: Median TE at trough for triweekly pembrolizumab regimens at various doses.

| **Dose (mg/kg)** | **Median TE** |
|---|---|
| 0.1 | 59 |
| 0.2 | 80 |
| 0.5 | 92 |
| 1 | 96 |
| 2 | 98 |
| 5 | 99 |

Noting that it takes approximately 19 weeks for a triweekly pembrolizumab regimen to reach steady-state concentrations [39], we consider 14 treatment cycles, corresponding to 294 days, to ensure that PD-1/pembrolizumab complex steady-state concentrations are achieved. We define $f(\lambda_{P_DA_1}, \xi_{\text{pembro}})$ as the overall TE at trough, in this case at 294 days, for a triweekly regimen with a dosage of $\xi_{\text{pembro}}$ mg/kg predicted by the model with a PD-1/pembrolizumab formation rate of $\lambda_{P_DA_1}$. To estimate the best value of $\lambda_{P_DA_1}$, we minimise the sum of squares of the differences at trough between $f(\lambda_{P_DA_1}, \xi_{\text{pembro}})$ and the true value based on the data from Table C.1. Thus, assuming pembrolizumab infusions every 3 weeks from $t = 0$days up until 294 days, we aim to minimise

$$\begin{aligned}\text{Objective} = &\left(f(\lambda_{P_DA_1}, 0.1) - 59\right)^2 + \left(f(\lambda_{P_DA_1}, 0.2) - 80\right)^2 + \left(f(\lambda_{P_DA_1}, 0.5) - 92\right)^2 \\ &+ \left(f(\lambda_{P_DA_1}, 1) - 96\right)^2 + \left(f(\lambda_{P_DA_1}, 2) - 98\right)^2 + \left(f(\lambda_{P_DA_1}, 5) - 99\right)^2.\end{aligned} \tag{C.7}$$

We perform a parameter sweep to minimise (C.7) and set the parameter space to be $\lambda_{P_DA_1} \in (0 \ \ (\text{molec/cm}^3)^{-1}\,\text{day}^{-1}, 10^{-12} \ \ (\text{molec/cm}^3)^{-1}\,\text{day}^{-1}]$. Solving this, the optimal value of $\lambda_{P_DA_1}$ was found to be

$$\lambda_{P_DA_1} = 4.63 \times 10^{-13} \ \left(\text{molec/cm}^3\right)^{-1} \text{day}^{-1}.$$
